# Comparing phenotypic manifolds with Kompot: Cluster-free differential expression at single-cell resolution

**DOI:** 10.1101/2025.06.03.657769

**Authors:** Dominik J. Otto, Erica Arriaga-Gomez, Elana Thieme, Ruijin Yang, Stanley C. Lee, Manu Setty

## Abstract

Multi-condition single-cell datasets are routinely compared by first discretizing cells into clusters or cell types, then contrasting the conditions cluster by cluster. This forces a trade-off between resolution and power. Further, aggregation discards changes that are subtle and distributed across the phenotypic manifold, or that reverse direction between related cell states. We introduce Kompot, a framework for cluster-free differential expression testing in multi-condition single-cell data. Kompot learns a continuous function of how gene expression varies across cell states in each condition using Gaussian process regression, then compares these functions to derive a per-cell log fold change for every gene. To assess significance, Kompot summarizes these fold changes with the Mahalanobis distance, which accounts for the statistical dependence between inferred expression values at related cell states, and calibrates it against an empirical null. Kompot therefore detects both the strong, broad changes and the subtle, coordinated changes that cancel under aggregation. We devised a semi-synthetic benchmark and demonstrate that Kompot outperforms existing methods in ranking precision across all five tested scenarios, particularly when expression changes are distributed across the manifold. Applied to murine hematopoietic aging, Kompot recovered age-related changes that are subtle in aggregate yet structured across cell states, including genes regulated in opposite directions across related cell types, without predefining which populations to test. In *Tal1^−/−^* mouse embryos, Kompot separated chimera-experiment artifacts from genuine signal and revealed that *Tal1* loss perturbs morphogenesis programs across many embryonic lineages beyond the previously described hematoendothelial compartment. In an immunotherapy cohort of advanced melanoma, Kompot showed that the greater efficacy of dual checkpoint blockade reflects an emergent, non-additive response that decouples T-cell activation from translation in cycling CD8 T cells, which neither monotherapy produces. Kompot scales near-linearly with the number of cells and is robust across parameter choices, and the same comparative framework extends to cell-state composition for differential abundance testing. Kompot therefore provides a unified, single-cell–resolution toolkit for comparative analysis of complex single-cell landscapes.

## Introduction

Multi-condition single-cell datasets offer unprecedented opportunities to dissect how mutations, treatments, and other perturbations reshape cellular states in differentiation and disease. Gene expression changes resulting from such perturbations are often subtle and distributed across the phenotypic manifold, differing between related cell states and sometimes reversing in direction across cell populations [1, 2]. Clustering is the primary approach for making such comparisons tractable and interpretable. Each cluster of cells is summarized by the mean expression of each gene in the condition, and differences between conditions are then tested cluster by cluster by comparing these means [3, 4]. This discretization forces a trade-off between resolution and power since broad clusters average over continuous changes (**Fig. 1A-B**), whereas finer clusters contain too few cells to reliably estimate means. Further, aggregation can discard subtle but biologically meaningful signals. For example, summarizing cells before testing can conflate genuine biological variation across cell states with technical variability (**Fig. 1A**), or eliminate structured changes that move in opposite directions across related cell states (**Fig. 1B**). Therefore, detecting these effects and distinguishing genuine state-dependent signal from technical noise requires quantifying them across the continuous phenotypic manifold at cell-state resolution.

**Figure 1:**
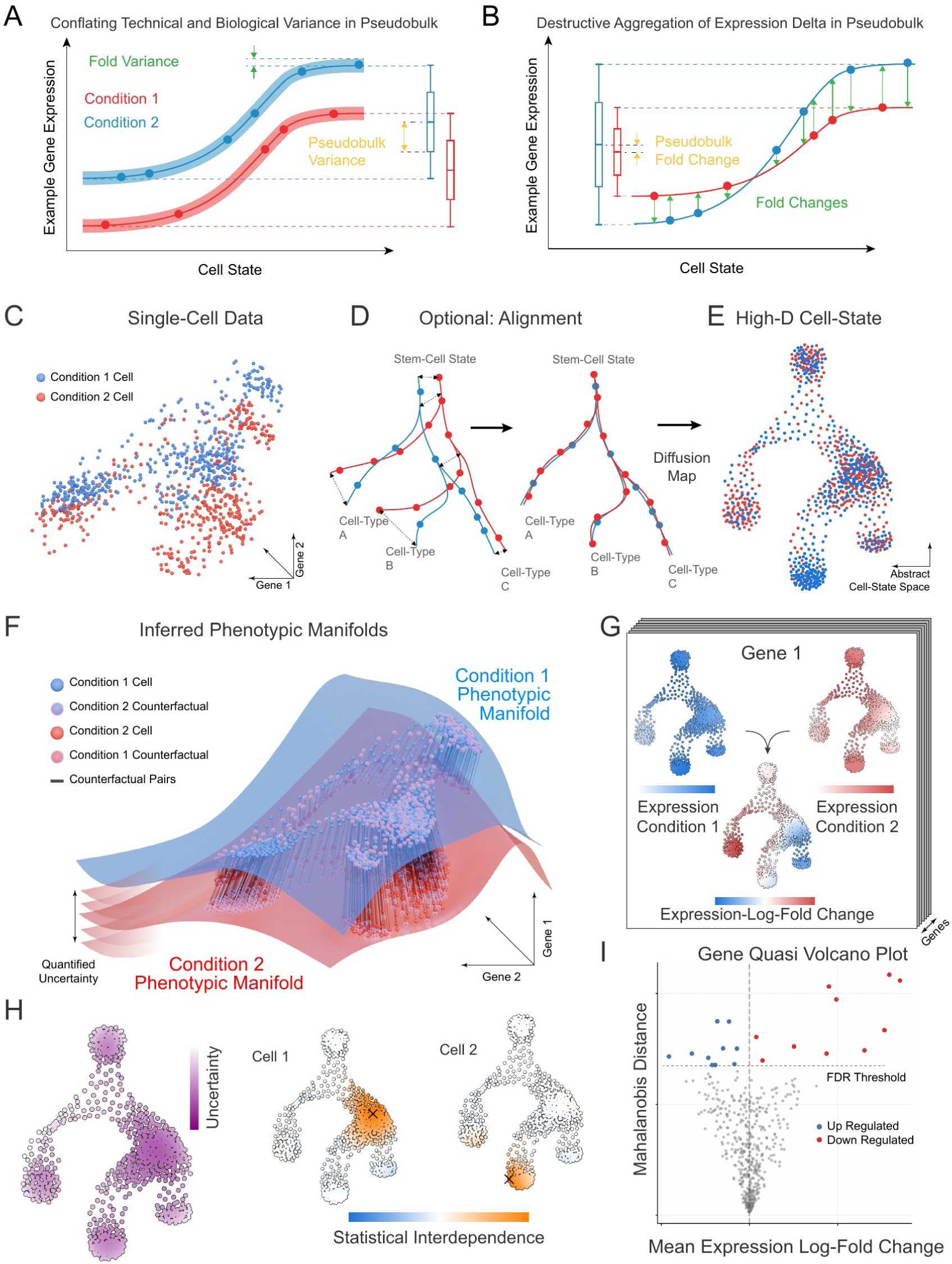
Overview of Kompot differential expression analysis framework. (**A**) Schematic of variance conflation: cells of each condition (red, blue) follow a smooth expression trajectory along the cell-state manifold, with biological structure within each condition that pseudobulk aggregation cannot separate from sample-to-sample technical variance. **(B)** Schematic of destructive aggregation: a gene whose per-cell differences are up in one region of the manifold and down in another, so that the pseudobulk log fold change averages to near zero. **(C)** High-dimensional single-cell transcriptomic data from two conditions in gene-expression space. **(D)** Optional alignment of comparable cell states between the two conditions (e.g., batch-effect correction), so that equivalent states co-localize before comparison. **(E)** Projection into a shared low-dimensional (*∼*50-dimensional) cell-state space using diffusion maps. **(F)** Condition-specific expression functions inferred over the cell-state manifold by Gaussian process regression, with counterfactual cell pairs (a cell and its equivalent state under the alternate condition) and the uncertainty of each mapping function. **(G)** Per-condition expression and fold change. For an example gene, the condition-specific expression levels are combined into a per-cell expression log fold change; stacking these across genes yields a gene *×* cell-state log fold change map. **(H)** Per-cell uncertainty of the inferred expression, and the covariance structure the Gaussian process induces: for each of two example cell states, one UMAP showing that state’s covariance with every other cell. **(I)** Gene-level quasi-volcano plot: average expression log fold change (*x*) against Mahalanobis distance (*y*).

Recent methods reduce the reliance on clustering, but they do not yet provide both cell-state resolution and calibrated significance. Neighborhood-based approaches such as miloDE [5] improve resolution by testing within local cell neighborhoods rather than predefined clusters, yet aggregating cells within each neighborhood still trades resolution against power. Continuous models like LEMUR [6] instead express change as a smooth function of cell state. However, these methods constrain this function to restricted classes such as a linear model in a low-dimensional embedding, which limits the expression patterns it can capture. Further, they do not provide a calibrated test of significance at cell-state resolution.

Here, we introduce Kompot, a framework for cluster-free differential expression testing in multi-condition single-cell data. Kompot first learns a continuous function of how gene expression varies across cell states in each condition using Gaussian Process Regression [7], a nonparametric class of functions that is not constrained to a predefined functional form. Kompot then compares these functions between conditions to derive a per-cell log fold change for every gene. To assess significance, Kompot uses Mahalanobis distance [8], which aggregates the per-cell-state fold changes across the phenotypic manifold into a single significance measure while accounting for the statistical dependence between the inferred expression values at related cell states. Mahalanobis distances are calibrated against an empirical null. As a result, Kompot can detect both strong, broad changes and the subtle, coordinated changes that occur in opposite directions across cell states. Further, the Kompot comparative framework can also be extended to cell-state composition for differential-abundance testing. Therefore, Kompot enables comprehensive and statistically rigorous comparisons of single-cell phenotypic landscapes.

We demonstrate the utility of Kompot by applying it to three diverse biological systems, uncovering novel biology in each. We identified that hematopoietic aging involves genes with opposing regulation across related cell types. Further, along the neutrophil lineage, we identified age-related changes that were confined to stem cells and did not persist as the cells differentiated. In *Tal1^−/−^* mouse embryos, Kompot disentangled experimental artifacts from biological signal and showed that loss of *Tal1* perturbs morphogenesis programs across many embryonic lineages in addition to the previously described hematoendothelial compartment. We then applied Kompot to a single-cell dataset of melanoma immunotherapy clinical trials and identified that the increased efficacy of dual checkpoint blockade likely reflects an emergent, non-additive response, that neither drug produces alone. Finally, we devised a semi-synthetic benchmarking framework and used it to compare Kompot against nine existing differential expression methods. Kompot outperforms these methods across a range of scenarios, with the largest gains where changes are distributed across the manifold. We also demonstrate that Kompot is robust to parameter choices and scales near-linearly with the number of cells and genes.

## Results

### Overview of the Kompot modeling framework

Kompot performs cluster-free differential gene expression analysis of multi-condition single-cell data. The central modeling assumption of Kompot is that gene expression varies smoothly along the underlying phenotypic manifold. Specifically we assume that cells in similar states share similar expression programs, and condition-induced transcriptional changes can be represented as smooth functions on the same manifold.

The input to Kompot is a latent representation of co-embedded multi-condition single-cell data (e.g., principal component analysis) (**Fig. 1C**). This latent representation can incorporate batch effect correction [9, 10] to ensure that cell states considered equivalent between conditions share similar locations in the latent space (**Fig. 1D**). Using this, Kompot employs diffusion maps [11] to derive a cell-state representation with biologically meaningful structure (**Fig. 1E**). Diffusion maps are particularly effective for single-cell analysis as they capture the intrinsic geometry of the data while providing interpretable measures of cell-to-cell distance [11–13]. Kompot can also operate on alternative low-dimensional embeddings, provided they preserve meaningful distances between cell states.

Kompot then constructs condition-specific mappings from cell-states to gene expression (**Fig. 1F**). These mappings are computed using Gaussian Processes (GPs) [7] to predict log-transformed gene expression as a continuous function of the diffusion map coordinates. GPs do not assume any functional forms and thus are particularly well-suited for this task as they can robustly capture the non-linear relationship between cell-states and gene expression [7]. Critically, GPs also provide uncertainty quantification for the mapping functions [7, 14], which we leverage for significance testing.

The continuous nature of these functions forms the fundamental basis for Kompot: A GP trained on cells from one condition defines an expression function over the entire manifold and can therefore be evaluated at all cell states observed in the other condition. This produces counterfactual expression profiles that reflect the expected gene expression in equivalent cell states under the alternate condition (**Fig. 1F, G**). By computing the difference between the expression in a cell and its counterfactual expression, Kompot quantifies gene expression log fold changes for every gene at single-cell resolution across the entire phenotypic landscape (**Fig. 1G**). Since this log fold change is defined separately for every cell, it can be summarized over any set of cells for interpretation.

Determining which genes show fold changes that are significant in a cluster-free manner requires accounting for both the uncertainty in the mapping function and the correlations between fold changes at nearby cell states (**Fig. 1H**). Therefore, instead of traditional p-values, which capture mapping uncertainty but not state interdependence, Kompot employs the Mahalanobis distance as the significance measure [8]. Mahalanobis distance effectively measures how many standard deviations a gene expression fold change is away from the expected variation, while accounting for the covariance structure between similar cell states. This enables Kompot to detect not only genes with significant changes in mean expression, but also genes with subtle yet consistent changes across related cell states, and genes with opposing changes across subpopulations that would cancel to near-zero fold change under any aggregation-based test (**Fig. 1A-B**). Kompot uses these distances as the score in a quasi-volcano plot (**Fig. 1I**) to identify significantly differentially expressed genes. Finally, to translate Mahalanobis distances into a significance threshold analogous to a p-value cutoff, we use a gene-shuffling procedure to build an empirical null distribution of distances (Methods).

### Benchmarking Kompot differential expression testing

To benchmark Kompot against existing approaches, we generated a novel scRNA-seq dataset of murine bone marrow at 10, 63, and 103 weeks of age (**Fig. 2A**, **Fig. 3A**). After quality control verification (**Supp. Fig. 1A-C**), we identified most hematopoietic cell types across all ages (**Fig. 2A**, **Fig. 3A, Supp. Fig. 1D–E**). Using the young mouse subset (2,917 cells, 16,285 genes), we constructed a semi-synthetic spike-in framework in which known gene expression changes are introduced across the manifold, enabling controlled evaluation of method performance. We constructed the following scenarios to span the range of expression-change patterns encountered in differential expression analysis, including settings that cluster-based methods are designed for and patterns that continuous methods are likely better suited for (**Fig. 2B**, **Supp. Fig. 2**):

- *Celltype*: gene expression change is confined to a particular cell type.
- *Celltype-var*: a uniform expression change applied to a coarse cell type whose finer subtypes differ in baseline expression, so the change must be recovered on top of real within-type variation. Since the change is uniform while the baseline varies, this biological variability presents as dispersion and can mask the change for cluster-based methods (**Fig. 1A**).
- *Multi-celltype*: expression changes in different directions across several cell types such that the global fold change cancels to near zero (**Fig. 1B**).
- *Gradient*: expression change along a lineage trajectory where no cluster boundary separates cells with and without expression change.
- *Uniform*: global expression change across the manifold.

**Figure 2:**
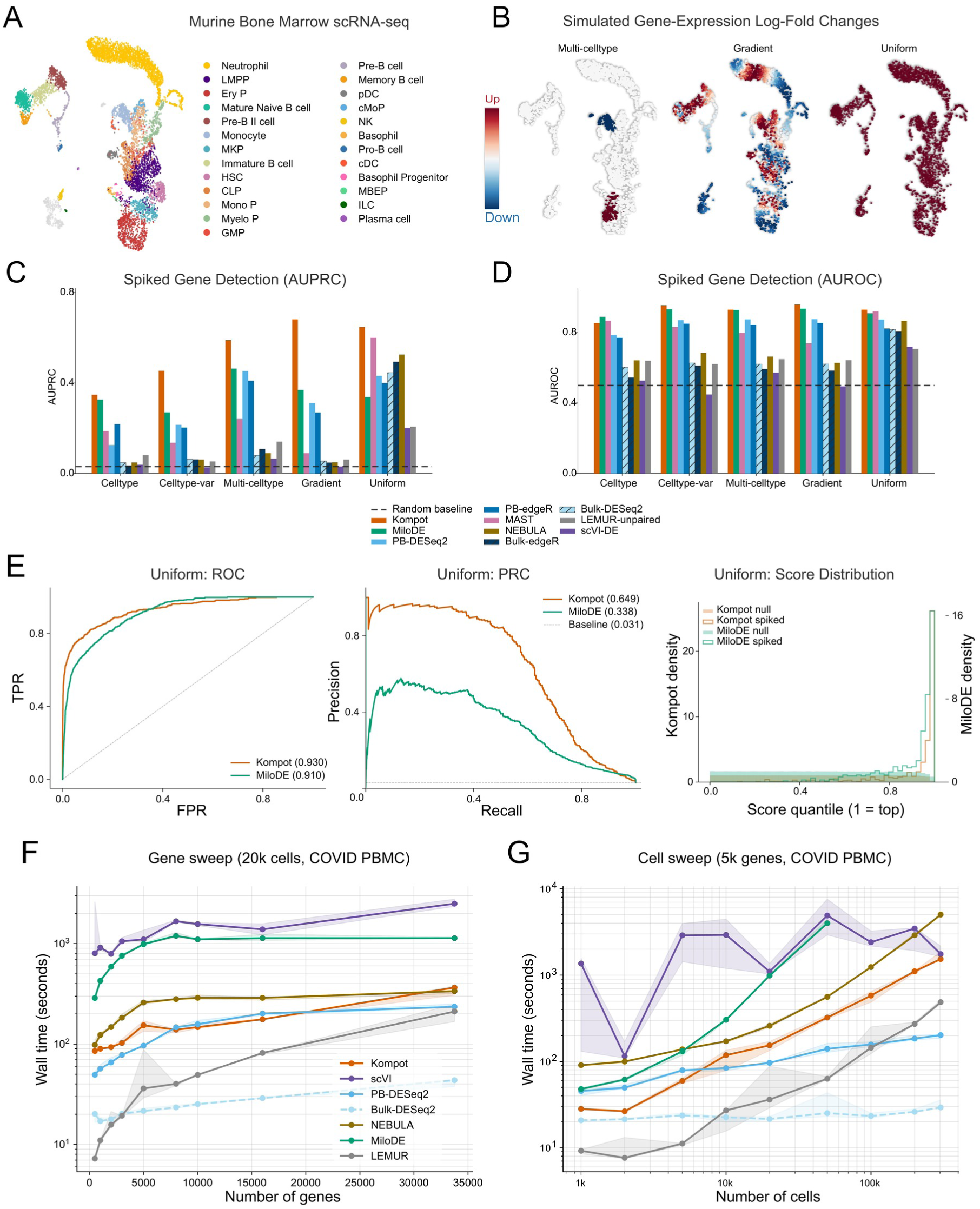
Semi-synthetic DE benchmark and runtime scaling. **(A)** UMAP of the murine bone marrow CITE-seq atlas (8,090 cells, 16,285 genes); the young subset (2,917 cells) is the base for the semi-synthetic spike-in. Cells are colored by manual cell-type annotation. **(B)** Representative per-cell ground-truth log fold change patterns for three of the spike-in scenarios (*Multi-celltype*, *Gradient* and *Uniform*) overlaid on the UMAP of (A). Each panel shows one representative gene per scenario; red/blue encode positive/negative LFC. Ground-truth patterns for all five scenarios are shown in **Supp. Fig. 2**. **(C)** Spiked-gene detection AUPRC across the five spike-in scenarios for Kompot, MiloDE [5], MAST, PB-DESeq2, PB-edgeR, NEBULA, Bulk-DESeq2, Bulk-edgeR, scVI-DE [15] and LEMUR-unpaired [6], grouped by scenario. Dashed line marks the chance baseline, 0.0307 (500 spiked of 16,285 tested genes). **(D)** Same as (C) for AUROC. Dashed line marks the random baseline (0.5). **(E)** Kompot versus MiloDE on the *Uniform* scenario. Left, ROC curves; middle, precision– recall curves, with the areas given in the legends and the chance baseline dashed; right, rank-quantile distributions of the null and spiked gene scores (1 = top). **(F)** Runtime scaling of representative DE methods as a function of the number of genes tested, holding the cell count fixed at 20,000 on the COVID-19 PBMC dataset [16]. Lines show median across 5 random seeds; shaded bands show the 25th–75th percentile. Methods: Kompot, scVI [15], PB-DESeq2, Bulk-DESeq2, NEBULA, MiloDE [5], LEMUR [6]. **(G)** Same as (F), sweeping the number of cells at a fixed 5,000 genes. MiloDE points are absent above *∼*50,000 cells, where the underlying MiloDE R implementation segfaults.

**Figure 3:**
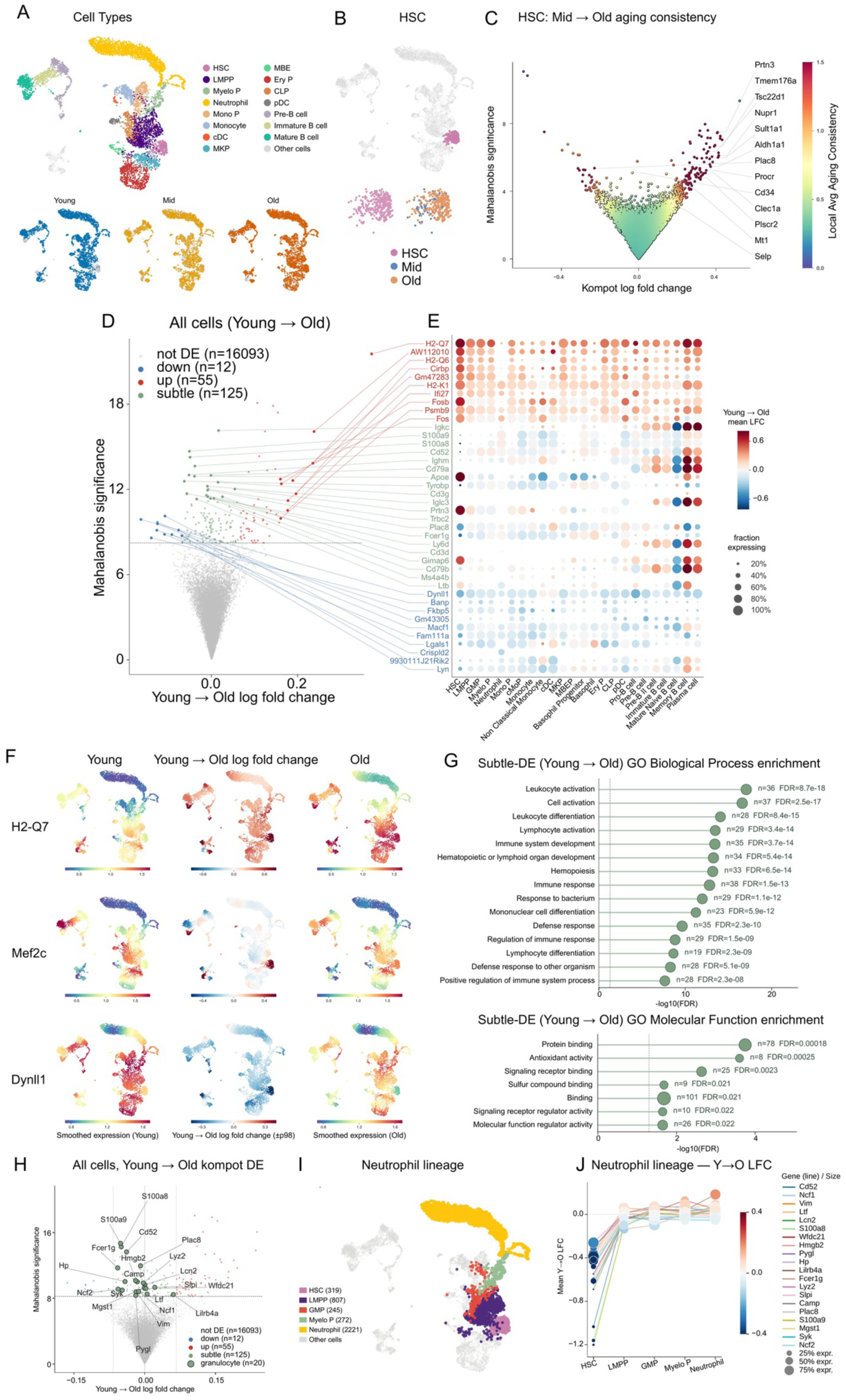
Kompot differential expression analysis in aging hematopoiesis. **(A)** UMAP of the murine aging bone marrow dataset colored by cell type. Bottom: Same embedding shown separately for Young, Mid and Old mice. **(B)** Hematopoietic stem cells (HSCs) highlighted on UMAP from (A), and the HSC subset alone colored by age group. **(C)** Mid-to-Old differential expression within HSCs. Quasi volcano plot with mean expression log fold change on the x-axis and Mahalanobis significance on the y-axis, colored by the local average aging-consistency score reported by the Hematopoietic Aging Atlas [29]. **(D)** Young-to-Old differential expression across all cells, on the volcano axes from (C). Genes identified as significant by Kompot’s local-FDR test are partitioned into up-regulated (*n* = 55), down-regulated (*n* = 12) and subtle (*n* = 125) classes. **(E)** Dot plot of Young-to-Old mean log fold change per cell type for genes selected from (D). Dot size encodes the fraction of cells expressing the gene. **(F)** Smoothed expression in young cells (left) and old cells (right), with the Young-to-Old log fold change between them (middle), for three example genes. **(G)** Gene-ontology enrichment of the subtle differentially expressed genes from (D), for GO Biological Process (top) and GO Molecular Function (bottom). **(H)** The Young-to-Old volcano of (D), highlighting the subtle differentially expressed genes associated with the neutrophil lineage. **(I)** UMAP of the aging bone marrow dataset with only the neutrophil lineage (HSC, LMPP, GMP, Myelo P, Neutrophil) colored. **(J)** Young-to-Old log fold change of the genes highlighted in (H) across the developmental stages of the neutrophil lineage (HSC, LMPP, GMP, Myelo P, Neutrophil). Dot size encodes the fraction of cells expressing the gene.

For each scenario, we synthetically spiked 500 of the 16,285 tested genes (3%) with a known per-cell log fold change pattern across eight simulated replicates per condition (**Supp. Fig. 2**). We scored each method on its ability to rank the spiked genes above the unperturbed background using AUPRC and AUROC (**Fig. 2C, D**). Since only 500 of the 16,285 genes are truly differentially expressed, AUPRC is the more informative metric for this imbalanced setting as it penalizes false-positive pile-up at the top of the ranking. We compared Kompot with MiloDE [5], NEBULA [17], MAST [18], scVI-DE [15] and LEMUR-unpaired [6], together with DESeq2 [19] and edgeR [20] applied in two ways: to pseudobulk profiles of unsupervised Leiden clusters [21], and to each sample as a whole in the manner of bulk RNA-seq. The full details of experimental design for benchmarking are described in **Supp. Note 4**.

Kompot is the most precise method in all five scenarios. Kompot AUPRC ranges from 0.348 to 0.681 across the five, and sits 0.02 to 0.31 above MiloDE [5], the strongest comparator overall (**Fig. 2C**). The margin is widest in *Gradient* (+0.31) and *Uniform* (+0.31), where the simulated gene expression change is spread over the manifold, and narrowest in *Celltype* (+0.02), where it is confined to one annotated population. MiloDE is the second best performing method across all scenarios except *Uniform* where MAST, NEBULA and bulk methods all perform better. We note that MiloDE returns no statistic for about a fifth of the genes, all of which are low-expression genes that were never eligible to be spiked. Our benchmarking framework credits these genes as correct rejections, while Kompot scores them and can rank one above a spiked gene. Restricting the comparison to the genes both methods score widens the margins (**Supp. Note 4**). It should also be noted that LEMUR-unpaired [6] is near chance across most scenarios since its statistical model requires a paired patient-by-condition design that our nested-replicate benchmark does not satisfy.

AUROC unlike AUPRC shows comparable performance across methods and scenarios although Kompot retains a marginal advantage over MiloDE in four of the five scenarios (**Fig. 2D**). The two metrics disagree because Kompot and MiloDE are accurate in different parts of the ranking (**Fig. 2E**). The AUROC gap is narrow because the two ROC curves converge at high false-positive rates (**Fig. 2E left**). At the low false-positive rates that are critical for interpretation, Kompot’s ROC curve lies above MiloDE’s (**Fig. 2E left**), and as a result the precision–recall gap is large (**Fig. 2E middle**). We then ranked genes by the differential expression statistics of each method (**Supp. Note 4**) and observed that ground-truth differentially expressed genes accumulate in the top quantiles for Kompot, whereas the MiloDE ranks overlap more with the null (**Fig. 2E right**). Therefore, Kompot’s advantage lies at the top of the ranked gene list, where genes that are important for downstream interpretation are chosen from.

Together, these results demonstrate the advantage of Kompot over competing methods for differential expression testing across both cluster-dependent and cluster-free scenarios.

### Scalable and robust differential expression testing

We next tested the scalability of Kompot using the large-scale COVID-19 peripheral mononuclear blood cells (PBMCs) dataset [16]. We restricted this dataset to healthy individuals and COVID-19 patients and balanced the two conditions, yielding a dataset of 302,624 cells from 60 samples (35 healthy individuals and 25 COVID-19 patients) that we used for this benchmark. Kompot runtime depends only weakly on the number of genes (**Fig. 2F**) and scales near-linearly with the number of cells (**Fig. 2G**), with memory consumption following the same trend (**Supp. Fig. 3**). MiloDE [5] segfaults above *∼*50,000 cells in this benchmark, and several generative-model and pseudobulk methods plateau at runtimes one to two orders of magnitude above Kompot.

The dominant cost driver for Kompot is the number of landmarks (default 5,000), which sets the sparse-GP rank and leads to a quadratic runtime contribution (**Supp. Fig. 3, 4**). We therefore tested the robustness of Kompot results to the number of landmarks and other parameters (**Supp. Figs. 5–8**). We varied each parameter over a broad range of values and compared the results to baseline using Pearson correlation. We performed all our robustness checks using our murine aging hematopoiesis dataset as representative of a dataset with continuous trajectories and comparison of healthy donors and COVID-19 patients [16] for changes in expression changes of PBMCs as representative of a dataset with discrete clusters.

Number of cells in a dataset is often a technical variable with tissue handling and costs being the predominant factor in the choice of dataset size rather than any representation of underlying biology. We therefore down sampled cells in both datasets and determined the correlation of Kompot differential expression log fold change or Mahalanobis distance between down sampled and the full dataset (**Supp. Figs. 5-8A**). Kompot fold changes and Mahalanobis distances are robust even when the number of cells is reduced by an order of magnitude. We next varied the number of diffusion components which is a user guided parameter that accounts for the biological complexity of the dataset (**Supp. Figs. 5-8D**). Kompot differential expression results are robust to large changes in the number of components demonstrating the results are not sensitive to the choice of number of diffusion components. Further, we tested robustness to the parameters of the mapping function: length scale factor which controls the extent to which to share information between neighboring states; number of landmarks, which is used for sparse Gaussian Processes for efficiency and sigma, which encodes the expected noise in the manifold to gene expression mapping function. Differential expression results are robust to large changes in all of these parameters suggesting that default parameters provide reliable, interpretable results (**Supp. Figs. 5-8B-C, E**). These results collectively highlight the robustness of Kompot differential expression results across continuous and discrete datasets.

### Kompot resolves subtle and heterogeneous aging changes across the hematopoietic landscape

Aging induces progressive, often subtle changes in cellular composition and gene expression rather than dramatic phenotypic transformations [22]. These changes need not be uniform across cell types [23], making aging a compelling context for evaluating differential expression methods, and one where a cluster-free approach is likely to aid interpretation. Hematopoiesis has emerged as a particularly informative system for studying aging [24], characterized by declining stem cell function [25], reduced lymphocyte production [26, 27], and increased susceptibility to myeloid malignancies [28]. To investigate these age-related changes, we applied Kompot to our murine bone marrow aging dataset (**Fig. 2A**, **Fig. 3A**).

We first applied Kompot to characterize gene expression changes in hematopoietic stem cells (HSCs) and monocytes, two cell types where transcriptional changes with age have been well characterized [30, 31] (**Fig. 3B–C, Supp. Fig. 9**). Aging in HSCs has been extensively studied, and the Hematopoietic Aging Atlas has synthesized results from multiple datasets to identify genes consistently up– or down-regulated with age [29]. For each gene, this atlas assigns a consistency score that reflects the reproducibility of age-associated expression changes across studies [29]. Kompot results comparing mid to old HSCs strongly prioritized genes with the highest consistency scores in the Hematopoietic Aging Atlas, with genes showing large fold changes and Mahalanobis distances highly enriched for reproducible aging signatures (**Fig. 3C, Supp. Fig. 10**). This included up-regulation of the cell adhesion molecule Selp, the most consistently deregulated gene in aging HSCs [29], which has been functionally linked to impaired long-term hematopoietic reconstitution [32]. Monocytes, similar to HSCs, exhibited clear transcriptional changes from young to old mice (**Supp. Fig. 9A–B**). Gene ontology analysis revealed that downregulated genes were enriched in core monocyte/macrophage functions such as defense response (**Supp. Fig. 9D**), consistent with their reduced functional capacity with age [31]. In contrast, upregulated genes were enriched for antigen processing and presentation pathways, driven by increased expression of MHC class II genes (**Supp. Fig. 9C**), which have been implicated in age-related immune remodeling [33]. Notably, our results recapitulate findings from a previous study that identified H2-Aa, H2-Ab1, H2-Eb1, Cd74, and Aw112010 as the top dysregulated genes in aging monocytes across both mice and humans, with only Igkc surpassing H2-Eb1 in our ranking [33]. Together, these results demonstrate that Kompot recovers established aging signatures in both HSCs and monocytes.

We next applied Kompot across the entire hematopoietic landscape comparing young and old mice to determine whether our cluster-free approach can provide novel insights into murine hematopoietic aging (**Fig. 3D–F**). Kompot identified genes spanning the full range from up-to down-regulation with age, including genes with subtle aggregate fold change that were nonetheless significant by Mahalanobis distance (**Fig. 3D–F**). Gene ontology analysis of upregulated genes (high positive fold change and high Mahalanobis distance) showed significant enrichment for antigen presentation and MHC class II genes. These results are consistent with the monocyte enrichments and indicate that MHC class II upregulation is a general feature of hematopoietic aging across all cell types. Similarly, genes with high Mahalanobis distance and strong negative fold change were consistently downregulated across multiple cell types (**Fig. 3E**). In contrast, genes ranked highly by Mahalanobis distance but with subtle fold change exhibited divergent expression patterns across cell types, with upregulation in some and downregulation in others (**Fig. 3E**).

We next performed gene ontology analysis and found strong enrichments for terms associated with hematopoietic differentiation and immune response (**Fig. 3G top**). This indicates that these genes are functionally relevant to hematopoietic aging and that biologically important age-related changes need not produce large, consistent fold changes across cell types. Further, enrichment using the molecular function terms revealed significant enrichment for antioxidant activity (**Fig. 3G bottom**), consistent with oxidative stress being an important factor in hematopoietic aging [34]. Interestingly, the fold change from young to old mice for the genes that led to this enrichment was cell-type dependent rather than uniform (**Supp. Fig. 11E**).

To further understand how genes with only a subtle aggregate fold change could still be relevant to aging, we examined their expression changes across cell types (**Supp. Fig. 11A**) and identified that the subtle aggregate signal arose from two distinct patterns. First, a number of genes involved in B cell differentiation showed opposing changes between naive and memory B cells that largely cancel in aggregate. Second, expression changes of many genes were confined to HSCs such that the aggregate change across the full population of cells remained subtle. We therefore grouped the genes by their HSC fold change into HSChi, HSCneutral, and HSClo sets (**Supp. Fig. 11B**). HSChi genes were enriched for hemopoiesis and its regulation, as expected for genes associated with HSC quiescence (**Supp. Fig. 11C**, [35, 36]), whereas HSClo genes were enriched for processes related to differentiated granulocyte function (**Supp. Fig. 11D**). We therefore asked whether the aging changes in these granulocyte-associated genes are retained as HSCs differentiate (**Fig. 3H–J**). We examined the expression of these genes along the granulocyte representative neutrophil lineage and identified that expression levels in older mice returned to near those in younger mice immediately as cells exited the HSC state and entered differentiation, rather than gradually across the lineage (**Fig. 3J**). This suggests that the aging signature in granulocyte-associated genes is largely confined to HSCs and is reset upon lineage commitment, potentially buffering mature granulocytes from HSC-intrinsic aging.

These results demonstrate that, by quantifying expression differences directly on the manifold without prior clustering, Kompot captures biologically meaningful, context-specific changes in gene expression that are subtle in aggregate yet coherent across related cell states.

### Kompot recovers *Tal1*-specific changes after correcting chimera artifacts in mouse embryos

We next applied Kompot to investigate gene expression changes during mouse embryonic development at midgestation (E8.5), using a single-cell RNA-seq dataset of wild-type and *Tal1*-knockout embryos [37] (**Fig. 4A**). *Tal1* is required for the development of primitive erythroid, megakaryocyte, and myeloid lineages [38, 39], but its effect on other lineages at mid-gestation is not as well characterized. We therefore used this dataset to characterize the effect of *Tal1* knockout across embryonic lineages.

**Figure 4:**
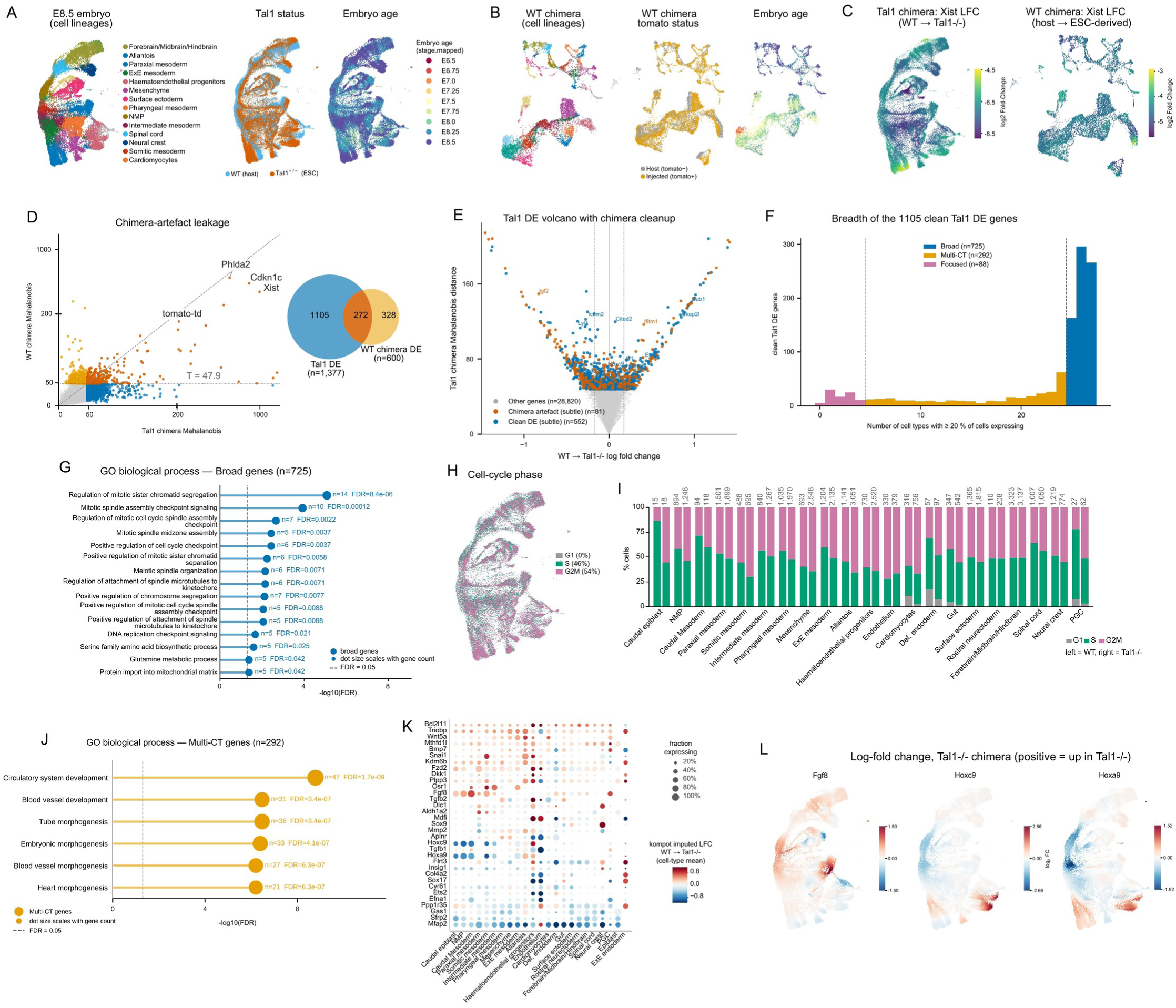
Kompot corrects chimera-experiment artefacts in *Tal1^−/−^* mouse embryos and identifies *Tal1*-specific changes in embryonic development. (A) UMAP of E8.5 mouse embryo cells [37] for the *Tal1^−/−^* chimera, colored by cell type, by *Tal1* status (host versus tdTomato^+^ ESC-derived) and by embryo stage. **(B)** The same three UMAPs for the WT-only control chimera (tdTomato^+^ wild-type ESC-derived cells injected into a wild-type host), colored by cell type, by chimera status (host versus injected) and by embryo stage. **(C)** Kompot per-cell log_2_ fold change for *Xist* on the UMAP, tdTomato^+^ ESC-derived relative to host: *Tal1^−/−^* chimera (left; host *n* = 16,762, ESC-derived *n* = 26,309) and WT-only chimera (right; *n* = 4,518 and 7,874). **(D)** Artefact cleanup: per-gene Mahalanobis distance in the *Tal1* chimera (*x*) versus the WT-only chimera (*y*). Genes ranked high in both are excluded at the matched-Mahalanobis cutoff (*T* = min Mahalanobis over *Tal1* local-FDR *<* 0.05). Color partitions the 633 *Tal1* subtle-DE genes into clean DE (*n* = 552) and chimera artefact (*n* = 81). **(E)** Quasi Volcano of the clean *Tal1* DE call: mean log fold change WT *→ Tal1^−/−^* (x) versus Mahalanobis distance (y), with representative genes labelled. **(F)** Clean DE genes split by how many cell types they are altered in — broadly altered (*≥* 25*/*27), multi-cell-type (5–24*/*27) and focused (*≤* 4*/*27). Of 1,105 clean DE genes, 725, 292 and 88 respectively. **(G)** GO Biological Process enrichment of the broadly altered set (725 genes; 312 terms at FDR *<* 0.05 against the 21,541-gene tested universe). **(H)** UMAP from (A) colored by the same per-cell phase. **(I)** Cell-cycle phase composition per cell type: one stacked bar pair per cell type, left wild-type, right *Tal1^−/−^*, normalized within each cell type *×* genotype and annotated with *n*. The 21 cell types with *≥* 10 cells in both conditions (43,025 of 43,071 cells) are shown. **(J)** GO Biological Process enrichment of the multi-cell-type set (292 genes). **(K)** Per-cell-type log fold change dotplot for the multi-cell-type genes intersected with GO:0048598 (embryonic morphogenesis); dot size encodes the fraction of cells expressing the gene and color represents the mean log fold change. **(L)** Per-cell Kompot log fold change (WT *→ Tal1^−/−^*; positive = up in *Tal1^−/−^*) on the UMAP for *Fgf8*, *Hoxc9* and *Hoxa9*, three of the multi-cell-type morphogenesis genes in (K).

Comparison of *Tal1* knockout and wild-type cells is however confounded by the experimental design. *Tal1* knockout was achieved by injecting tdTomato^+^ ESC-derived *Tal1^−/−^* cells into a wild-type host blastocyst. The donor ESC line is male while the host cells are largely female (72%), so mutant and host cells are separated by sex as well as by genotype (**Fig. 4C**). As a result, differences between mutant and host cells reflect not only *Tal1* loss but also sex-linked and imprinted genes. These differences are likely to be systematic across lineages since they arise from the chimera design.

To correct for these artifacts, we used a control chimera from the same dataset, generated identically but with wild-type (WT) donor cells (**Fig. 4B**). This chimera thus carries the same artifacts without *Tal1* loss. We applied Kompot to compare *Tal1^−/−^* and wild-type cells, after excluding the lineages depleted by *Tal1* loss since they are almost entirely wild-type in the chimera (**Supp. Fig. 12A**). As expected, the top-ranked genes were dominated by chimera artifacts, including ambient hemoglobin RNA, sex-linked genes, and imprinted genes (**Supp. Fig. 12B**). Applying Kompot to the WT chimera produced a similar set of top-ranked genes (**Supp. Fig. 12C**). Therefore, using the WT chimera as the matched null, we identified a set of genes with expression changes that could be attributed specifically to *Tal1* loss by excluding genes that were ranked high in both WT and *Tal1^−/−^* chimeras (**Fig. 4D**). This correction was made possible by the design of Kompot, which summarizes the expression change of each gene across the manifold as a single Mahalanobis distance and thus enables comparisons between the two chimeras. The two chimeras shared a large fraction of their top-ranked genes, showing that these artifacts are detected reproducibly and excluding them yields a smaller set of changes attributable to *Tal1* loss.

Similar to hematopoietic aging, *Tal1*-induced genes showed a spectrum of changes from down-to up-regulation, including a substantial number with low fold change but high significance (**Fig. 4E**). Among these genes, those with the highest fold changes tended to be broadly altered across cell types, whereas more subtle changes tended to be restricted to fewer cell types (**Supp. Fig. 12G**). To dissect these effects, we classified the genes by how broadly they are expressed across cell types: broadly expressed (*≥* 25*/*27 cell types), focused (*≤* 4*/*27), or multi-cell-type (the remainder) (**Fig. 4F**).

Gene ontology analysis of the broadly expressed genes revealed enrichment for cell-cycle terms, suggesting that *Tal1* loss alters cell-cycle dynamics across embryonic lineages (**Fig. 4G**). To examine this, we assigned cells to cell-cycle phases and observed a systematic shift toward G2M at the expense of S phase in the *Tal1^−/−^* chimera (**Fig. 4H–I**). Stage-matched wild-type cells, in contrast, did not show the same systematic shift across all cell types, suggesting that this imbalance is in part a consequence of Tal1 loss (**Supp. Fig. 12D**).

Changes in the focused set were largely confined to hematoendothelial progenitors and endothelial cells, consistent with the canonical role of *Tal1* in these lineages [40] (**Supp. Fig. 12F**). Notably, within these cells, many known *Tal1* targets changed in both direction and magnitude across cells, so that their aggregate fold change remained low (**Supp. Fig. 12E**). The multi-cell-type set, in turn, was enriched for morphogenesis programs, including blood vessel development (**Fig. 4J**). These genes were dysregulated across multiple cell types (**Fig. 4K**), indicating that the effects of *Tal1* loss on morphogenesis extend across many embryonic lineages rather than being confined only to hematoendothelial cells. Further, Fgf8 and Hox genes known to play a critical role in emergence, organization and differentiation of the mesodermal lineages [41, 42] were among these dysregulated genes reinforcing the broader role of *Tal1* loss during embryonic development (**Fig. 4K–L**).

These analyses demonstrate that testing for differential expression directly on the manifold allowed Kompot both to correct systematic experimental artifacts and to resolve *Tal1*’s effects across the developing embryo without the need to cluster cells beforehand.

### Kompot reveals determinants of immunotherapy in advanced melanoma

We next tested Kompot in a disease context, applying it to investigate how immune checkpoint blockade reshapes CD8 T-cell states in advanced melanoma. Combination checkpoint blockade with anti-LAG-3 (Relatlimab, Rela) and anti-PD-1 (Nivolumab, Nivo) provides greater clinical efficacy than Nivo alone [43]. Cillo et al. [43] profiled CD8 T cells from patients in a clinical trial of these therapies using single-cell RNA-seq (**Fig. 5A, B**) and proposed that combination treatment promotes an altered differentiation trajectory with increased cytotoxicity and exhaustion. CD8 T-cell activation and exhaustion span a continuous spectrum [2], making this a natural setting for Kompot’s cluster-free quantification of expression differences. We therefore reanalyzed baseline and ontreatment samples within each therapy arm to determine whether Kompot could recover complementary insights to those reported previously.

**Figure 5:**
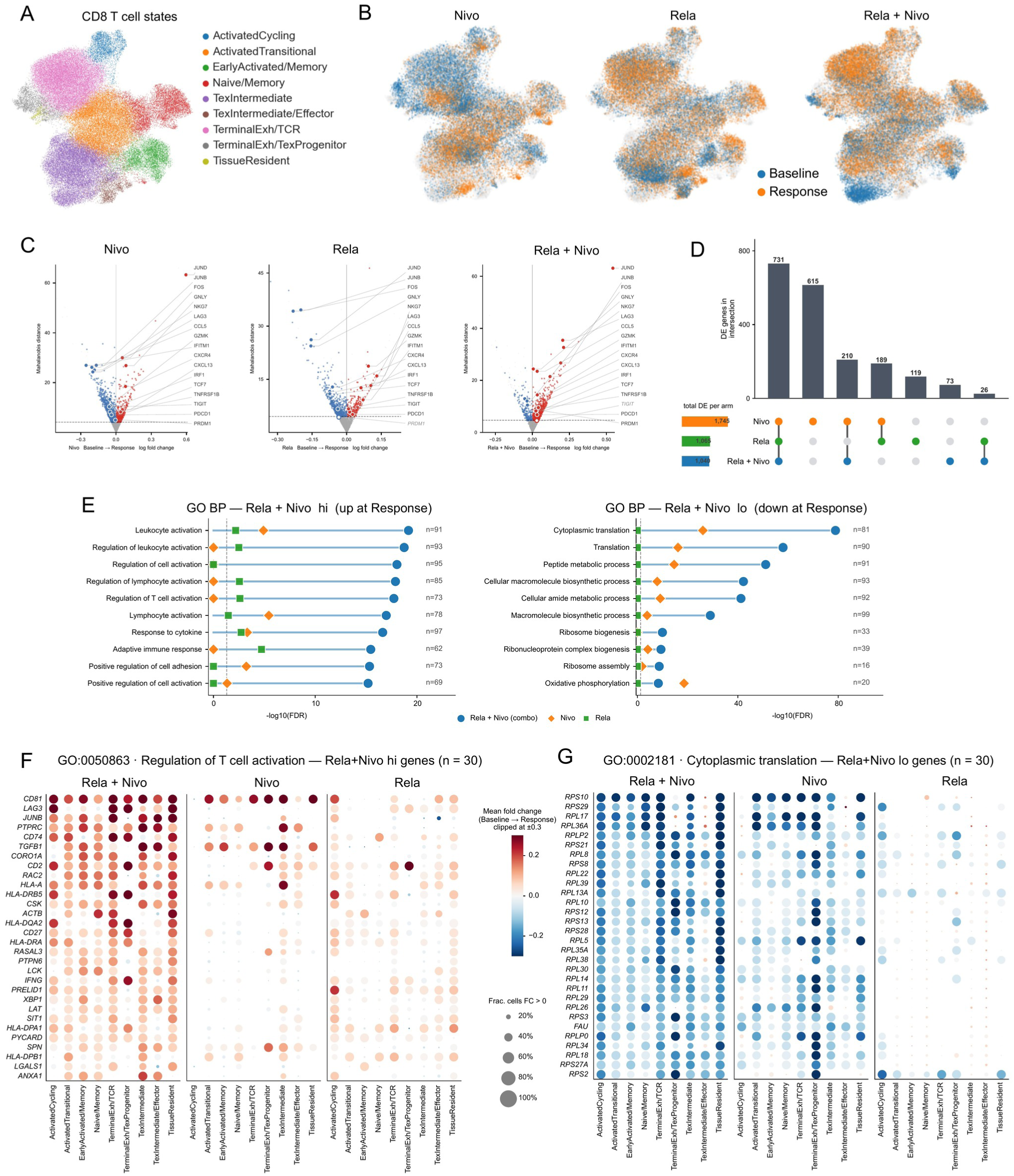
Kompot reveals determinants of immunotherapy in advanced melanoma. **(A)** UMAP of the CD8 T-cell landscape [43], colored by CD8 T-cell state annotation. **(B)** UMAP in (A) shown per treatment arm. Cells are colored by sample timepoint (Baseline vs. Response): Nivo: Nivolumab, anti-PD-1, Rela: Relatlimab, anti-LAG-3, and the Rela + Nivo combination. **(C)** Per-arm Kompot quasi-volcanoes (Mahalanobis distance, *y*; mean log fold change Baseline *→* Response, *x*) for Nivo (1,745 DE genes), Rela (1,065), and Rela+Nivo (1,040) at the calibrated local-FDR cutoff. **(D)** Overlap of the three per-arm differentially expressed gene sets. **(E)** GO Biological Process enrichments for the combination up-regulated (combo-hi, 733 genes) and down-regulated (combo-lo, 307 genes) gene sets, with the corresponding Nivo and Rela enrichments overlaid. **(F)** Per-cell-type log fold change dotplot for GO:0050863 “regulation of T-cell activation” across the three arms. Top 30 combination out of 73 up-regulated genes of that term with the largest increase in the combination arm are shown. **(G)** Per-cell-type log fold change dotplot for GO:0002181 “cytoplasmic translation” across the three arms. Top 30 combination out of 81 down-regulated genes of that term with the largest decrease in the combination arm are shown.

Kompot identified a large per-arm differentially regulated gene expression program (**Fig. 5C**). Interestingly, differential genes in the combination arm largely overlapped with those of the two monotherapies (**Fig. 5D**), indicating that its greater clinical efficacy does not arise from a combination-specific gene expression program. Gene ontology analysis of the combination genes however revealed striking differences (**Fig. 5E**). Upregulated genes in the combination arm were enriched for terms related to T cell activation and function, whereas upregulated genes in either monotherapy failed to achieve the same level of significance (**Fig. 5E left**). The downregulated genes in the combination arm were enriched for translation and related metabolic processes, again more significantly than in either monotherapy (**Fig. 5E right**).

To understand how different CD8 T cell states respond to combination treatment, we compared the fold change patterns of genes involved in T cell activation and cytoplasmic translation, as representatives of upregulation and downregulation respectively (**Fig. 5F–G**). In the combination arm, the T cell activation program was induced across all CD8 T cell states, including exhausted states, as observed in the original study [43] (**Fig. 5F left**). Combination treatment also drove a substantial reduction in the expression of cytoplasmic translation and related metabolic genes, particularly within the activated cycling CD8 T cell compartment (**Fig. 5G left**, **Supp. Fig. 13**). Proliferative expansion of effector CD8 T cells is normally coupled to a coordinated upregulation of the translation machinery [44]. The downregulation of translation-machinery genes within the same compartment where these genes are expected to be highly expressed indicates a decoupling of the translational program from the proliferative state. This suggests that combination treatment retasks the biosynthetic program away from ribosome gene expression and toward an effector program.

While Rela treatment led to a marginal upregulation of T cell activation genes in the cycling T cells (**Fig. 5F right**), the translation related genes remained broadly unchanged (**Fig. 5G right**). Nivo treatment on the other hand, did not substantially upregulate activation genes yet still downregulated translation genes (**Fig. 5F– G center**). Thus, within the activated cycling compartment, the combination treatment alone decoupled the T cell activation program from the translation program, through the same genes the monotherapies regulate. This therefore likely represents an emergent, non-additive consequence of dual blockade and a possible molecular basis for the combination’s greater clinical efficacy.

These results show that what separates the combination from either monotherapy is not which genes change but how those changes are distributed across CD8 T-cell states. The three arms draw on largely the same gene sets, and the distinction appears only once the fold changes are resolved by cell state: the combination induces activation across the compartment while suppressing translation in the cycling cells that typically upregulate it.

### Kompot resolves compositional changes in hematopoietic aging

The single-cell, cluster-free comparative framework used for differential expression testing in Kompot can be naturally extended to differential abundance testing, which identifies compositional changes in multi-condition single-cell data [45, 46]. Specifically, rather than comparing condition-specific expression functions over cell states, we compare condition-specific compositional functions between conditions. We previously developed Mellon [14] to estimate cell-state density functions, which describe how cells are distributed across the cell-state space and thus provide the compositional function needed for differential abundance testing. As in differential expression testing, we first compute a co-embedded cell-state representation of the two conditions, with batch correction as needed, and then estimate a condition-specific density function over this shared representation (**Fig. 6A**). The continuous nature of these estimates allows them to be evaluated at any cell state, including those observed predominantly in the alternate condition (**Supp. Fig. 14**). The ratio of the two density estimates at each cell state then gives a measure of differential abundance between conditions at single-cell resolution (**Fig. 6A**). We further extended Mellon to estimate the uncertainty in each density function, from which we derive a per-cell posterior tail probability (PTP) as a measure of statistical significance (Methods). We can then use effect size and PTP as the axes of a differential-abundance volcano plot to identify cell states with significant compositional changes (**Fig. 6A**).

**Figure 6:**
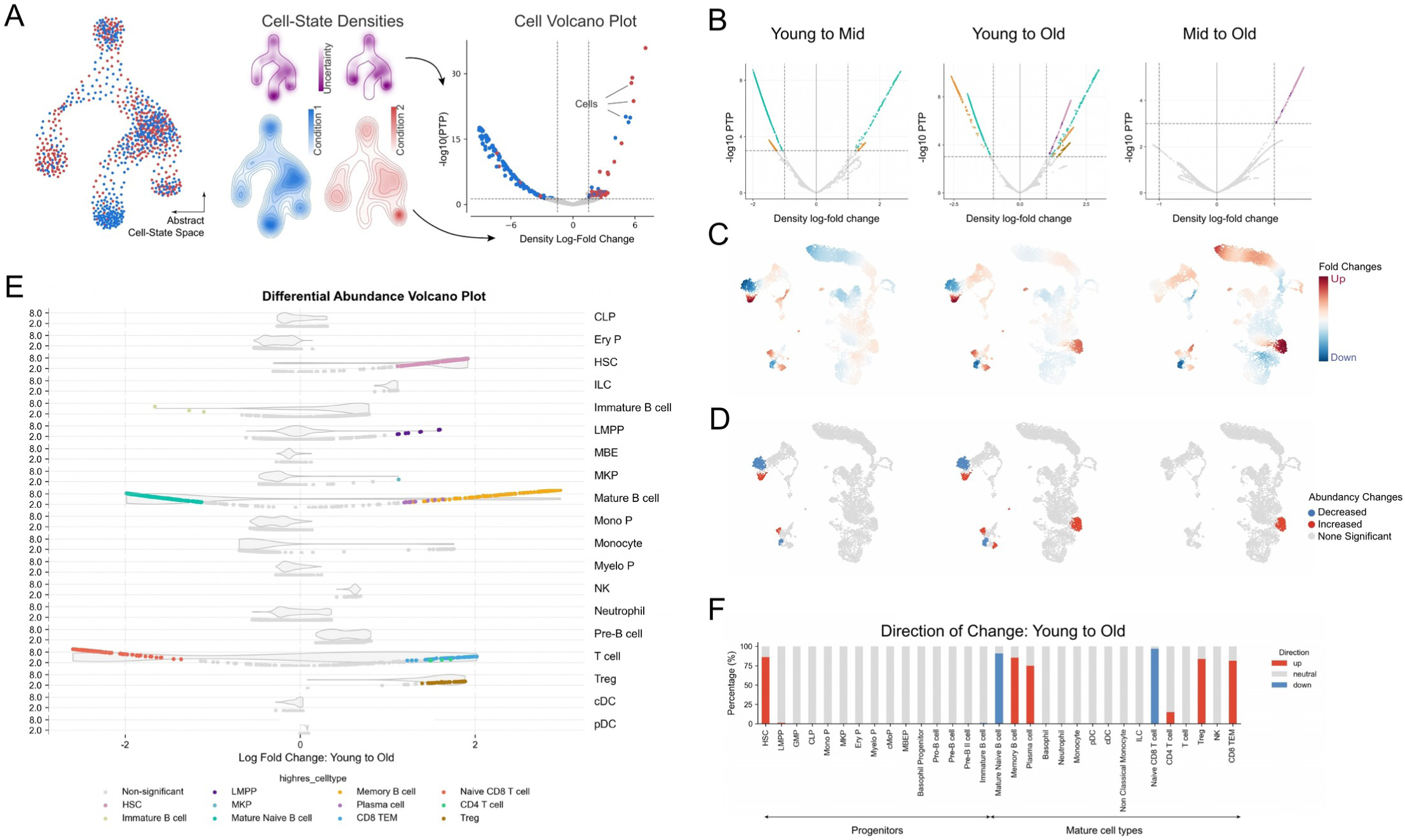
Differential abundance analysis of aging hematopoiesis. **(A)** Schematic of the Kompot differential-abundance test. Middle: Mellon [14] is used to estimate a cell-state density and its posterior variance per condition. Right: Log ratio of the two densities gives a per-cell density log fold change and the posterior gives a per-cell posterior tail probability (PTP). **(B)** Differential abundance results for the Young to Mid (left), Young to Old (middle) and Mid to Old (right) comparisons for the aging hematopoiesis dataset from **Fig. 3A**. Quasi-cell volcano plot with density log fold change on the x-axis and Posterior Tail Probabilities (PTP) on the y-axis. Significantly different cell-states (absolute log fold change *>* 1, PTP *<* 0.05) are highlighted by their cell type annotations. **(C)** UMAPs from **Fig. 3A** displaying the estimated local density log fold changes across the full joint cell-state space derived from all samples (Young, Mid, and Old) for comparisons in (B). **(D)** UMAPs highlighting the subset of cell states with statistically significant abundance changes, as defined by the thresholds shown in the volcano plots in (B). **(E)** Differential abundance volcano plots showing density log fold changes across cells, stratified by major cell types and colored by subtype in the Young to Old comparison. y-axis indicates posterior tail probability. Violin plots in the background indicate abundance of cells at the respective density log fold change in this group. **(F)** Direction and magnitude of abundance changes per cell type in the Young to Old comparison. Bars show the fraction of significantly changing cell-states within each annotated cell type, color-coded by the direction of change. Cell types are ordered along a progenitor-to-mature axis.

We first benchmarked the performance of Kompot differential abundance relative to existing approaches. Recent benchmarking studies have highlighted Meld [46] and Milo [45] as amongst the top-performing and most widely used differential abundance analysis approaches [47]. We therefore compared Kompot’s differential abundance analysis with Meld and Milo using the simulated (linear, cluster, branch) and COVID-19 datasets [48] described in the benchmarkDA study [47]. Kompot performance on both AUROC and AUPRC was comparable to both Meld and Milo, with Kompot showing greater robustness to batch effects (**Supp. Fig. 15**). While Meld was the top-performing method in these datasets with no introduced batch effects (**Supp. Fig. 15**), differential abundance analysis with Meld does not provide a measure of statistical significance. Our approach thus combines the individual advantages of Meld and Milo, providing single-cell resolution differential abundance with confidence measures. We next performed extensive robustness checks, as for differential expression, and determined that differential abundance testing is robust to the number of cells in a dataset (**Supp. Fig. 16A, 17A, 18A, 19A**), the number of diffusion components (**Supp. Fig. 16D, 17D, 18D, 19D**), and the parameters of Mellon density estimation (**Supp. Fig. 16B-C, 17B-C, 18B-C, 19B-C**).

We next applied differential abundance testing to our murine hematopoietic aging dataset (**Fig. 3A**) to determine age-dependent composition changes, performing pairwise comparisons between young, middle-aged, and old mice. Early aging (young *→* middle-aged) was dominated by remodeling of the mature compartment (**Fig. 6B–D**, left; **Supp. Fig. 20A**). We observed a marked shift from naïve to memory cells in both the B– and T-cell compartments, with no significant abundance changes in the progenitor compartment. The depletion of naïve B-cells and enrichment of memory B-cells with age is a well-characterized phenomenon of hematopoietic aging in both mice and humans [27, 49]. This reflects fundamental changes in differentiation dynamics: naïve B-cells rely on continuous influx from hematopoietic progenitors, whereas memory B-cells accumulate over time through antigen exposure [26, 49, 50]. The T-cell compartment showed the same naïve *→* memory transition with age, a hallmark of immunosenescence that contributes to diminished adaptive immune responses in aging [51]. Late aging (middle-aged *→* old), in contrast, shifted the abundance changes to the progenitor compartment, with no significant changes in the mature compartment (**Fig. 6B–D**, right; **Supp. Fig. 20B**). The most significant change was an accumulation of hematopoietic stem cells (HSCs) in older mice (**Fig. 6E–F**). Progressive enrichment of HSCs with age is well documented [30], and our analysis places this expansion specifically in the middle-to-old window rather than across the full lifespan. Finally, the full-lifespan young *→* old comparison (**Fig. 6B–D**, middle) aggregated both phases and recovered every cell type significant in either window, including the naïve *→* memory shifts in B and T cells from early aging and the HSC expansion from late aging (**Fig. 6E–F**).

These results, together with differential expression testing, demonstrate the versatility of Kompot’s cluster-free framework in quantifying changes in both gene expression and cell-state composition between conditions.

## Discussion

Kompot is a framework for comparing single-cell phenotypic landscapes between conditions without first partitioning cells into clusters or cell types. Kompot represents gene expression as a continuous function of cell state, learned separately for each condition by Gaussian process regression, and compares these functions to derive a per-cell log fold change for every gene. Each function is defined over the entire manifold and can therefore be evaluated at all cell states irrespective of the condition. The two conditions are then contrasted at every cell for differential expression testing. Differential abundance testing follows from the same procedure applied to cell-state density in place of expression. Kompot therefore provides a unified framework for systematic comparison of multi-condition single-cell datasets.

Kompot resolves the trade-off that discretization imposes. Testing cluster by cluster requires clusters broad enough to estimate a mean reliably, yet any cluster broad enough to do so averages over the continuous variation within it. A change confined to part of a cluster is diluted, a change that reverses direction inside one cancels, and biological variation among the sub-states a cluster combines is treated as technical noise. Kompot avoids this trade-off by never aggregating before testing. To assess significance, Kompot summarizes the per-cell fold changes with the Mahalanobis distance. Fold changes at neighboring cell states are not independent, and the Mahalanobis distance combines them while accounting for the posterior covariance the Gaussian process induces between them. Coordinated low-magnitude gene expression changes therefore accumulate into a single score instead of averaging away, and genes that change in opposite directions across subpopulations are also recovered. Calibrating this distance against a gene-shuffling empirical null with local-FDR control leads to a significance measure that holds across diverse datasets and parameter choices.

This resolution changed what we could interpret in each system we examined. In murine hematopoietic aging, only the antigen-processing genes are upregulated nearly uniformly across all cell types. Ranking by Mahalanobis distance revealed that most other age-altered genes, involved in hematopoietic function and antioxidant programs, show only a subtle aggregate change of two kinds. In one, the direction varies across cell types. In the other, the change is confined to stem cells. Resolving the latter along the neutrophil lineage showed that the aging changes in granulocyte-associated genes likely do not persist, with expression returning to near-young levels after cells exit the stem-cell state. Neither pattern would be detected by a test that reduces each cell type to a single mean, and only per-cell fold changes recover them.

The per-gene summary enabled a different kind of correction in *Tal1^−/−^* mouse embryos, where genotype is confounded with donor origin by the chimera design. Since Kompot summarizes each gene by a single Mahalanobis distance across the manifold, the mutant chimera can be compared directly against a wild-type chimera processed identically, and genes ranked high in both can be excluded as design artifacts rather than effects of *Tal1* loss. The changes that remain show that *Tal1* loss extends into morphogenesis programs across many embryonic lineages, beyond the hematoendothelial compartment it is known to regulate.

In advanced melanoma, resolving fold change by cell state showed that dual checkpoint blockade acts on largely the same genes as either monotherapy but redistributes their changes across the CD8 compartment. It raises T-cell activation broadly while suppressing translation in the cycling cells that would ordinarily upregulate it, a response neither monotherapy produces. This non-additive decoupling is detectable only because Kompot resolves each gene’s fold change across cell states rather than averaging it within clusters.

Kompot also supports differential abundance testing at single-cell resolution and assigns significance to every compositional change, without predefining clusters. We applied Kompot to hematopoietic aging and identified two phases that a single young-to-old contrast merges. Early aging remodels the mature compartment, where both B and T cells move from naïve to memory states while the progenitor compartment is unchanged. Late aging shifts the changes to the progenitors, most prominently an accumulation of hematopoietic stem cells.

Kompot has four principal limitations, each with a practical mitigation (**Supp. Note 5**). First, as in any Bayesian procedure, uncertainty due to model selection is not fully represented. Since the Gaussian process is non-parametric, this reduces largely to the covariance kernel and its length scale, and Kompot results are robust across a wide range of length scales. Second, Kompot inherits the quality of the input latent representation, and if the two conditions are misaligned it will identify the misalignment as biology. This is the same correspondence problem that batch correction addresses, and it can be diagnosed and corrected in the same way. Third, Kompot models log-transformed expression with a Gaussian rather than an explicit count distribution, so it does not model count overdispersion directly. In practice this matters little since an empirical, cell-state-dependent variance absorbs most of it and barely changes gene rankings (**Supp. Note 5**). Fourth, Mahalanobis distances should not be used to rank genes at the lower end, where sparsely measured genes pick up small distances that reflect measurement sparsity rather than biology. This does not impact any of our results, since all discoveries come from the high-distance genes, and the effect shrinks further as the length scale is increased (**Supp. Fig. 21**).

## Methods

### Kompot framework

Kompot is a toolkit for comparing single-cell phenotypic manifolds across conditions. Its core capability is differential gene expression testing at single-cell resolution, without requiring predefined cell types or clusters, and the same framework extends naturally to differential abundance testing. The guiding assumption is that both gene expression and cell-state abundance change continuously along cell-state space [14].

The input to Kompot is a latent representation of co-embedded multi-condition single-cell data (e.g. PCA, Harmony [9], scVI [15], LSI [52]). When needed, upstream batch correction is used to align equivalent biological states across conditions so that cells of similar state occupy similar locations in latent space. This alignment does not remove condition-specific gene-expression differences; rather, it standardizes the coordinate system so Kompot can compare cells we deem equivalent by proximity in state space [10].

Diffusion map representations are used as the default cell-state representation because Euclidean distances in diffusion components capture biologically meaningful differences between cells [11, 13, 53], but Kompot can operate on any embedding with a well-defined distance metric.

Kompot models condition-specific expression profiles and cell-state densities with Gaussian processes, enabling differential analysis that accounts for uncertainty and structured variation along the phenotypic manifold. This allows detection of consistent, state-dependent expression or abundance shifts that may be smaller than the feature’s global variability, including cases where local effects differ in direction (e.g., reduced dynamic range). By explicitly estimating posterior uncertainty via variational inference, Kompot provides principled significance measures for its findings at minimal additional computational cost.

#### Differential expression testing

Differential expression testing identifies gene expression changes between two conditions across a shared manifold. The goal is to detect genes that differ between conditions, rather than between cell types or clusters within a single condition, and in particular to capture subtle, structured transcriptional shifts that vary across cell states even when they do not follow a consistent direction. For each gene in each condition, Kompot constructs a condition-specific expression function that maps the diffusion-map representation to observed expression using Mellon [14]. Since these functions are continuous over the manifold, testing spans the entire phenotypic landscape and integrates all cells jointly, rather than partitioning them into predefined clusters.

We leverage Mellon’s Gaussian process implementation for these gene-expression functions (**Supp. Note 1**). For each condition, Kompot computes a function that predicts an expression value for each gene at any location in the cell-state space. These functions provide a smooth, continuous representation of gene expression across the manifold and integrate all data points without requiring predefined cluster boundaries. The resulting gene-expression functions can be used to compute local gene-expression differences, with local fold change providing the effect size for each gene at each cell-state.

Formally, Gaussian processes map the expression values for each gene *g* at cell states 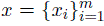 through the function *f_g_*(*x*). Under our modeling assumption, the true expression values *f_g_*(*x*) are distributed according to a multivariate normal posterior:

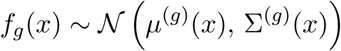

Therefore, the difference between conditions *a* and *b* also follows a normal distribution:

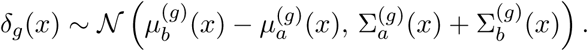

The posterior mean difference 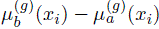 defines the local expression fold change at each cell state and serves as the effect size.

Significance is not assessed cell state by cell state and instead Kompot summarizes expression differences of gene over the whole manifold into a single statistic. Gaussian process expression functions aggregate information across cells since they map state space to observed expression. Specifically, for gene *g*, the gene expression values from function *f_g_*(*x_i_*) at different cell states *x_i_* are not *independent* measurements that can be aggregated directly. Kompot resolves this by explicitly modeling covariance across states using Mellon’s uncertainty framework. Mellon provides not only variances for each predicted expression value, but also covariances that reflect the smoothness of gene expression across the manifold i.e., the uncertainty covers not only the variance for a function value *f* (*x_i_*) but also the covariance to other function values *f* (*x_j_*) based on the similarity of cell states *x_i_* and *x_j_*. To summarize these correlated local differences into a single per-gene statistic, Kompot uses the Mahalanobis distance (**Supp. Note 2**):

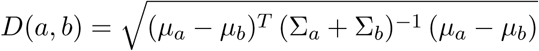

This statistic accounts for the covariance between similar cell states, ensuring that consistent changes are emphasized while redundant information is down-weighted. As a result, Kompot can detect differential expression patterns across the manifold, even when changes differ in magnitude or direction.

*Thresholds to identify significantly changed genes.* To enable robust identification of significantly changed genes, Kompot generates an empirical null distribution of Mahalanobis distances by randomly shuffling gene expression values (default: 2000 genes). From the resulting null distribution, empirical p-values are computed and used to estimate false discovery rates (FDR) via local FDR methods [54]. By default, Kompot applies a cutoff of 0.05 on the local FDR, yielding a direct Mahalanobis distance threshold for identifying genes significantly differentially expressed (**Supp. Note 3**). The behavior of the null-density estimator that converts distances to local FDR is shown in **Supp. Fig. 22**.

#### Differential abundance testing

The same posterior-difference framework extends from gene expression to the abundance of cell states across the phenotypic manifold. Instead of a per-gene expression function, Kompot compares condition-specific density functions that describe how cells from each condition are distributed across the manifold. Because these densities are continuous, the comparison spans the entire phenotypic landscape, enabling cluster– and cell-type–free abundance testing at single-cell resolution.

Kompot estimates each condition’s density with Mellon [14]. While Mellon was previously introduced as a framework for continuous density estimation, here we extend it with explicit uncertainty estimation at each cell state. This Bayesian inference procedure yields, separately for each condition, a best-estimate function mapping each cell state *x_i_*to a log density value *µ*(*x_i_*) together with a quantification of uncertainty *σ*(*x_i_*) (**Supp. Note 1**). As for expression, the change in density between conditions *a* and *b* at cell state *x_i_*is normally distributed,

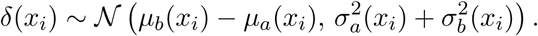

*Normalization for comparability across conditions.* Mellon density estimates are in units of log number of cells per volume, so direct comparisons between conditions with different numbers of cells are not accurate. Kompot therefore transforms these values into log fraction of cells per volume by subtracting the log total number of cells,

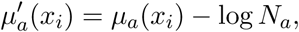

where *N_a_*is the total number of cells measured for condition *a*. This ensures density values reflect the relative occupancy of the phenotypic manifold independent of the number of cells; it assumes the intrinsic dimensionality of the manifold is estimated correctly, which Mellon provides and treats as constant across the dataset. The normalized local log fold change representing the abundance difference is therefore

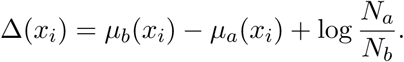

*Significance testing.* Since abundance is described by a single density function rather than one function per gene, significance is assessed cell state by cell state rather than summarized into a per-gene statistic. At each cell state, Kompot computes a posterior tail probability (PTP): the probability that the true change has the opposite sign to the estimated change Δ(*x_i_*). Since the posterior is normal,

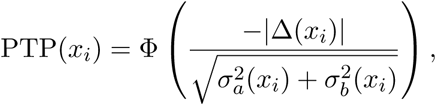

where Φ is the cumulative distribution function of the standard normal distribution. Kompot thereby provides both an effect size (normalized density log fold change) and a cell-wise significance measure for every cell state.

### Benchmarking of differential expression methods

We constructed a semi-synthetic benchmark from the aging haematopoietic dataset by spiking a known log fold change patterns into a fixed set of 500 genes per scenario and asking each method to recover them, using eight simulated replicates per condition so that between-sample variability is realistic. The 500 spiked genes are 3.07% of the 16,285 tested genes, approximating a realistic differential-expression rate. Five scenarios span the range of patterns the method should handle. Ten methods were compared: Kompot, MiloDE [5], scVI-DE [15], LEMUR-unpaired [6], MAST, NEBULA, pseudobulk DESeq2 and edgeR on unsupervised Leiden clusters, and bulk DESeq2 and edgeR. Each was scored by AUROC and AUPRC on its ability to rank the 500 spiked genes above the remaining 15,785, so the AUPRC chance baseline is 0.0307. Scenario construction, per-scenario effect sizes and the gene-selection constraints are described in more detail in **Supp. Note 4**.

The pseudobulk comparators, DESeq2 and edgeR, were run on unsupervised Leiden clusters [21] rather than on the manual cell-type annotation, so that no method is given ground-truth labels. The annotation was itself produced by clustering these same cells, so the Leiden clusters still fall close to the boundaries the scenarios were built around. Pseudobulk performance reported here is therefore an optimistic estimate of what these methods would achieve on data where the clustering and the effect are not aligned.

### Kompot Scalability

Kompot inherits the scalability of the Mellon framework through its sparse Gaussian process implementation, enabling efficient inference across datasets spanning several orders of magnitude in size. We systematically benchmarked runtime scaling across four representative datasets: *aging* (*∼*8,000 cells, 16,000 genes), *BCR-XL* (CyTOF dataset of *∼*172,000 cells, 24 proteins) [55], *COVID-19* (*∼*422,000 cells, 34,000 genes) [16], and a *post-COVID* variant comprising the same cells but distinct condition comparisons.

To ensure consistent memory reporting, all memory values correspond to the maximum resident set size (MaxRSS) recorded by Slurm. MaxRSS robustly captures total host RAM usage across all components involved in execution, including pure Python allocations and memory consumed by NumPy, JAX, and Dask kernels. Importantly, MaxRSS does not include GPU device memory; VRAM usage during accelerated runs was therefore not tracked in these benchmarks, and the reported values reflect only CPU-side memory consumption.

#### Scaling with number of cells

Runtime is primarily governed by the number of cells in the compared conditions. Scaling with number of cells were determined using default parameters including 5000 landmarks. When the number of cells exceeds the number of landmarks (default: 5,000), the sparse Gaussian process formulation achieves approximately linear scaling with number of cells (**Supp. Figs. 5–8, 16–19 A, Supp. Fig. 4A, E**). While differential abundance runtime depends almost exclusively on the number of cells and landmarks, differential expression additionally scales with the number of genes analyzed.

#### Scaling with number of landmarks

Runtime increases quadratically with the number of landmarks *k*, consistent with the expected *O*(*k*^2^*n*) complexity of sparse Gaussian processes for a number of cells *n* (**Figs. 2B, D Supp. Fig. 4B, F**; memory in **Supp. Fig. 3B, F**). The quadratic trend becomes apparent above *∼*1,000 landmarks, once constant overhead is exceeded. When the landmark count surpasses the number of cells, Kompot reverts to a full Gaussian process, resulting in constant runtime independent of further increases in *k*. Differential expression was performed with gene batching, so the curves diverge slightly, since gene batching reduces memory requirements at the cost of increased per-batch computation time.

#### Scaling with number of genes and parallelization

Differential expression runtime depends only weakly on the number of genes (**Fig. 2E**, **Fig. 2F, Supp. Fig. 4D**). The total complexity is *O*(*nk* + *nkg*), where *n* is the number of cells, *k* the number of landmarks, and *g* the number of genes. The shared *O*(*nk*) operations (covariance construction and decomposition) are performed once, while the per-gene *O*(*nkg*) term scales linearly with *g*. Each gene is modeled by an independent Gaussian process with closed-form inference, making computation trivially parallelizable.

#### Landmark generation

As in Mellon [14], Landmarks are computed using *k*-means clustering, which scales as *O*(*tkdn*) with *t* iterations, *k* landmarks, *d* dimensions, and *n* cells. For very large datasets, this step can become a bottleneck. Kompot therefore allows users to provide custom landmarks, enabling faster or approximate alternatives. Because inference relies on covariance statistics, the exact landmark placement has limited influence as long as landmarks adequately cover the state space. Redundant landmarks contribute negligibly to test statistics, as their strong covariance prevents inflation of significance.

### Peak memory usage and runtime analysis

Peak memory usage was measured using SLURM’s sacct utility (MaxRSS), which captures the maximum resident set size across all associated processes (JAX, Dask, NumPy, Python). Runtime was measured as the total execution time of Kompot on the AnnData object. Benchmarks were performed on the *covid-19* dataset. Each configuration was repeated 10 times to characterize the variance differences in memory pressure and hardware availability across HPC nodes. Kompot was run with single CPU, 16 CPUs and single GPU to assess the scalability.

### Ethics statement

All animal experiments were conducted in accordance with protocols approved by the Institutional Animal Care and Use Committee (IACUC) of Fred Hutchinson Cancer Center (protocol #[NUMBER]). C57BL/6 mice were maintained under standard housing conditions. The neuroblastoma single-cell RNA-seq data were obtained from Yu et al. [56], which was conducted under IRB approval at the Children’s Hospital of Philadelphia. The melanoma clinical trial data were obtained from Cillo et al. [43], conducted under IRB approval at the University of Pittsburgh.

### Aging hematopoiesis single-cell RNA-seq data generation

#### Marrow collection

Bone marrow mononuclear cells were collected from the femur, tibia, and hip bones from male C57BL/6 mice aged young (10 weeks, n=3), mid (63 weeks, n=2), and old (103 weeks, n=3). Whole marrow samples were washed twice in PBS supplemented with 10% FBS and were gently passed through a 40-micron strainer. The cell suspension was split 1:10 with one part set aside for FACS sorting of mature hematopoietic cells, and the rest for hematopoietic stem and progenitor cell (HSPC) sorting.

#### HSPC processing

Cells intended for HSPC sorting were first subjected to MACS lineage depletion with BioLegend antibodies targeting the following mature lineage marker antigens: Ter119 (CAT# 79748), B220 (CAT# 103204), CD3e (CAT# 79751), CD4 (CAT# 100404), CD8a (CAT# 100704), CD11b (CAT# 79749), Gr1 (CAT# 79750), Fc*ε*R1*α* (CAT# 134304), CD19 (CAT# 115504), NK1.1 (CAT# 108704), CD11c (CAT# 117304). Cells were incubated in lineage stain for 10 minutes, washed in MACS buffer (0.5% bovine serum albumin and 2 mM EDTA in PBS), then incubated for another 15 minutes with anti-biotin MicroBeads (5 *µ*L for every 10e7 cells, Miltenyi Biotec CAT# 130-090-485), washed in MACS buffer, and passed through a 40 micron filter immediately prior to processing with the AutoMACS Pro Separator. After MACS lineage depletion, samples were incubated for 10 minutes with TruStain FcX PLUS (1%, BioLegend CAT# 156604), then each sample was stained for 30 minutes with a cocktail of (1) a unique hashing antibody (BioLegend CAT#s 155831, 155833, 155835, 155837, 155839, 155841, 155843, or 155845) plus (2) the flow sorting antibodies mCD45, anti-biotin and cKit plus (3) BioLegend CITE Total Seq antibodies targeting the following: CD135 (CAT# 135319), CD16/32 (CAT# 101345), Sca1 (CAT# 108149), CD127 (CAT# 135055), CD150 (CAT# 115951), CD48 (CAT# 103457), CD41 (CAT# 133941), CD34 (CAT# 128625). Samples were pooled then washed three times in Cell Staining Buffer (BioLegend CAT# 420201) then were sorted for DAPI (*−*), Ter119 (*−*), mCD45 (+), lineage (*−*), cKit (+).

#### Mature processing

Each of the 8 samples intended for Mature sorting were stained for 30 minutes with a cocktail of (1) a unique hashing antibody (BioLegend CAT#s 155831, 155833, 155835, 155837, 155839, 155841, 155843, or 155845) plus (2) the flow sorting antibodies mCD45, and Ter119 plus (3) BioLegend CITE Total Seq antibodies targeting the following: CD4 (CAT# 100573), CD8a (CAT# 100783), Ly-6C (CAT# 128053), CD11b (CAT# 101273), Ly-6G (CAT# 127659), CD44 (CAT# 10371), CD19 (CAT# 115563), CD25 (CAT# 102067), B220 (CAT# 103271), CD115 (CAT# 135543), CD11c (CAT# 117359), CD62L (CAT# 104465), F4/80 (CAT# 123155), I-A/I-E (CAT# 107657), NK1.1 (CAT# 108763), CD3 (CAT# 100257), CD335 (CAT# 137641), CD27 (CAT# 124247), CCR2 (CAT# 150633), CXCR4 (CAT# 146521), C3CR1 (CAT# 149045). Samples were pooled then washed three times in Cell Staining Buffer (BioLegend CAT# 420201) then were sorted for DAPI (*−*), Ter119 (*−*), mCD45 (+).

#### Sequencing

Single cell barcoded libraries were generated using chromium Next GEM Single Cell 3’ Reagent Kits, Dual Index (v3.1 Chemistry) and were sequenced on an Illumina HiSeq.

## Data Analysis

### Aging hematopoiesis scRNA-seq

*Preprocessing.* CellRanger (v6.1.2) was used for demultiplexing and count matrix generation. We next performed quality control to remove low-quality cells. Doublets were identified using Scrublet (v0.2.3) [57]. Suspected apoptotic cells were identified as any cell with 10% mitochondrial reads, and cells with 20% or more molecules derived from hemoglobin genes were identified as red blood cell contamination. Cells with abnormally low total molecule counts were also flagged. All cells failing any QC metric were removed. Genes expressed in fewer than 10 cells were excluded.

After QC, the dataset was processed in Scanpy as follows. Raw gene expression counts were normalized using the Scanpy [58] normalize_total function so that all cells have the same total count over all genes. A pseudo-count of 0.1 was added and base-2 logarithms were taken. The top 25% most highly variable genes (HVGs) were selected using the Scanpy highly_variable_genes function with the cell_ranger flavor and batch_key=Compartment (HSPC vs. Mature compartments). Cell cycle scores were computed using the Scanpy score_genes_cell_cycle function, then the cell cycle effect was regressed out with the Scanpy regress_out. PCA was carried out on the HVGs, and batch correction across compartments was performed with Harmony [9]. Neighborhood graphs were computed from the Harmony-corrected PCs, followed by UMAP embedding and Leiden clustering. Cell types were assigned manually based on marker gene expression obtained from public data sources

*Differential Abundance, Differential Expression, and Gene Set Enrichment Analysis.* Diffusion maps were computed with Palantir [12] using 40 diffusion components derived from the Harmony-corrected PCA representation, and this diffusion map embedding served as the latent cell-state representation for all Kompot analyses. Differential abundance was computed with Kompot using default parameters (ls_factor = 10, n_landmarks = 5000) for each pairwise comparison between the three age groups (Young vs. Old, Young vs. Mid, and Mid vs. Old). Differential expression was performed on the same comparisons with Kompot defaults (ls_factor = 10, n_landmarks = 5000, sigma = 1). For DE, we used the Scanpy-normalized expression matrix described above, applying a log10 transformation to normalized counts after adding a 0.1 pseudocount, and included all 16,285 genes in the analysis.

Gene set enrichment analysis was carried out using the Kompot StringDBReport function, which interfaces with the STRING [59] database through its API. Gene names were mapped to STRING protein identifiers using the default mapping, unmapped genes were skipped, and enrichment was computed using STRING’s default annotation categories for any gene set derived from the Kompot DE results.

### Mouse gastrulation data

Processed and annotated data for the *Tal1^−/−^* chimera and the matched WT-only chimera (Host vs. tdTomato^+^ ESC-Injected) from E8.5 mouse embryos were downloaded from ArrayExpress (E-MTAB-7325) [37]. For each chimera independently, cell types exclusively present in one condition (*>*99%) were excluded so that the comparison was restricted to lineages with sufficient cells in both groups (notably this removes all erythroid lineages from the *Tal1^−/−^* chimera, which lacks them by construction; the same exclusion was applied to the WT chimera so the two analyses share a canonical cell-type subset). PCA was recomputed on the cleaned data, batch-corrected with Harmony [9] on a composite [tomato, sample] key (sample– and condition-aware), and used to compute diffusion maps with 50 components. Cell-cycle phases were determined with Scanpy’s score_genes_cell_cycle on the Tirosh et al. signature [60], scored on all cells of a chimera

Kompot differential expression was run with default parameters in each chimera. Per-gene Mahalanobis distance was calibrated against the gene-shuffling null and converted to a local FDR; the *Tal1* significance cutoff was set to local FDR *<* 0.05. To remove chimera-experiment artefacts we used the WT-only chimera as a Mahalanobis-matched null: with *T* = min(Mahalanobis) across the *Tal1* hits, any *Tal1* hit whose WT-chimera Mahalanobis was *≥ T* was flagged as a chimera artefact; in addition, a fixed exclusion list of imprinted and sex-linked genes (*Xist*, *Cdkn1c*, *Phlda2*, *Grb10*, *H13*, *Nnat*, *Impact*, *Kcnq1ot1*, *Slc38a4*, *Meg3*, *Igf2*, *Igf2r*, *Dlk1*, *Peg3*, *Peg10*, *Mest*, *Plagl1*, *Zdbf2*, *Gt(ROSA)26Sor*) and the *Hb[ab]\** hemoglobin paralogues was removed. The remaining “clean Tal1 DE” set is used for all downstream interpretation.

STRING enrichments were run against an *explicit* background of the genes actually tested in the object rather than STRING’s genome-wide default. Reported GO terms are those with FDR *<* 0.05, restricted to terms with at least 5 genes in the query set and at most 1,000 in the background, and ranked by FDR; the top 15 are shown. FDR rather than enrichment strength is used for the ranking, since strength has no size bound and is maximised by very small terms, which would promote marginal categories over the strongest ones.

### Melanoma CD8 T cell landscape

Processed and annotated data was downloaded from GEO (GSE225063) [43]. The dataset consisted of 75160 cells spanning different CD8 T cell states from melanoma patients at baseline and following treatment with different ICI. All cells from baseline, week4 and week16 timepoints were used for analysis (74644 cells), with the union of week4 and week16 cells used as cells in the response condition. Pre-computed scVI latent representation was used to compute diffusion maps with 10 components.

Kompot differential expression testing was performed separately for each treatment arm (Baseline *→* Response) using genes detected in at least 100 cells in the dataset (15,247 genes). Significance was assessed at the calibrated local-FDR cutoff (gene-shuffling null, default *<* 0.05), yielding 1,745 DE genes in the Nivo arm, 1,065 in Rela, and 1,040 in Rela+Nivo. STRING queries were performed the human species identifier (9606).

Genes highlighted in **Fig. 5C** were derived as follows. Genes were ranked by Mahalanobis distance within the combo arm and the leading 12 are shown, partitioned into up-, down– and subtle-changing classes. Before that ranking, a fixed pattern filter removes gene families that dominate single-cell rankings by abundance rather than by treatment biology: cytoplasmic and mitochondrial ribosomal proteins, mitochondrially encoded genes, histone-cluster genes, translation elongation and initiation factors, beta-thymosins, heat-shock chaperones, and ubiquitous cytoskeletal housekeeping genes (*ACTB*, *ACTG1*, *TUBB*, *TUBA1A–C*, *GAPDH*, *B2M*). Cytotoxicity genes (*PRF1*, *GZMB*, *NKG7*) are deliberately retained, since cytotoxicity is the biology under study and removing them would remove the result; noncoding transcripts and unnamed Ensembl identifiers are likewise retained.

#### Software versions

Analyses were performed using Kompot v0.8.0, Mellon v1.7.1, Palantir v1.4.5, Scanpy v1.10.4, and Python 3.11. Diffusion maps were computed with Palantir. Pseudobulk comparisons used DESeq2 via PyDESeq2 v0.4.12. Exact versions of all Python and R dependencies, including the comparison methods (MiloDE, edgeR, NEBULA, LEMUR, MAST), are mentioned in the companion code repository.

## Data Availability

Raw single-cell data for the murine aging hematopoiesis are available at GEO under accession GSE309924. Processed anndata objects with annotations are available to download from Zenodo [61]. The neuroblastoma single-cell data were obtained from Yu et al. [56]. The melanoma CD8 T cell data were obtained from Cillo et al. [43].

## Code Availability

Kompot is available as a Python module at https://github.com/settylab/Kompot. Jupyter notebooks detailing the usage of Kompot differential expression and differential abundance testing are available at https://github.com/settylab/kompot/tree/main/examples and documentation at https://kompot.readthedocs.io/en/latest/. Code for reproducing the figures and analysis are available at https://github.com/settylab/kompot_reproducibility|, for memory and run time scaling at https://github.com/settylab/kompot_scaling for differential expression benchmarking at https://github.com/settylab/kompot_de_benchmark, and for differential abundance benchmarking at https://github.com/settylab/kompot_benchmarkDA.

Code was developed with the assistance of generative AI (Claude, Anthropic). The authors reviewed and validated all code, analyses and results, and take full responsibility for the content of the published article.

## Acknowledgements

This study was supported by National Institute of General Medical Studies grant no. R35 GM147125 to M.S.; National Cancer Institute grant no. R01CA292932, Mark Foundation for Cancer Research Endeavor Award and Edward P. Evans Foundation Discovery Research Grant to S.C.L; Brotman Baty Institute Pilot Award and Translational Data Science IRC New Collaboration Award to M.S. and S.C.L; National Institute of General Medicine Studies grant no. K99GM159044 to D.O.; National Institute of General Medicine Studies grant no. T32GM136534 to E.T.; National Institutes of Health grant no. ORIP S10OD028685 to support high-performance computing at the Fred Hutchinson Cancer Center.

## Author Contributions

D.J.O.: Conceptualization, Methodology, Software, Formal Analysis, Writing – Original Draft, Visualization. E.A-G.: Investigation, Data Curation. E.T.: Investigation, Data Curation. R.Y.: Investigation. S.C.L.: Conceptualization, Supervision, Funding Acquisition, Writing – Review & Editing. M.S.: Conceptualization, Supervision, Funding Acquisition, Writing – Review & Editing.

