## Supplementary Figures for "Comparing phenotypic manifolds with Kompot: Cluster-free differential expression at single-cell resolution"

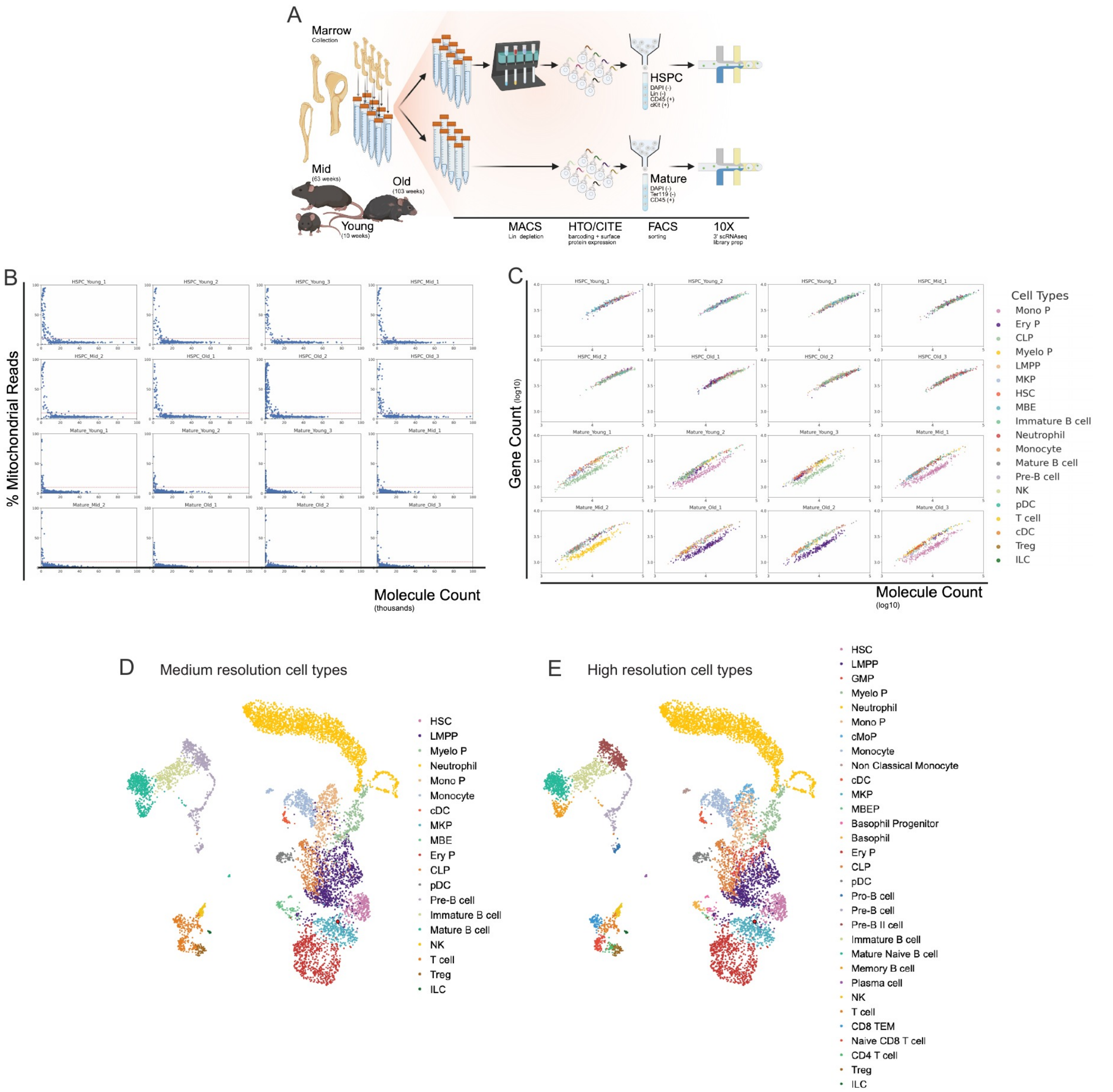

**Supplementary Figure 1: Generation and quality control of the murine aging hematopoiesis CITE-seq dataset.**

(A) Schematic of the experimental setup used to generate the dataset.

(B) Quality control, one panel per sample: total molecule count against the percentage of mitochondrial reads, one dot per cell. Cells above 10% mitochondrial reads were excluded from downstream analysis.

(C) Molecule count over gene count, one panel per sample and one dot per cell, colored by annotated cell type.

(D-E) UMAP from **Fig. 3A** colored by medium and high resolution cell types.

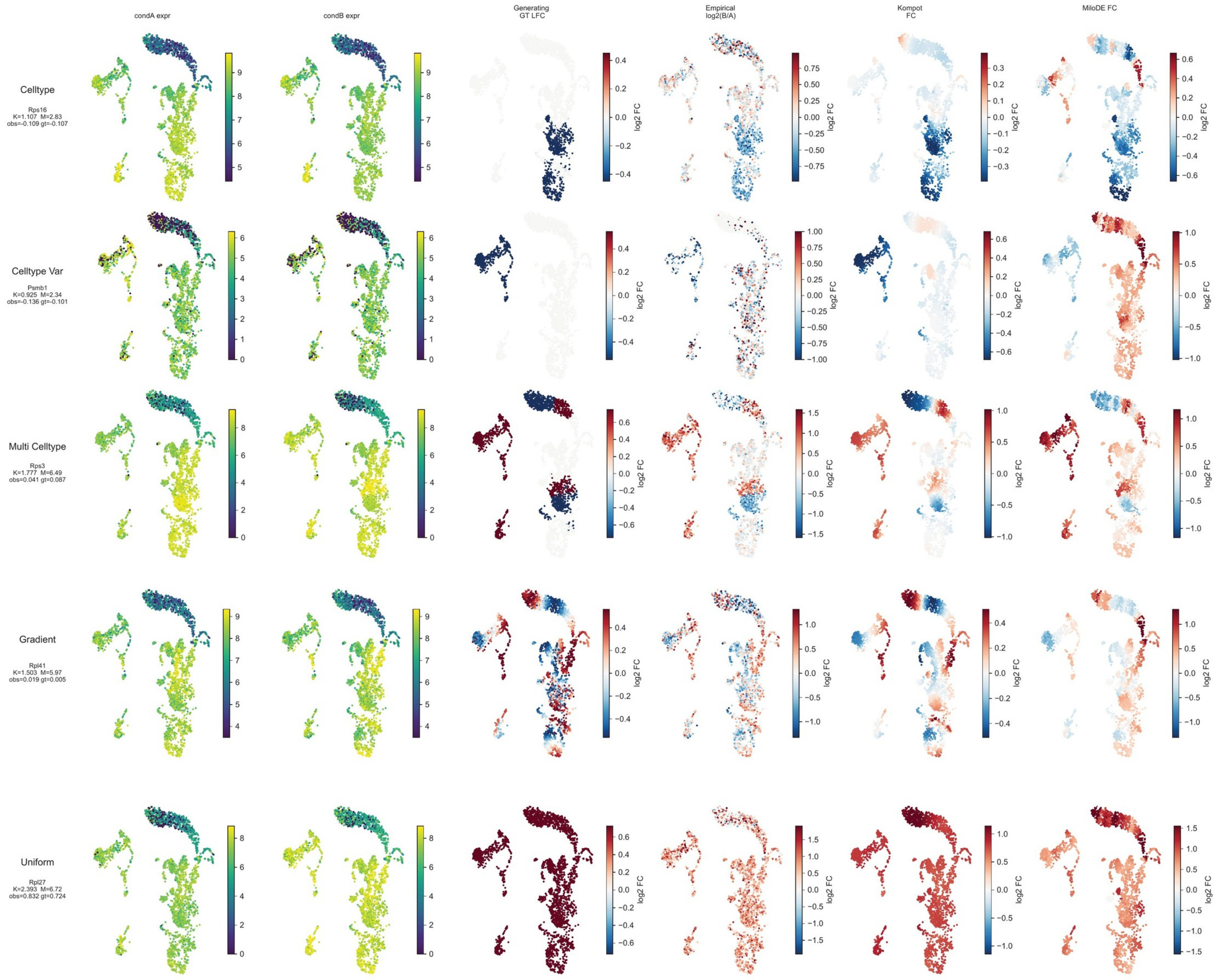

**Supplementary Figure 2: Spike-in patterns and their recovery in the semi-synthetic benchmark.**

One representative spiked gene per scenario, companion to **Fig. 2B**. Rows are the five reported scenarios: *Celltype*, *Celltype-var*, *Multi-celltype*, *Gradient*, and *Uniform*. Every panel is a UMAP of the young hematopoietic subset the benchmark is built from, colored, from left to right, by expression in condition A, expression in condition B, the generating ground-truth log fold change, the empirical $\log_2(B/A)$ after noise, the log fold change recovered by Kompot, and the log fold change recovered by miloDE.

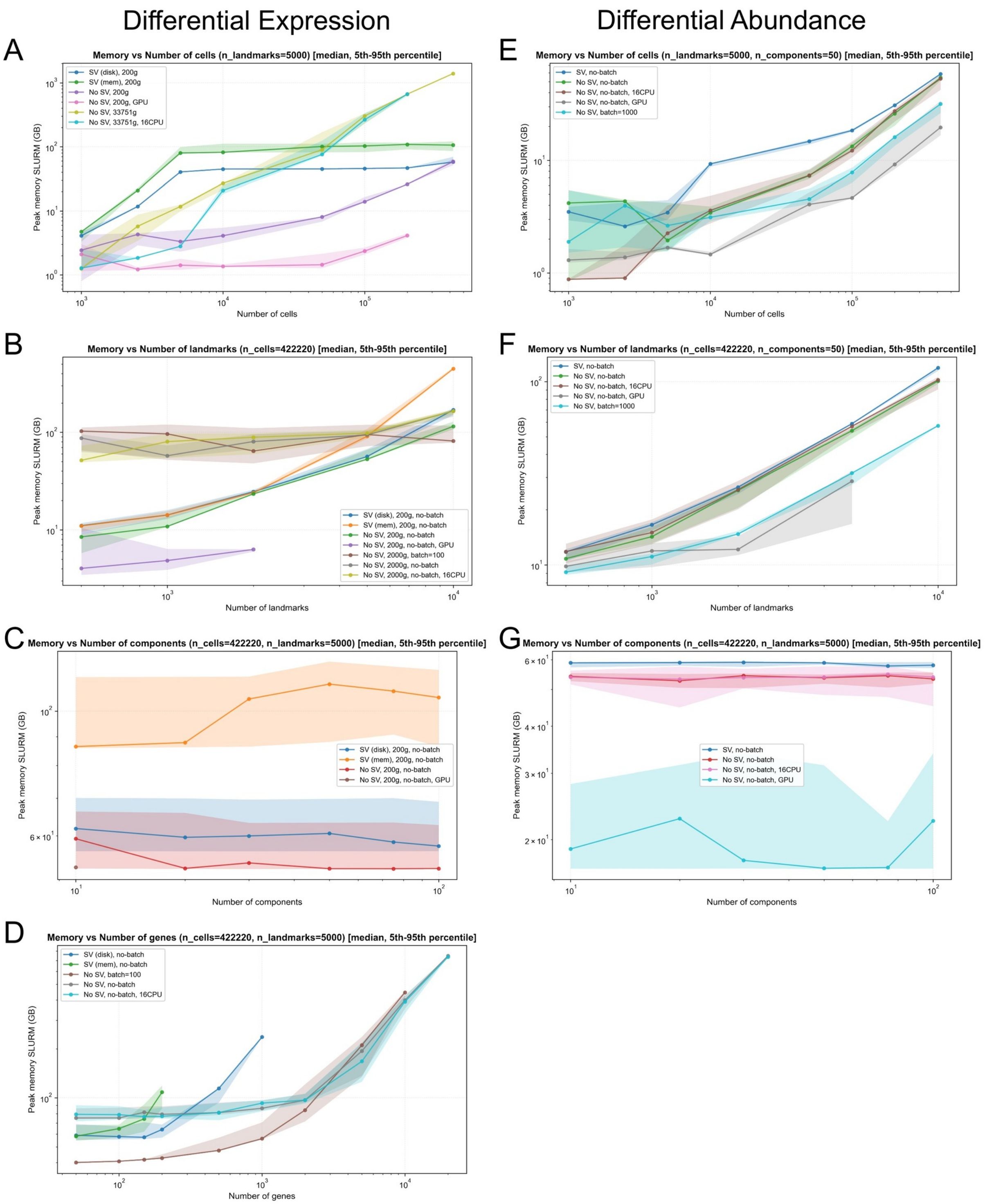

**Supplementary Figure 3: Kompot peak memory usage scaling analysis.**

All benchmarks were performed on the “covid” dataset (Healthy vs. COVID comparison). Lines show median run times for 10 experiments and shaded regions indicate the 5th–95th percentile intervals. SV: Sample variance estimation (the most computationally expensive option). Gene counts are abbreviated (e.g., “200g” = 200 genes), 16CPU: indicates parallel execution with 16 CPUs. GPU: GPU acceleration (NVIDIA L40S). The left column shows peak memory usage for differential expression analysis and the right column shows differential abundance analysis.

(A–D) Peak memory usage for differential expression analysis as a function of (A) number of cells using 5000 landmarks and 50 diffusion components, (B) number of landmarks using 422,220 cells and 50 diffusion components, (C) number of diffusion components using 422,220 cells and 5000 landmarks, (D) number of genes tested using 422,220 cells and 5000 landmarks.

(E–G) Peak memory usage for differential abundance analysis as a function of (E) number of cells using 5000 landmarks and 50 diffusion components, (F) number of landmarks using 422,220 cells and 50 diffusion components, (G) number of diffusion components using 422,220 cells and 5000 landmarks.

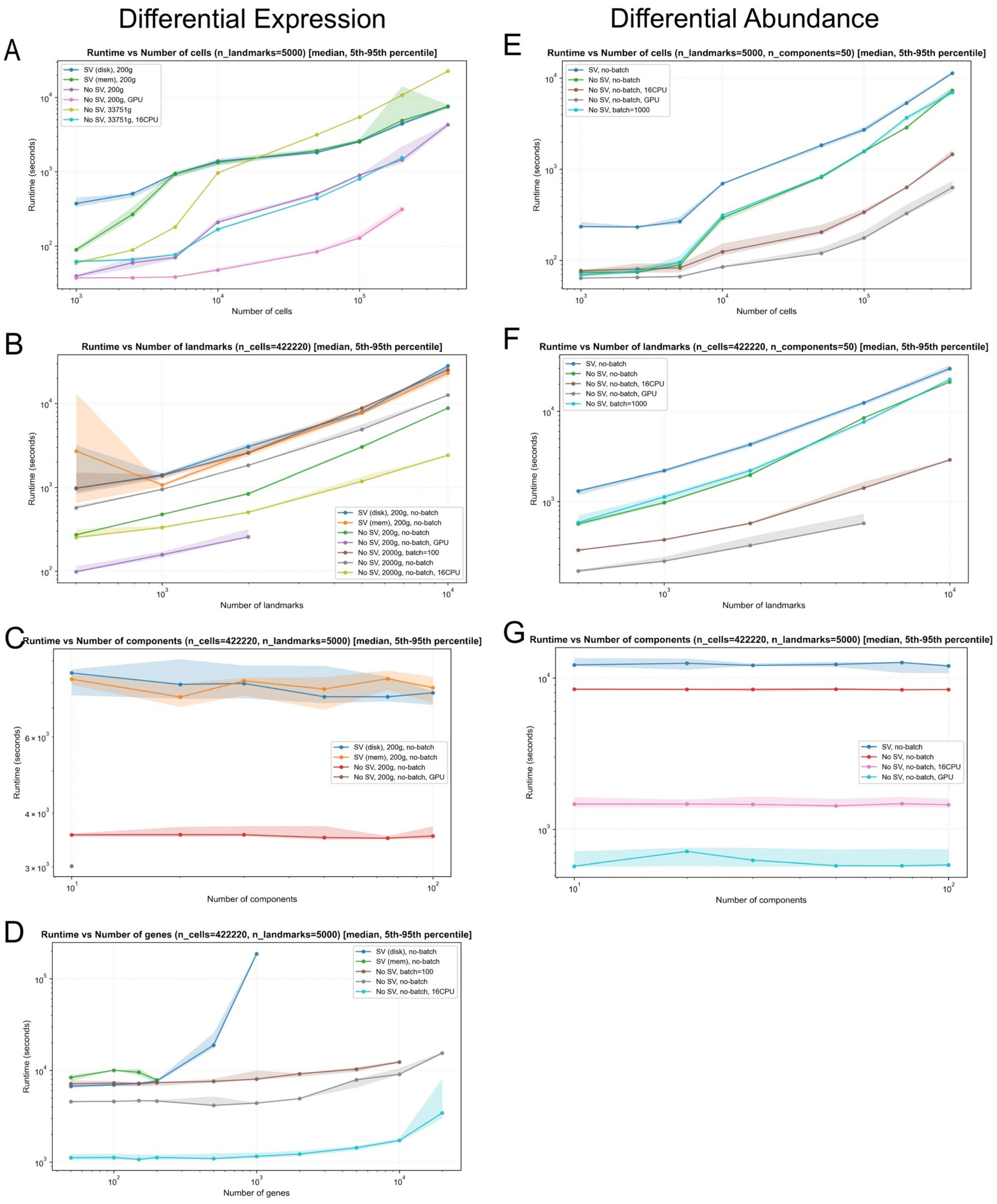

**Supplementary Figure 4: Kompot runtime scaling analysis.**

All benchmarks were performed on the “covid” dataset (Healthy vs. COVID comparison). Lines show median run times for 10 experiments and shaded regions indicate the 5th–95th percentile intervals. SV: Sample variance estimation (the most computationally expensive option). Gene counts are abbreviated (e.g., “200g” = 200 genes), 16CPU: indicates parallel execution with 16 CPUs. GPU: GPU acceleration (NVIDIA L40S). The left column shows run times for differential expression analysis and the right column shows differential abundance analysis.

(A–D) Runtime analysis for differential expression analysis as a function of (A) number of cells using 5000 landmarks and 50 diffusion components, (B) number of landmarks using 422,220 cells and 50 diffusion components, (C) number of diffusion components using 422,220 cells and 5000 landmarks, (D) number of genes tested using 422,220 cells and 5000 landmarks.

(E–G) Runtime analysis for differential abundance analysis as a function of (E) number of cells using 5000 landmarks and 50 diffusion components, (F) number of landmarks using 422,220 cells and 50 diffusion components, (G) number of diffusion components using 422,220 cells and 5000 landmarks.

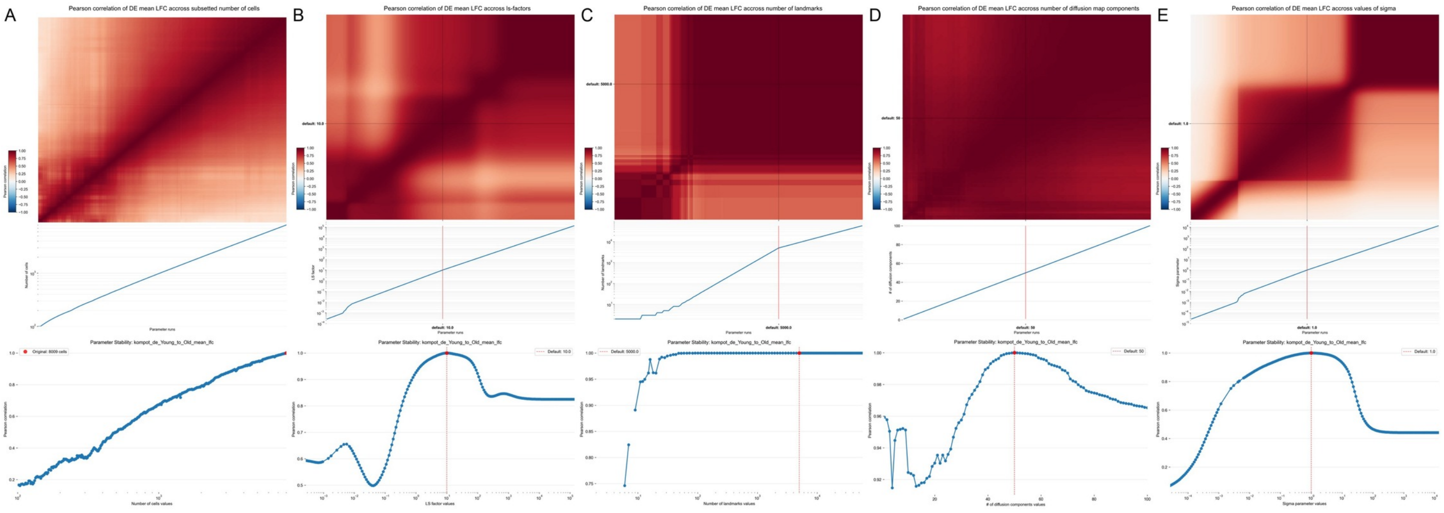

**Supplementary Figure 5: Robustness of Kompot differential expression mean log fold change for the hematopoietic aging dataset**

(A–D) Same as Supplementary Fig. 16 showing robustness of differential expression mean log fold change across all genes instead of abundance log fold change across all cells.

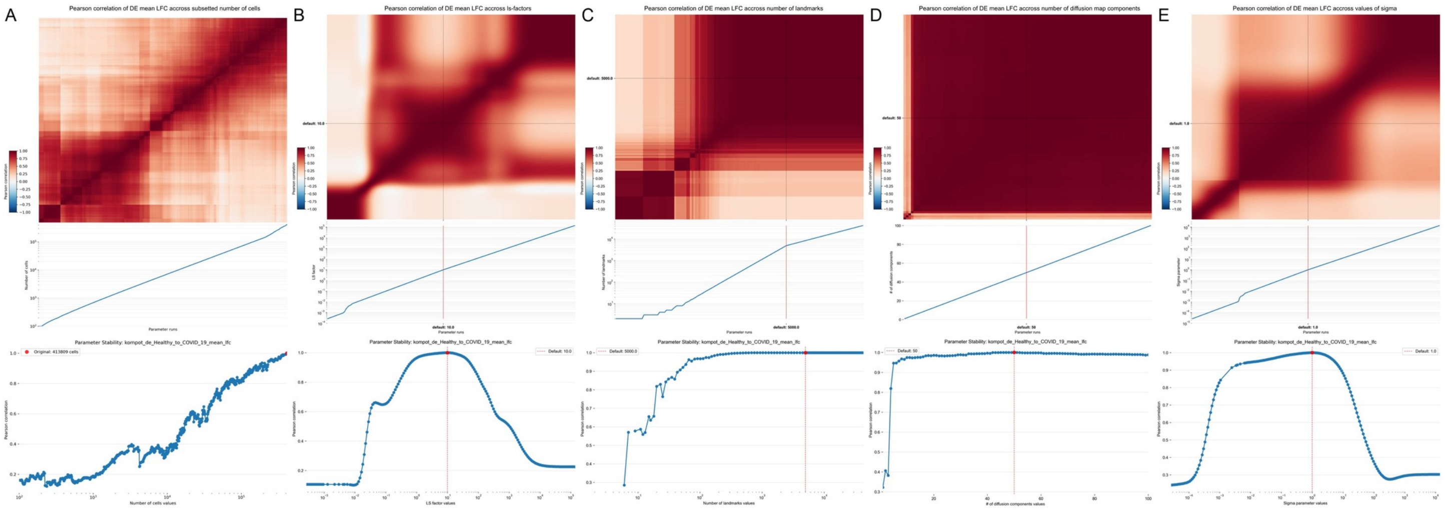

**Supplementary Figure 6: Robustness of Kompot differential expression mean log fold change for the COVID-19 dataset**

(A–E) Same as Supplementary Fig. 5 for the COVID-19 dataset.

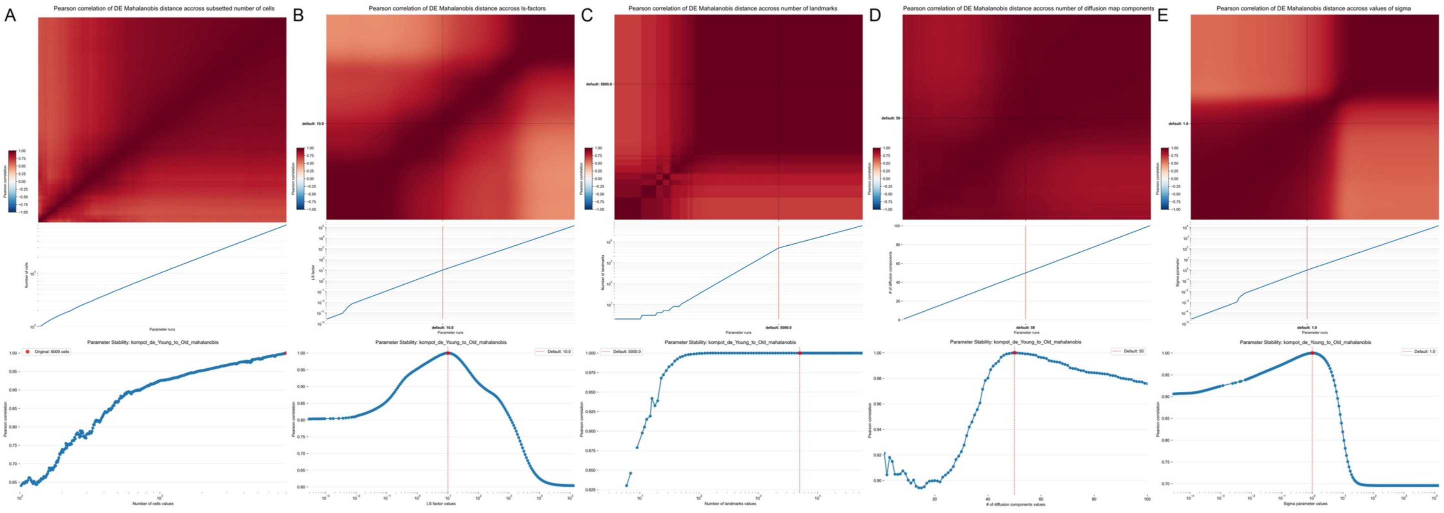

**Supplementary Figure 7: Robustness of Kompot differential expression Mahalanobis distance for the hematopoietic aging dataset**

(A–E) Same as Supplementary Fig. 5 showing robustness of Mahalanobis distance instead of mean log fold change.

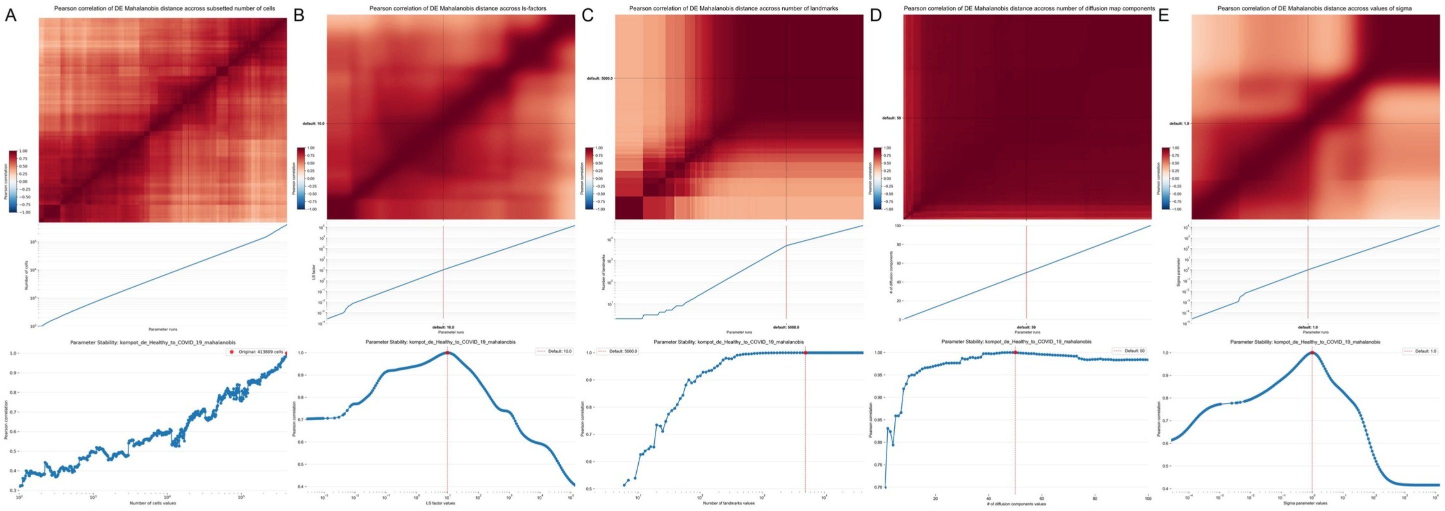

**Supplementary Figure 8: Robustness of Kompot differential expression Mahalanobis distance for the COVID-19 dataset**

(A–E) Same as Supplementary Fig. 7 for the COVID-19 dataset.

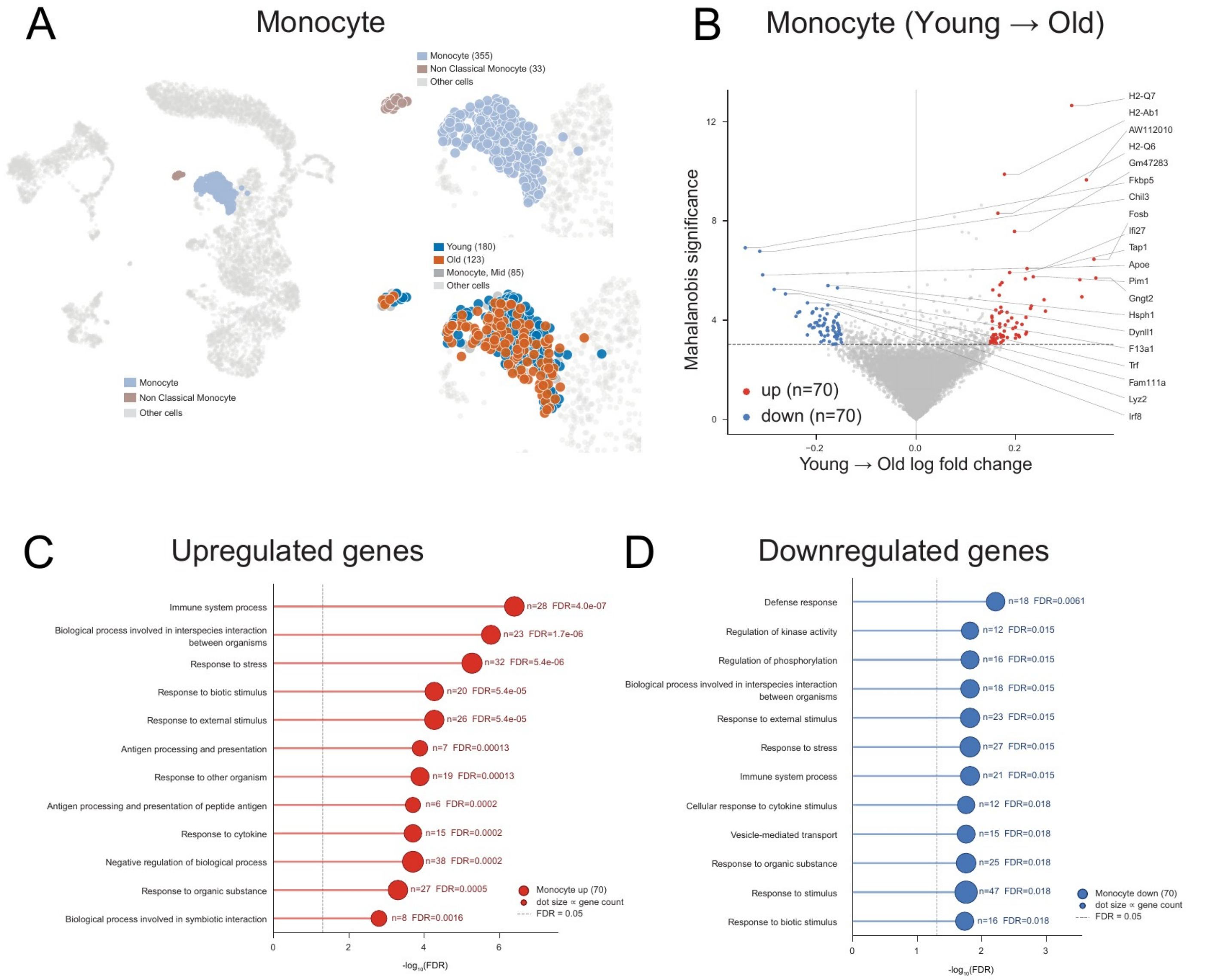

**Supplementary Figure 9: Kompot differential expression analysis of aging monocytes.**

(A) UMAP from **Fig. 3A** with only the monocytes colored. Right: Insets restricted to those monocytes, colored by cell type and by age.

(B) Kompot differential expression within the monocytes, young to old: mean log fold change against Mahalanobis distance, with the genes in the upper and lower quantile of the fold change highlighted.

(C) Gene-ontology enrichment of the *up*-regulated genes, dominated by antigen processing and presentation.

(D) Gene-ontology enrichment of the *down*-regulated genes, dominated by core monocyte and macrophage functions such as defense response.

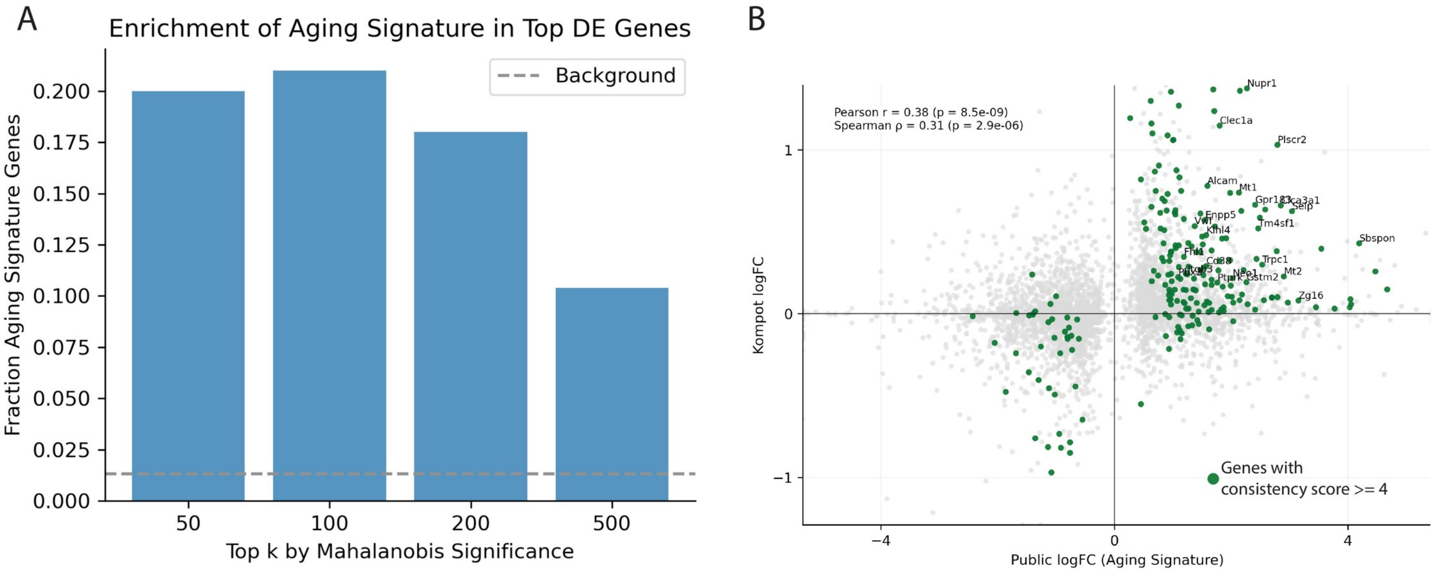

**Supplementary Figure 10: Consistency of Kompot differential expression results with hematopoietic aging atlas**

(A) Enrichment of aging-associated genes among Kompot's top differentially expressed genes. Bars show the fraction of genes within the top $k$ ranked by Mahalanobis significance that are part of a published hematopoietic aging signature, defined as genes with a reported consistency score $\geq 4$. The dashed horizontal line indicates the background frequency of such genes across all genes tested.

(B) Comparison of average log fold changes (logFC) per gene between Kompot (Mid to Old transition; y-axis) and the published aging signature (x-axis). Genes with consistency scores $\geq 4$ are highlighted in green; those with scores $\geq 8$ are additionally labeled. Pearson and Spearman correlation coefficients between the two datasets are shown.

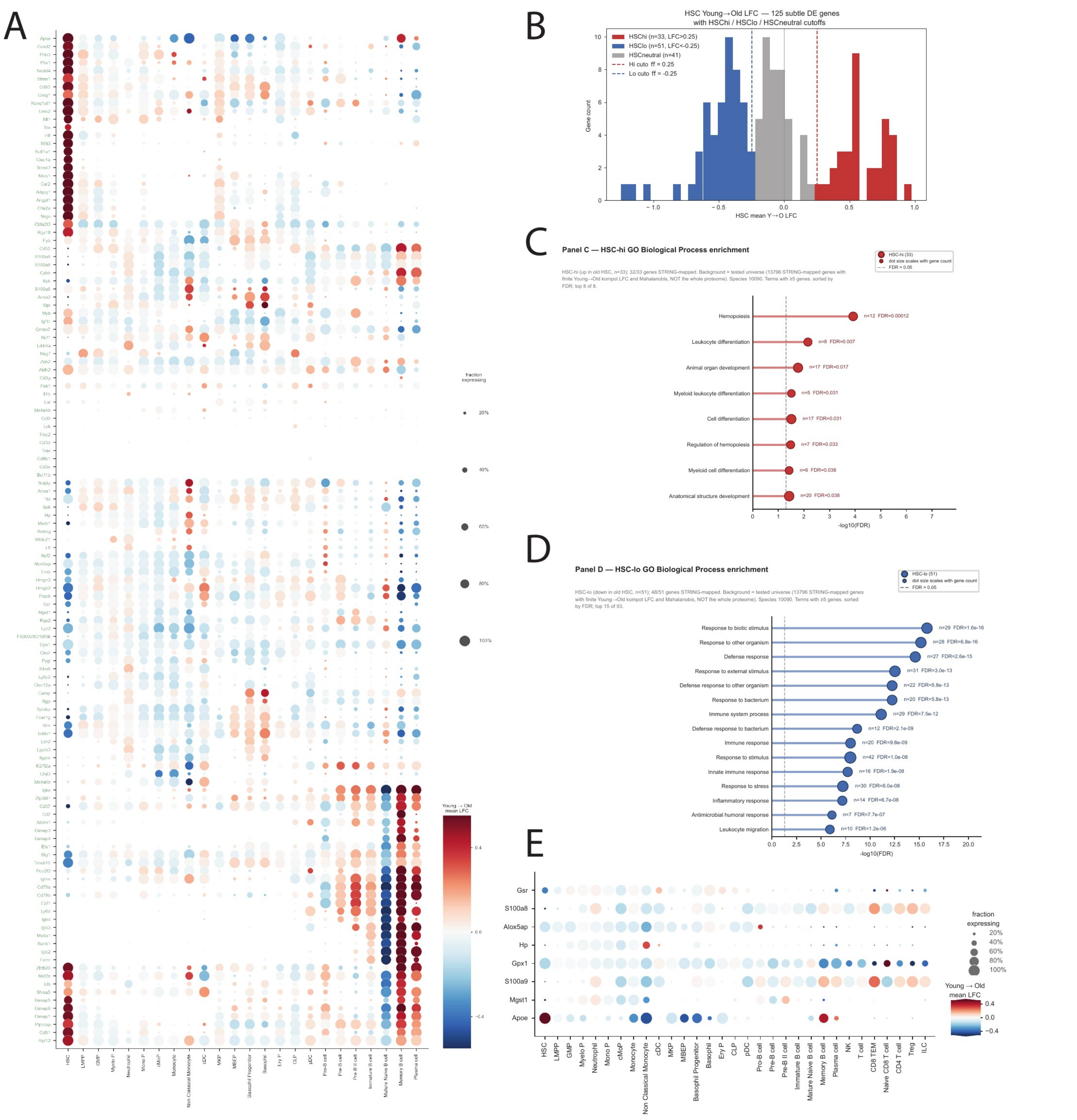

**Supplementary Figure 11: Dissection of the subtle aging signal across the hematopoietic landscape.**

(A) Per-cell-type Young-to-Old mean log fold change dot plot for 125 subtle differentially expressed genes from **Fig. 3D**; dot size encodes the fraction of cells expressing the gene and color represents the mean log fold change.

(B) The subtle-class genes from (A) grouped by their HSC fold change into $HSC^{hi}$, $HSC^{neutral}$ and $HSC^{lo}$ sets.

(C) Gene-ontology Biological Process enrichment of the $HSC^{hi}$ set.

(D) Same as (C), for the $HSC^{lo}$ set.

(E) Per-cell-type Young-to-Old mean log fold change dot plot for the eight antioxidant-activity genes (GO:0016209) driving the GO Molecular Function enrichment of **Fig. 3G bottom**. Dot size encodes the fraction of cells expressing the gene.

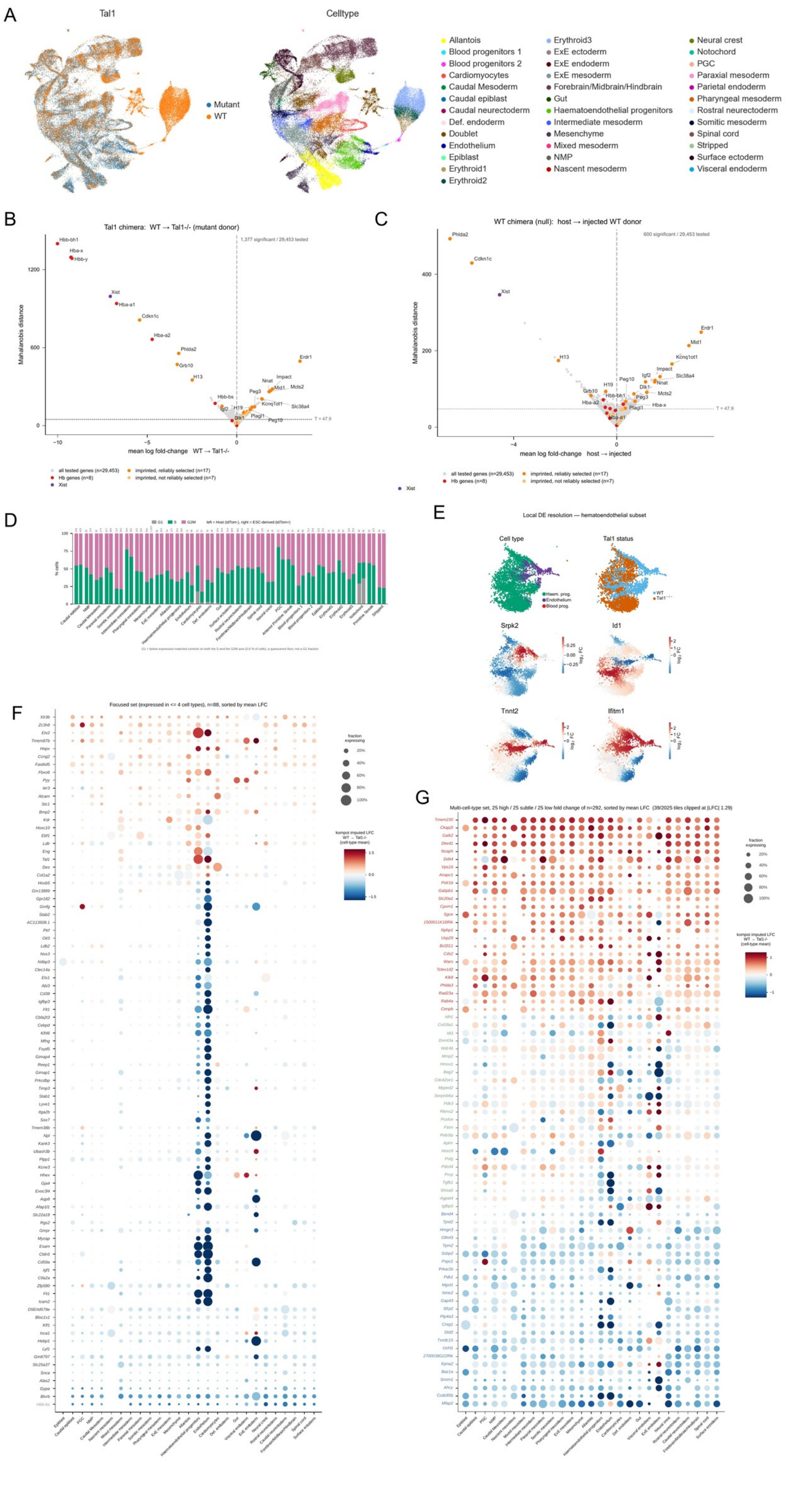

**Supplementary Figure 12: Chimera-artefact correction and breadth-resolved differential expression in the *Tal1*$^{-/-}$ embryo**

(A) UMAPs of the *Tal1*$^{-/-}$ chimera, colored by *Tal1* status and by cell type, before the depleted-lineage exclusion. The erythroid and blood-progenitor compartments are almost entirely wild-type and are the lineages excluded before testing.

(B) Quasi volcano of the uncorrected wild-type $\rightarrow$ *Tal1*$^{-/-}$ comparison (mean log fold change against Mahalanobis distance), with hemoglobin paralogues, *Xist* and imprinted genes highlighted (1,377 significant of 29,453 tested).

(C) Same as (B), for the WT-only control chimera (600 significant of 29,453 tested).

(D) Cell-cycle phase composition per cell type in the WT-only chimera, left host (tdTomato$^{-}$) and right ESCderived (tdTomato$^{+}$), drawn and scored as in **Fig. 4I**.

(E) Differential expression within the hematoendothelial subset: the subset colored by cell type and by *Tal1* status, and per-cell log fold change for *Srpk2*, *Id1*, *Tnnt2* and *Ifitm1*.

(F) Per-cell-type dot plot of the focused set ($n = 88$, expressed in $\leq 4$ cell types), sorted by mean log fold change; dot size encodes the fraction of cells expressing the gene and color the mean WT $\rightarrow$ *Tal1*$^{-/-}$ log fold change.

(G) Per-cell-type dot plot of the multi-cell-type set (expressed in 5–24 cell types), drawn as in (F), showing 75 of the $n = 292$ genes: the 25 with the highest, 25 with the lowest and 25 with the most subtle mean log fold change, sorted by mean log fold change. 39 of 2,025 tiles are clipped at $|\mathrm{LFC}| = 1.29$.

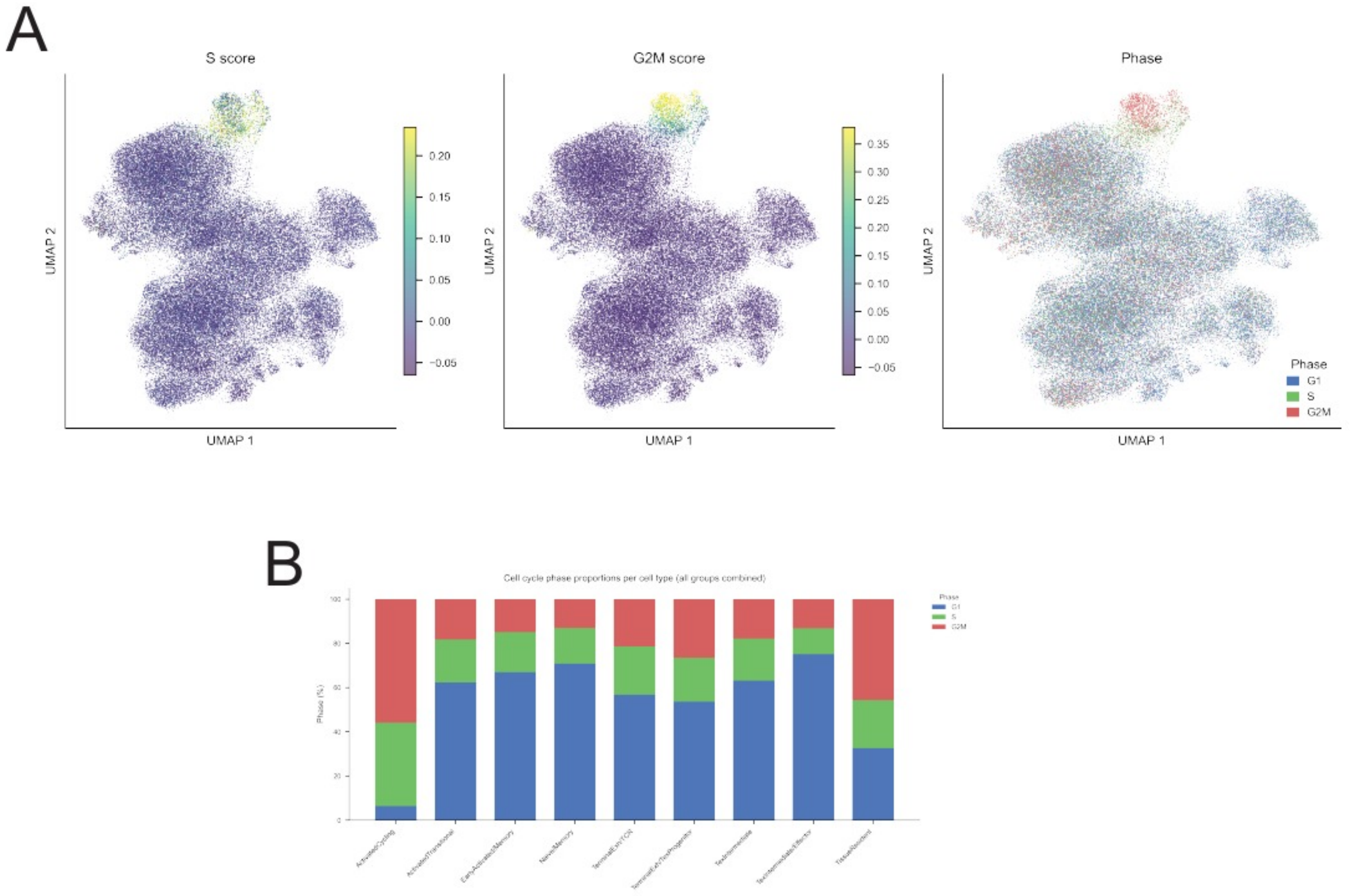

**Supplementary Figure 13: CD8 T cell compartment in the melanoma immunotherapy analysis.**

(A) UMAP of the CD8 T cell compartment colored by S score, by G2M score, and by the resulting cell-cycle phase call

(B) Cell-cycle phase proportions per CD8 T cell state, all treatment arms combined.

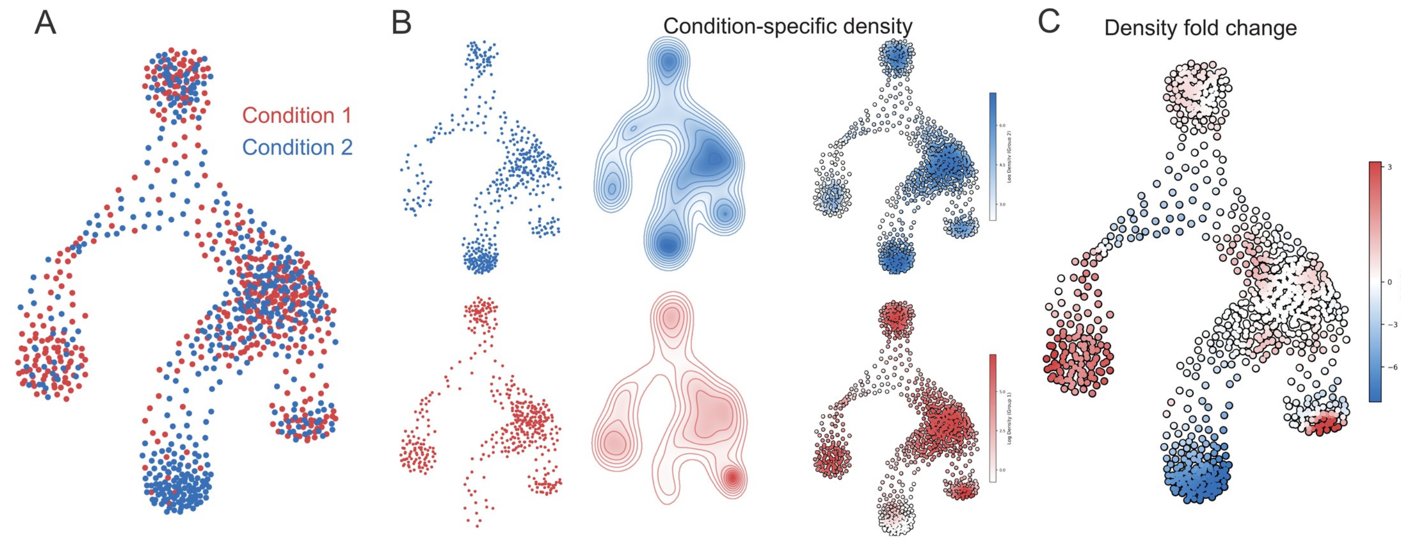

**Supplementary Figure 14: Schematic of differential density estimation.**

(A) The co-embedded cell states of the two conditions, as in the schematic of **Fig. 1A**.

(B) One row per condition, in three columns: the cell states of that condition alone; the density function inferred from them; and that condition's log density evaluated at every cell state, including those observed in the other condition.

(C) The log fold change in density between the two conditions, across all cell states.

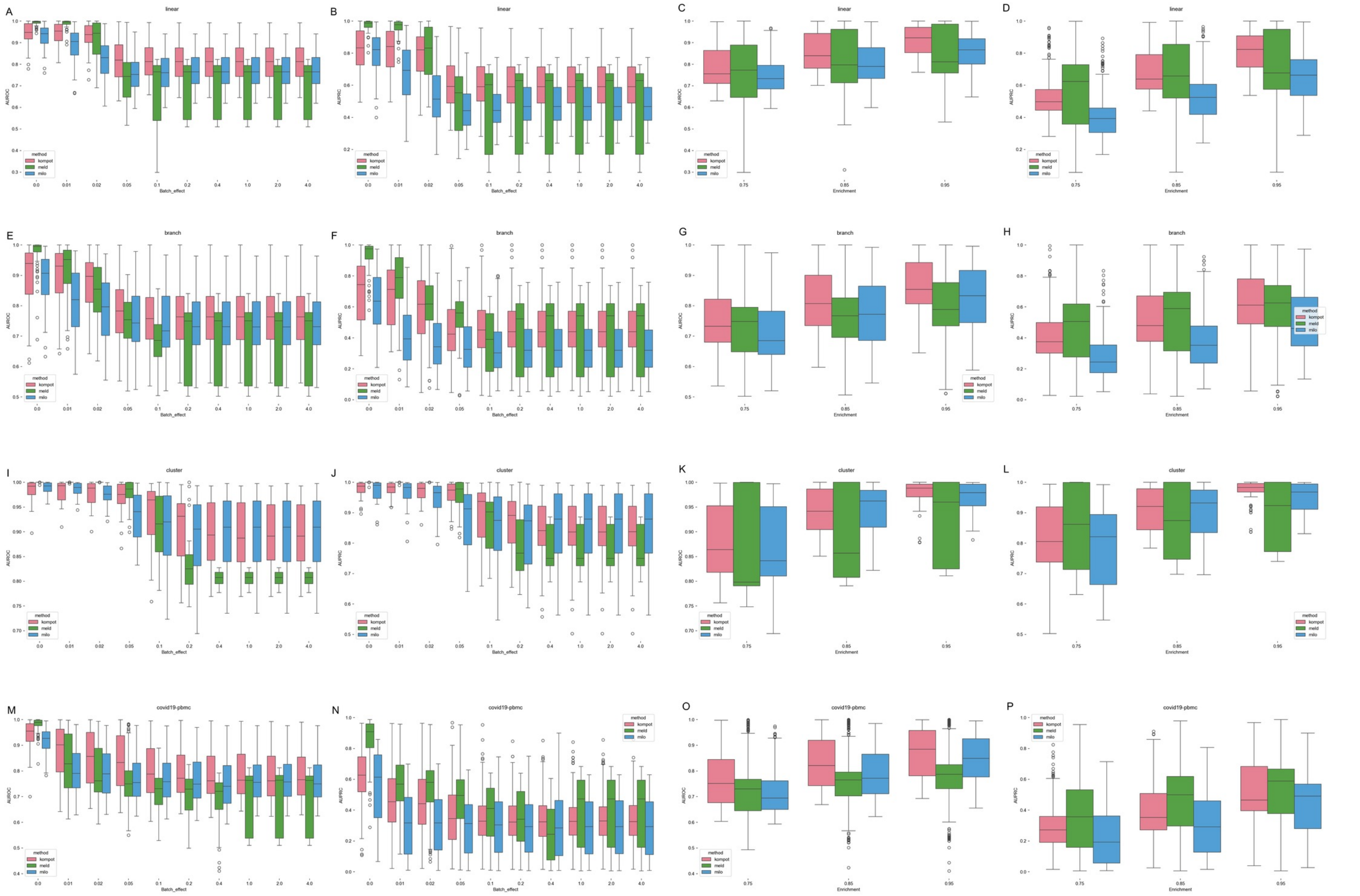

**Supplementary Figure 15: Benchmarking of differential abundance analysis.**

Benchmarking was performed the framework defined in benchmarkDA [1]. Three simulated datasets (linear, branch and cluster) and one real-world datasets with artificially generated labels (covid-19) were used. Kompot performed was compared against Meld [2] and Milo [3]. AUROC and AUPRC were used for performance comparison.

(A–B) AUROC (A) and AUPRC (B) for Kompot, Meld and Milo for the linear dataset with different batch effect levels. Statistics were computed for different resolutions and cell types.

(C–D) AUROC (C) and AUPRC (D) for Kompot, Meld and Milo for the linear dataset with resolutions. Statistics were computed for different batch effects and cell types.

(E–H) Same as (A–D) for the branch dataset.

(I–L) Same as (A–D) for the cluster dataset.

(M–P) Same as (A–D) for the covid-19 dataset.

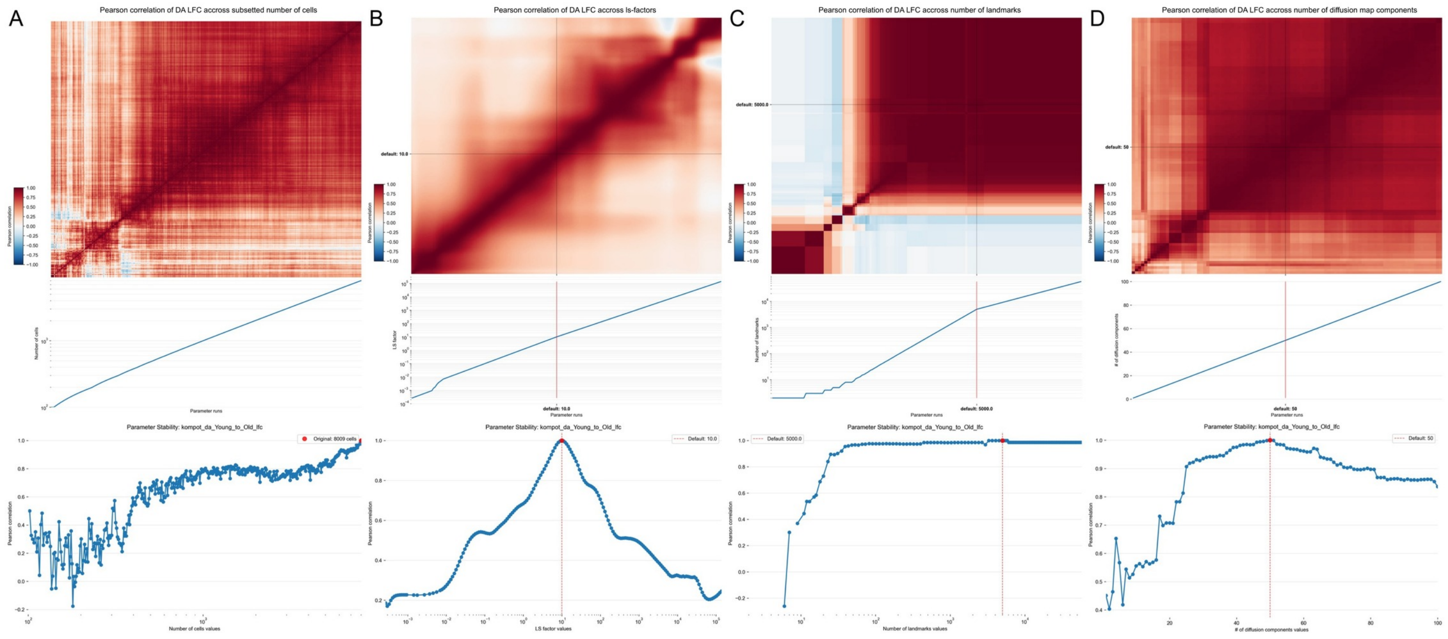

**Supplementary Figure 16: Robustness of Kompot differential abundance mean log fold change for the hematopoietic aging dataset**

Benchmarking was performed by computing the Pearson correlation of differential abundance log fold change across all cell-states between pairs of parameter choices.

Top: Heatmaps shows Pearson correlations pairs of parameter values, computed for the Young vs. Old comparison in the aging dataset. Each column shows a different parameter being varied: (A) number of cells (dataset subset to varying number cells), (B) length-scale factor (default: 10), (C) number of landmarks (default: 5000), and (D) number of diffusion map components (default: 50). Default values are indicated by black lines on the matrix

Middle: Margin plots mapping each column of the correlation matrix to its corresponding parameter value for the respective datasets(note log scaling for some parameters).

Bottom: Correlation of abundance LFC to a reference setting as a function of the varied parameter. The reference is the maximum cell count in (A) and the default parameter value in (B–D), indicated by a red dashed line. This corresponds to extracting a single row from the heatmap in (A).

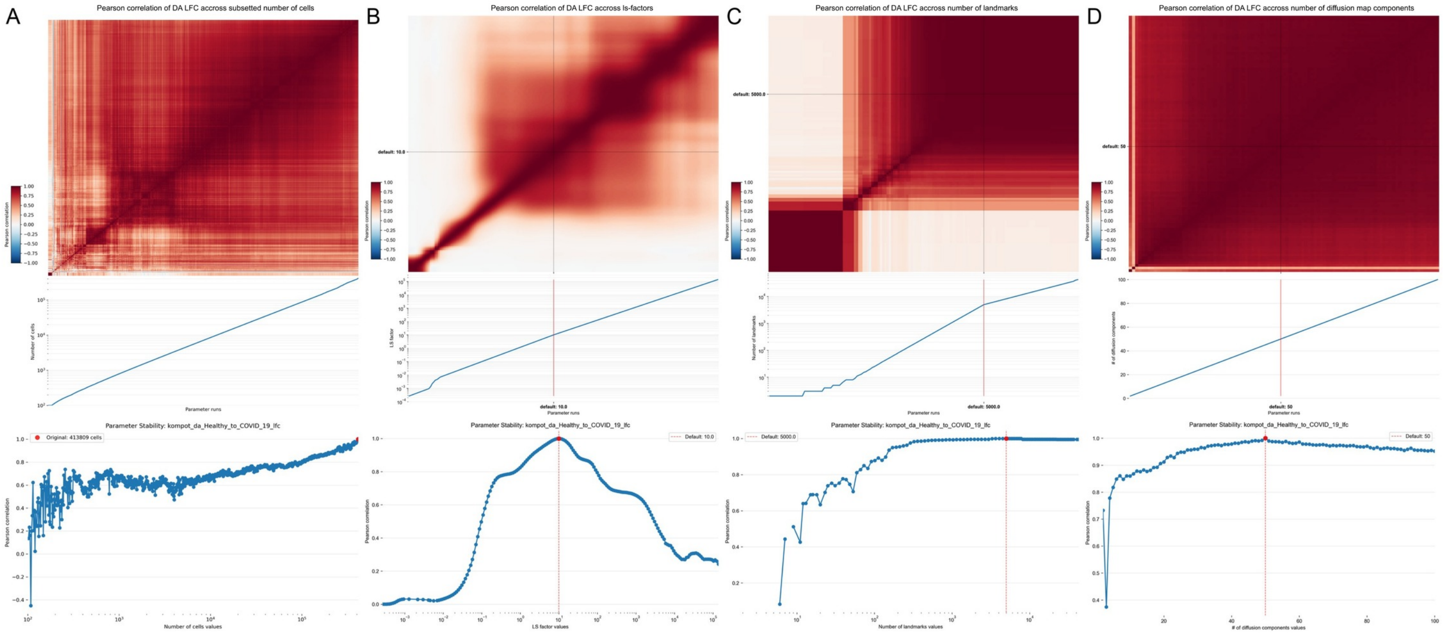

**Supplementary Figure 17: Robustness of Kompot differential abundance mean log fold change for the COVID-19 dataset**

(A–D) Same as Supplementary Fig. 16 for the COVID-19 dataset.

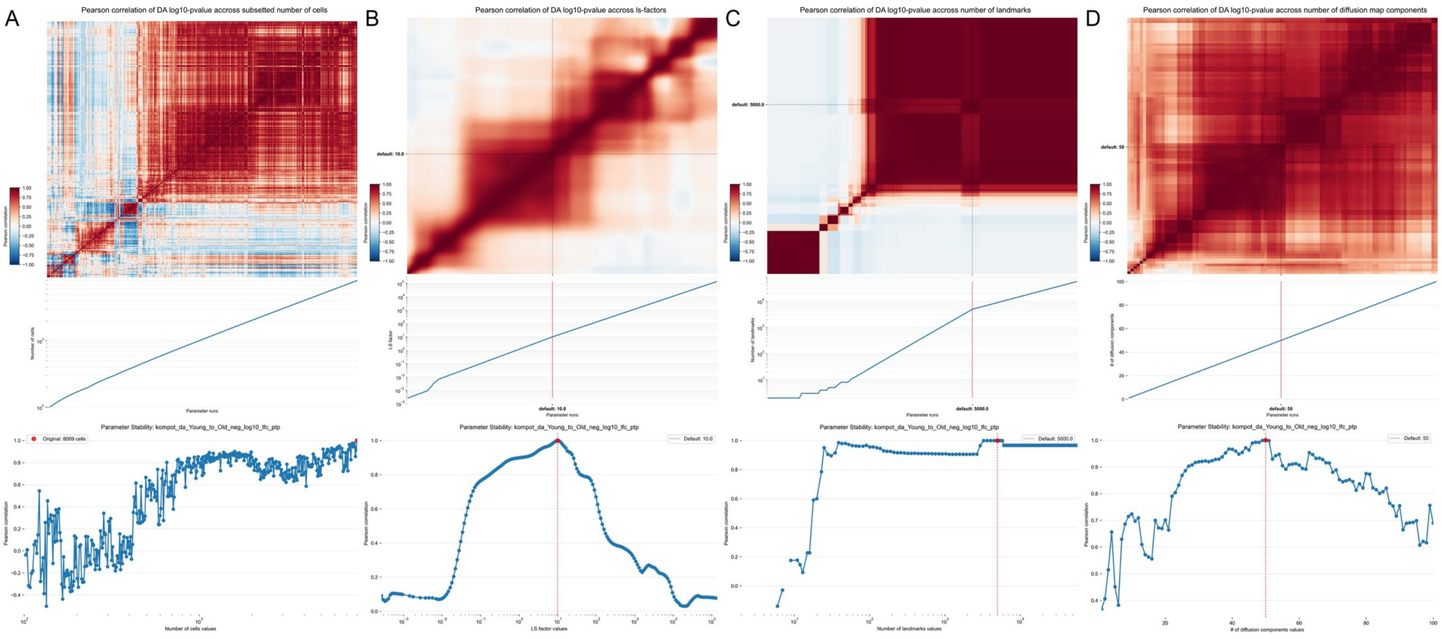

**Supplementary Figure 18: Robustness of Kompot differential abundance posterior tail probabilities for the hematopoietic aging dataset**

(A–D) Same as Supplementary Fig. 16 showing robustness of posterior tail probabilities instead of log fold change.

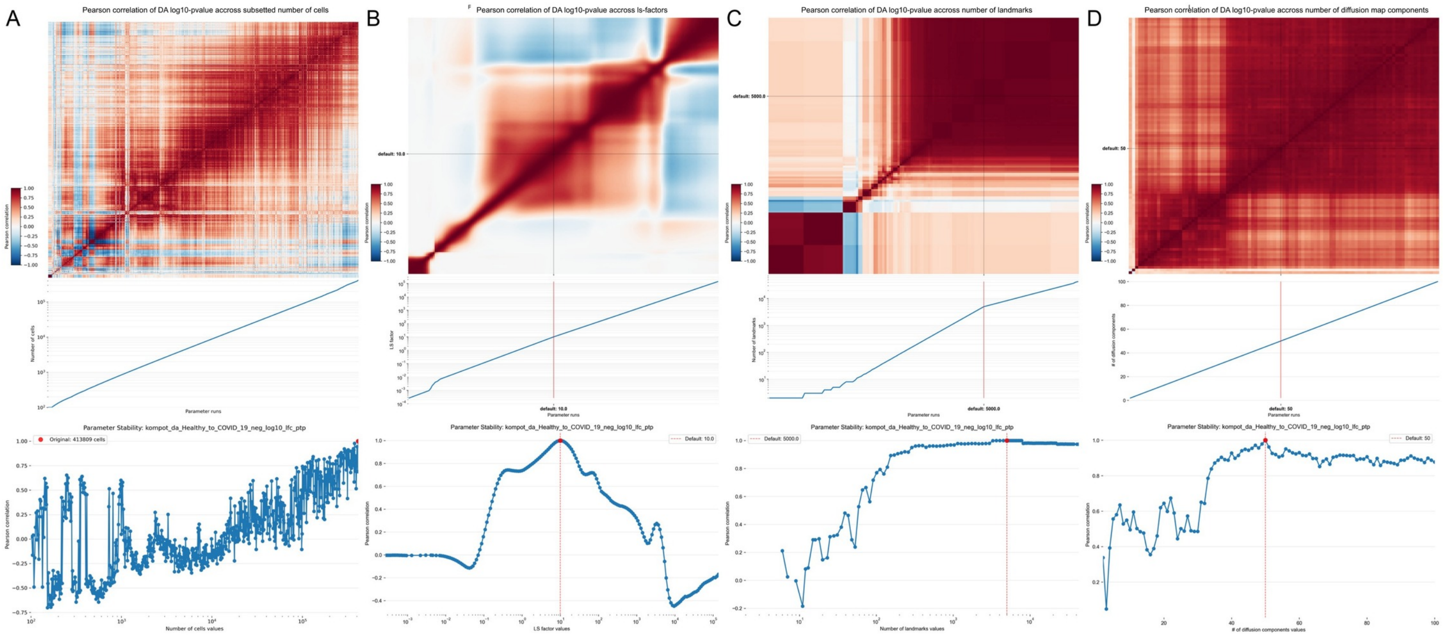

**Supplementary Figure 19: Robustness of Kompot differential abundance posterior tail probabilities for the COVID-19 dataset**

(A–D) Same as Supplementary Fig. 18 for the COVID-19 dataset.

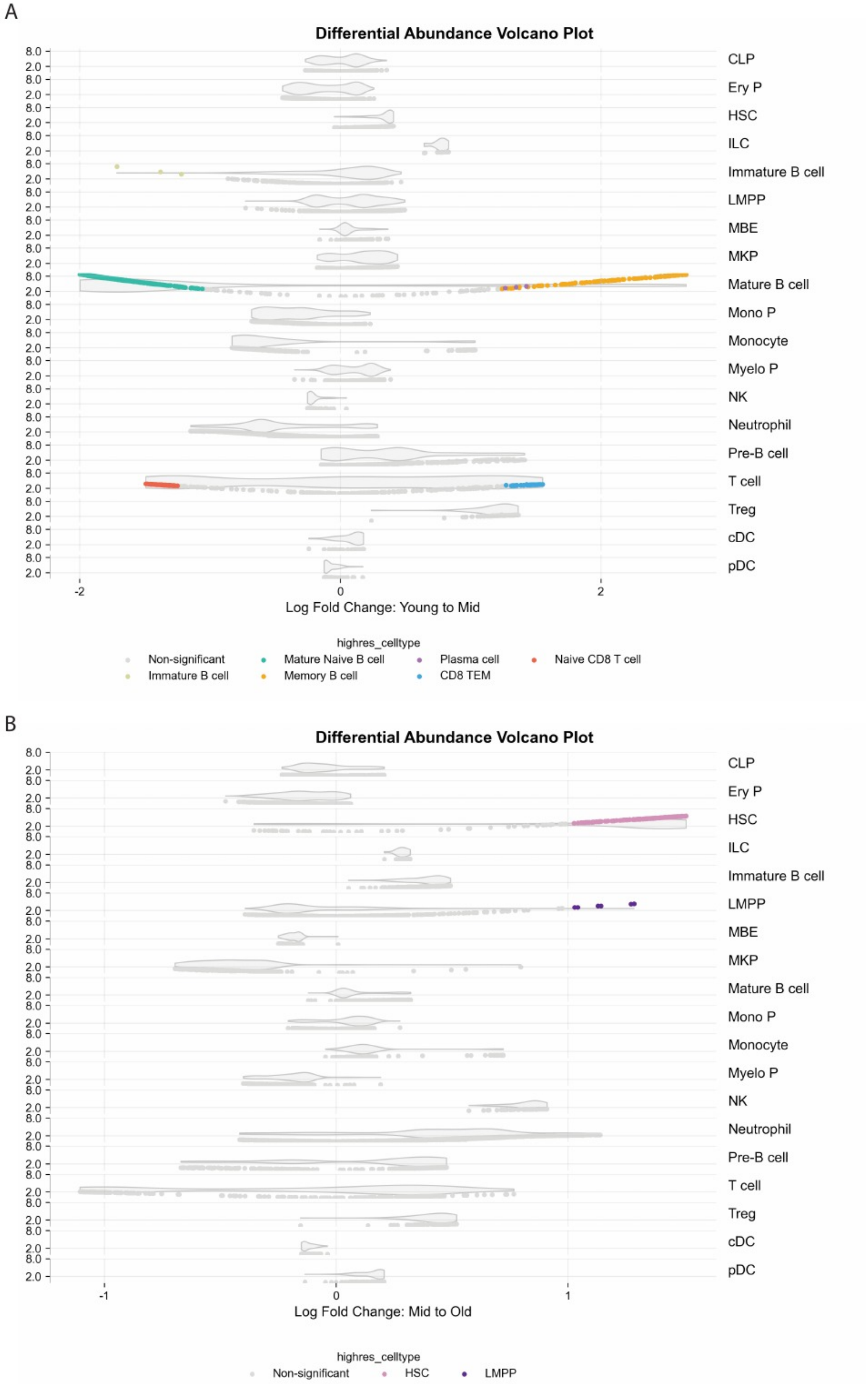

**Supplementary Figure 20: Kompot differential abundance analysis in aging hematopoiesis.** Same as **Fig. 6F**, for Young to Mid and Mid to Old comparisons.

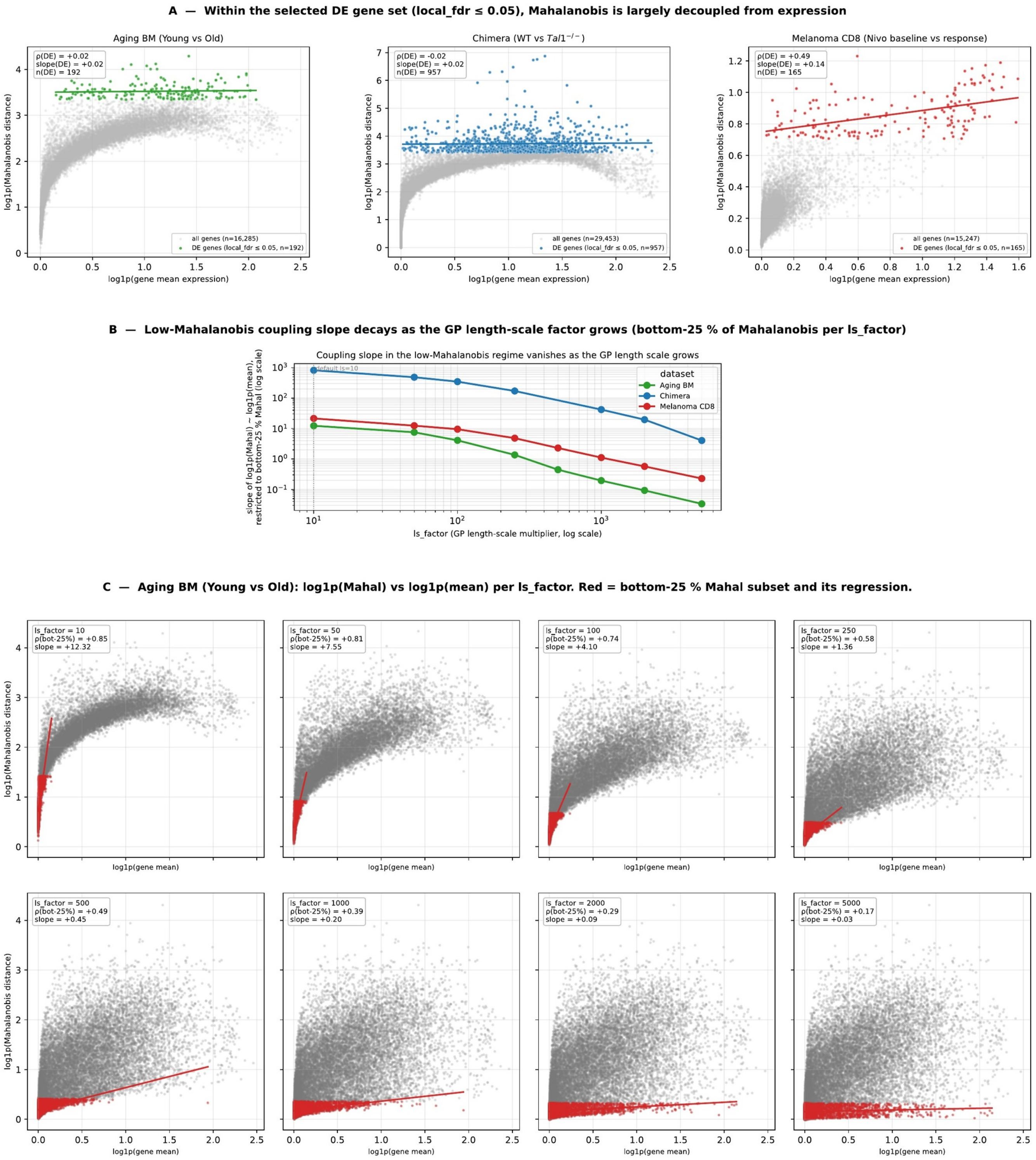

**Supplementary Figure 21: Coupling between Mahalanobis distance and gene expression level.**

(A) Per-dataset scatter of $\log_{1p}$(Mahalanobis) against $\log_{1p}$(gene mean), restricted to genes passing the reported significance gate (local FDR $\leq 0.05$), with regression line and Spearman $\rho$; within the selected set the coupling is absent.

(B) Slope of $\log_{1p}$(Mahalanobis) against $\log_{1p}$(mean) restricted to the bottom quartile of Mahalanobis values, as a function of the GP length-scale factor, for all three datasets on log-log axes; the low-distance coupling collapses as the length scale grows.

(C) Small multiples for the aging bone marrow dataset showing the full-gene scatter at length-scale factors 10, 50, 100, 250 and 500, with the bottom-quartile regression line in red.

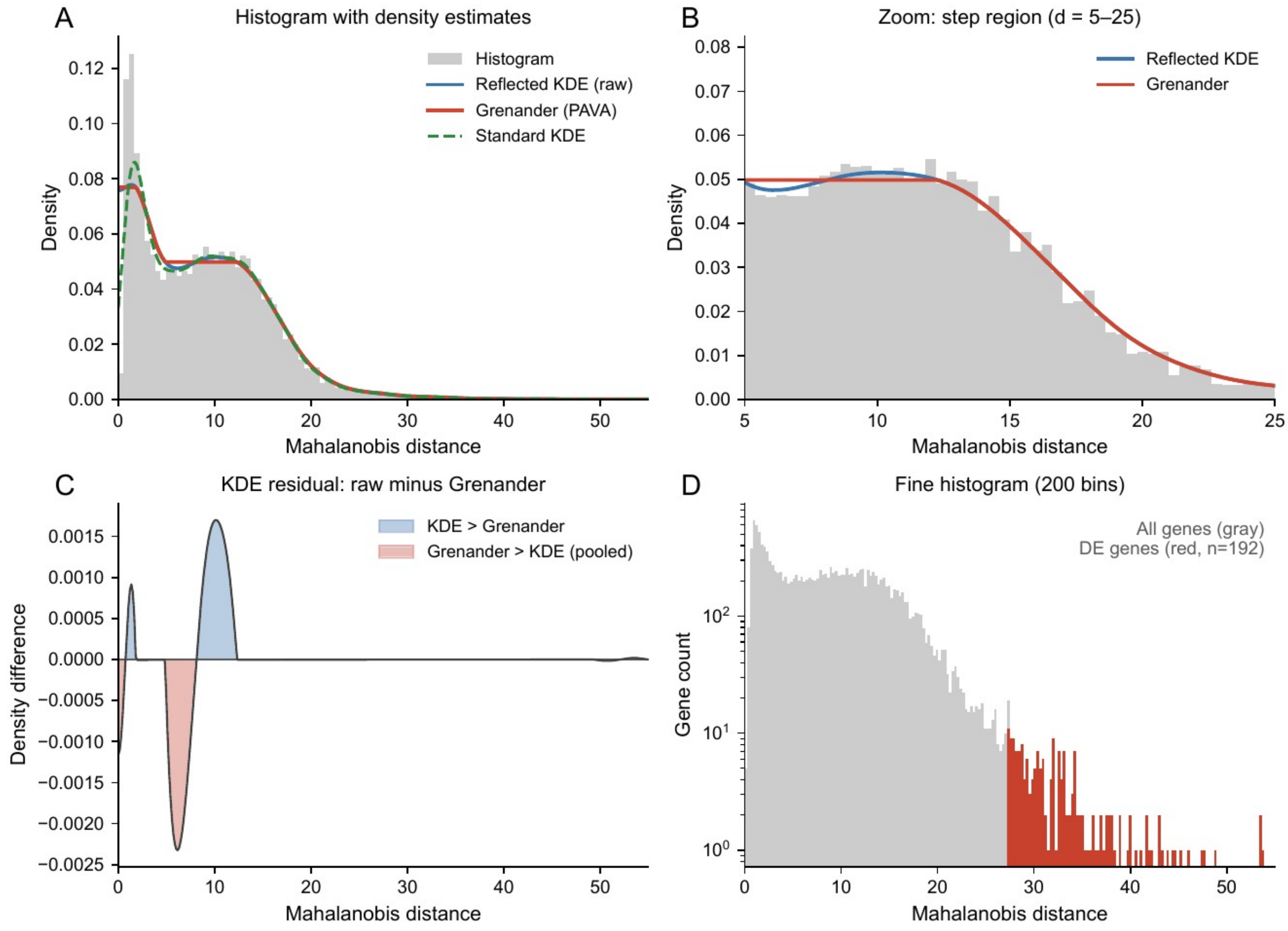

**Supplementary Figure 22: Null-density estimator behaviour on real Mahalanobis distances.**

Diagnostic of the gene-shuffling-null density estimator that converts Mahalanobis distances to local FDR, evaluated on the young-to-old comparison of our aging atlas ($n = 192$ DE genes at the calibrated local-FDR cutoff).

(A) Mahalanobis-distance histogram (gray) with three density estimators overlaid: Reflected KDE (blue), Grenander PAVA monotone estimator (orange-red), and Standard KDE (green dashed). The Grenander PAVA estimator is what Kompot uses for the null mass; it enforces monotonicity in the right tail without rounding off the step-shaped shoulder near $d \approx 5$–$12$ that the standard KDE smooths through.

(B) Zoom of the step region ($d = 5$–$25$) showing that the reflected-KDE and Grenander curves are visually indistinguishable in this range.

(C) Density residual (raw reflected-KDE minus Grenander): the standard KDE puts excess mass above Grenander between $d \approx 10$–$13$ (blue lobe) and deficit mass between $d \approx 4$–$8$ (red lobe), confirming that the only material disagreement between the two estimators is in the bulk shoulder, not the tail that drives FDR.

(D) Fine 200-bin histogram on log-count axis, separating all genes (gray) from DE genes (red, $n = 192$); the DE bucket cleanly separates from the null bulk above $d \approx 27$.
