## Supplementary Notes for "Comparing phenotypic manifolds with Kompot: Cluster-free differential expression at single-cell resolution"

### Supplementary Note 1

### Gaussian Processes and Posterior Uncertainty

In this section, we discuss the different sources of uncertainty for the functions we derive through a Gaussian Process (GP). We use GPs to compute feature-expression functions, which predict gene expression or another feature at arbitrary points in the cell-state space, and to infer cell-state density functions through *Mellon*. In both cases, we employ Bayesian inference to compute a posterior distribution of functions that quantifies our uncertainty about the *true* gene-expression or cell-state density function. Different sources of uncertainty are considered, including those arising from latent representations, measurement noise, and approximation techniques.

#### 1.1 Theoretical Framework

Gaussian Processes (GPs) define a distribution over functions, characterized by a mean function $m(x)$ and a covariance function $k(x, x')$. For a set of observed data points $\mathcal{D} = \{(x_i, y_i)\}_{i=1}^N$, the GP posterior distribution is given by:

$$f(x) \mid \mathcal{D} \sim \mathcal{GP}\left(m_{\text{post}}(x),\, k_{\text{post}}(x, x')\right).$$

The posterior mean and covariance functions are computed as:

$$m_{\text{post}}(x) = m(x) + K_{x,X}\, K_{X,X}^{-1}\, (Y - m(X))\,, \tag{1.1}$$

$$k_{\text{post}}(x, x') = k(x, x') - K_{x,X}\, K_{X,X}^{-1}\, K_{X,x'}. \tag{1.2}$$

- $K_{X,X}$ is the covariance matrix for the observed inputs $X$.
- $K_{x,X}$ is the covariance vector between a test point $x$ and the observed inputs.
- $Y$ is the vector of observed outputs.

The covariance matrices are computed using a kernel function $K_{x,x'} = k(x, x')$ that introduces a prior statistical connection between function values based on the similarity of cell states. The details of this choice are discussed in Section 1.1.1.

##### 1.1.1 Covariance

The prior covariance kernel encodes our assumptions about the smoothness and similarity of function values at different input locations. In our framework, the kernel $k(x, x')$ defines a statistical relationship between the function values $f(x)$ and $f(x')$ according to the distance or similarity between the corresponding cell states $x$ and $x'$. This prior correlation structure is critical for capturing the underlying biological relationships in cell-state space.

While our implementation permits the free choice of covariance functions, we, by default, employ a Matérn $\frac{5}{2}$ kernel. This kernel is widely favored over alternatives such as the squared exponential or Gaussian kernel because it allows for a controlled level of roughness in the modeled functions, reflecting the moderate smoothness observed in many biological systems. The Matérn $\frac{5}{2}$ kernel strikes a good balance between flexibility and smoothness, and its well-defined derivative structure makes it particularly useful for applications that require differentiability. Importantly, choosing a less restrictive kernel over the infinitely smooth squared exponential is critical for uncertainty estimation: assuming an overly smooth gene expression function can unnaturally reduce the resulting posterior uncertainty, leading to overconfident inferences [1, 2].

A further advantage of our approach is the incorporation of a heuristic for selecting the length scale parameter of the Matérn $\frac{5}{2}$ covariance function. In the *Mellon* manuscript [3], we demonstrated that the results are remarkably stable with respect to variations in the length scale. The heuristic consistently chooses length scales that are close to the optimal values, thereby preserving the quality of the inferences. Importantly, this heuristic is the primary factor that accelerates computations, making the approach feasible for datasets comprising millions of cells, since exhaustive hyperparameter tuning would be computationally prohibitive.

##### 1.1.2 The Sparse GP

Sparse Gaussian Processes (SGPs) address the computational limitations of standard GPs when dealing with large datasets. In SGPs, the full set of observations $\mathcal{D}$ is approximated using a smaller set of $M$ inducing points $Z = \{z_j\}_{j=1}^M$, where $M \ll N$. The key approximation is to assume that the function values $f(X)$ are conditionally independent given $f(Z)$. The joint distribution over $f(X)$ and $f(Z)$ is given by:

$$\begin{bmatrix} f(Z) \\ f(X) \end{bmatrix} \sim \mathcal{N}\left( \begin{bmatrix} m(Z) \\ m(X) \end{bmatrix}, \begin{bmatrix} K_{Z,Z} & K_{Z,X} \\ K_{X,Z} & K_{X,X} \end{bmatrix} \right).$$

From this joint distribution, the conditional distribution of $f(X)$ given $f(Z)$ is:

$$f(X) \mid f(Z) \sim \mathcal{N}\left(\mu_{X|Z},\, \Sigma_{X|Z}\right),$$

where:

$$\mu_{X|Z} = m(X) + K_{X,Z}\,K_{Z,Z}^{-1}\,(f(Z) - m(Z))\,, \tag{1.3}$$
$$\Sigma_{X|Z} = K_{X,X} - K_{X,Z}\,K_{Z,Z}^{-1}\,K_{Z,X}. \tag{1.4}$$

Using this result, the posterior mean and covariance functions of the sparse GP are approximated as:

$$m_{\text{post}}(x) = m(x) + K_{x,Z}\,K_{Z,Z}^{-1}\,K_{Z,X}\,K_{X,X}^{-1}\,(Y - m(X))\,, \tag{1.5}$$
$$\begin{aligned} k_{\text{post}}(x,x') &= k(x,x') - K_{x,Z}\,K_{Z,Z}^{-1}\,K_{Z,x'} \\ &\quad + K_{x,Z}\,K_{Z,Z}^{-1}\,K_{Z,X}\,K_{X,X}^{-1}\,K_{X,Z}\,K_{Z,Z}^{-1}\,K_{Z,x'}. \end{aligned} \tag{1.6}$$

The use of $K_{Z,Z}$, the covariance matrix over inducing points, significantly reduces the computational cost of inversion from $\mathcal{O}(N^3)$ to $\mathcal{O}(NM^2)$. The sparse-GP approximation in Eqs. 1.5–1.6, including the Nyström-style inducing-point construction and the Cholesky-based posterior sampler, is implemented in *Mellon* [3]; Kompot consumes Mellon's posterior mean and covariance through its predictor interface rather than re-deriving these formulas internally.

##### 1.1.3 Latent Representation and Cholesky Decomposition

When inferring the cell-state density function with *Mellon*, we cannot directly condition the posterior on preexisting data due to the absence of explicit observations. Instead, we impose a Gaussian prior on a latent representation $z \in \mathbb{R}^M$ and link the function values to a nearest-neighbor distribution. As detailed in the *Mellon* manuscript [3], this formulation allows us to define the prior in terms of both the latent space and the function's smoothness properties:

$$z \sim \mathcal{N}(0, \mathbb{I}), \tag{1.7}$$
$$f(X) = L_X\,z + m(X), \tag{1.8}$$
$$L_X L_X^T = K_{X,Z}\,K_{Z,Z}^{-1}\,K_{Z,X} \approx K_{X,X}, \tag{1.9}$$
$$f(X) \sim \mathcal{N}\left(m(X),\, K_{X,X}\right). \tag{1.10}$$

Thus, instead of conditioning directly on existing data $Y$, we link the resulting function values to our data stochastically via $f(X) \sim g(Y)$ and perform Bayesian inference on $z$. The approximation $L_X L_X^T \approx K_{X,X}$ is achieved by leveraging the inducing points $Z$ to efficiently represent the covariance structure, reducing the computational complexity associated with directly inverting $K_{X,X}$ and instead applying the Cholesky decomposition on the low-dimensional $K_{Z,Z}$. Expressing $f(X)$ in terms of $z$ and $L_X$ enables efficient posterior inference while respecting the underlying structure of the cell-state space. However, it also introduces an additional source of uncertainty that must be propagated from the posterior distribution of $z$.

#### 1.2 Sources of Uncertainty

##### 1.2.1 Latent Representations

For the cell-state density inference, we use a latent representation of the inferred function as explained in 1.1.3. The latent parameters are optimized to their maximum a posteriori (MAP) estimate, and posterior uncertainty is obtained from a Laplace approximation [1], which models the posterior as a Gaussian centred at the mode with covariance equal to the inverse Hessian of the negative log-posterior. We use the diagonal of that Hessian, so the resulting posterior over the latent representation is again parameterized by a mean and a per-parameter variance:

$$z \sim \mathcal{N}\left(\mu_{\text{post}},\, \text{diag}(\sigma^2_{\text{post}})\right). \tag{1.11}$$

This propagates into a posterior uncertainty of the mean function from (1.8):

$$m_{\text{post}}(x) \sim \mathcal{N}\left(L_x\,z + m(x),\, L_x\,\text{diag}(\sigma^2_{\text{post}})\,L_x^T\right). \tag{1.12}$$

Note that this is the posterior for the mean function at any cell state $x$. The posterior distribution for the actual function value includes additional terms (see Section 1.3.2).

##### 1.2.2 Measurement Noise

Feature–expression functions are conditioned on observed feature values $y_i$. While raw molecular counts are better described by discrete distributions such as the negative binomial, after normalization and log-transformation we approximate measurement noise as Gaussian with constant variance $\sigma^2$. This homoscedastic assumption does not account for overdispersion (variance-mean relationships). A fuller discussion of this limitation and potential solutions is provided in Supplementary Note 5. This Gaussian approximation is essential in the Gaussian Process framework, as it enables a closed-form posterior. We therefore model noisy observations as

$$y_i = f(x_i) + \epsilon, \quad \epsilon \sim \mathcal{N}(0, \sigma^2),$$

where $f(x_i)$ is the true underlying function. The covariance function incorporates this noise term by modifying $K_{X,X}$ as

$$K_{X,X} \to K_{X,X} + \sigma^2 I.$$

##### 1.2.3 Sparse GP Approximation

To handle large datasets, we employ sparse GPs using inducing points $Z = \{z_j\}_{j=1}^M$, where $M \ll N$. The posterior distribution is defined in Section 1.1.2 with equation (1.6) for $k_{\text{post}}(x,x')$ featuring $K_{X,X}$. Note that in the case of cell-state density inference, we do not add noise to $K_{X,X}$ since we condition on the posterior mean function $f(X)$.

#### 1.3 Applications

##### 1.3.1 Feature-Expression Prediction

Unlike the density case of Section 1.1.3, where the function is inferred through a latent representation and the posterior must be obtained for that representation first, feature values are observed. The Gaussian process is therefore conditioned on the measurements themselves and the posterior is the standard Gaussian process conditional, with no latent layer in between. However, we must account for measurement noise by modifying $K_{X,X}$ as described in Section 1.2.2. The posterior distribution for function values at cell states $\xi = \{x_i\}_{i=1}^{m}$ is:

$$f(\xi) \sim \mathcal{N}\left(m_{\text{post}}(\xi),\, k_{\text{post}}(\xi)\right). \tag{1.13}$$

Hence, both the posterior mean (Eq. 1.1) and posterior covariance (Eq. 1.2), as well as their sparse counterparts (Eqs. 1.5–1.6), are available in closed form. This closed-form structure allows us to compute the posterior directly without the typical expense of sampling-based methods or optimization required in variational inference.

For efficient numerical computation, we avoid explicitly inverting large covariance matrices such as $K_{X,X}$. Instead, we apply a Cholesky decomposition

$$K_{X,X} + \sigma^2 I = L\,L^T,$$

with $L$ a lower triangular matrix. Posterior quantities are then computed via forward and backward triangular solves, which are both numerically stable and computationally efficient, allowing our framework to scale to large single-cell datasets.

##### 1.3.2 Cell-State Density Functions

Cell-state density is inferred through a latent representation, so its posterior combines two sources of uncertainty. For cell-state density inference, we combine the uncertainties of the posterior mean function $m_{\text{post}}$ with the uncertainty of the function value given by $k_{\text{post}}$. This includes the variance of the function values at any new cell state $x_i$ as well as the covariance between any pair of states in $\xi = \{x_i\}_{i=1}^{m}$:

$$f(\xi) \sim \mathcal{N}\left(L_\xi\, z + m(\xi),\, L_\xi \;\text{diag}(\sigma^2_{\text{post}})\, L_\xi^T + k_{\text{post}}(\xi)\right). \tag{1.14}$$

Conceptually, this is the result of stacking two inferences. First, we infer the latent representation of the function (subject to the posterior uncertainty in (1.12)). Then, we compute the function values at new cell states $x_i$, applying the additional uncertainty of the GP through $k_{\text{post}}$. Hence, $f(\xi)$ can be written as the sum of two random variables:

$$f(\xi) = a(\xi) + b(\xi), \tag{1.15}$$

$$a(\xi) \sim \mathcal{N}\left(0,\, k_{\text{post}}(\xi)\right), \tag{1.16}$$

$$b(\xi) \sim \mathcal{N}\left(L_\xi\, z + m(\xi),\, L_\xi \;\text{diag}(\sigma^2_{\text{post}})\, L_\xi^T\right). \tag{1.17}$$

Here, $a(\xi)$ represents the GP's posterior uncertainty about deviations from zero, and the distribution of $b(\xi) = m_{\text{post}}(\xi)$ represents the uncertainty in the inferred mean function.

### Supplementary Note 2

### Mahalanobis Distance

We compute functions over the cell-state space using the Gaussian Process framework discussed in previous sections. The function of primary interest is gene expression: for each gene, a GP maps cell state to expression levels, and the comparison of those functions between conditions is the differential-expression test. The same construction applies to cell-state density, which yields the differential-abundance test, and to other phenotypic features that vary across cellular states. When single-cell datasets from multiple experimental conditions are available, and certain cell states are well aligned across conditions, separate GP functions can be inferred for each condition, conditioned on the respective data. This allows a quantitative comparison of any such feature between conditions. Nothing in the derivation that follows depends on which feature is chosen: the only feature-specific ingredient is the posterior covariance, whose origin in each case is given at the end of this section.

A key advantage of the GP framework is its ability to compute the posterior mean value of the compared function within a given condition at any cell state $m_{\text{post}}(x)$, which is our best estimate even where no cell was observed. The associated uncertainty is captured by the posterior covariance, as discussed in Section 1.2.

#### 2.1 Definition

The Mahalanobis distance quantifies the deviation of a point from a multivariate normal distribution while accounting for variances and covariances among dimensions. Given a point $x$ and a normal distribution with mean 0 and covariance matrix $\Sigma$, the Mahalanobis distance is defined as:

$$D(x) = \sqrt{x^T \, \Sigma^{-1} \, x}. \tag{2.1}$$

For our analysis, we compare two experimental conditions, $a$ and $b$, where the compared functions $f_a$ and $f_b$ (for differential expression, the expression functions of a given gene) are inferred as independent normal distributions. The covariance matrices $\Sigma_a$ and $\Sigma_b$ quantify both our uncertainty about the true function values and the statistical interdependencies emerging from our inference process:

$$f_a \sim \mathcal{N}\left(\mu_a,\, \Sigma_a\right), \tag{2.2}$$
$$f_b \sim \mathcal{N}\left(\mu_b,\, \Sigma_b\right). \tag{2.3}$$

Thus, the difference between the two functions is also normally distributed:

$$f_a - f_b \sim \mathcal{N}\left(\mu_a - \mu_b,\, \Sigma_a + \Sigma_b\right). \tag{2.4}$$

We seek to quantify the likelihood that the true difference is effectively zero. To do so, we compute the Mahalanobis distance of this difference from zero:

$$D(a, b) = \sqrt{\left(\mu_a - \mu_b\right)^T \, \left(\Sigma_a + \Sigma_b\right)^{-1} \, \left(\mu_a - \mu_b\right)}. \tag{2.5}$$

This measure quantifies the significance of the difference while accounting for uncertainty in both conditions.

#### 2.2 Interpretation

When (2.5) is computed for a single dimension (comparing function values at a single cell state), it reduces to a standard z-score. However, when additional cell states are included, the computation generalizes to a multivariate setting:

- If the included cell states are sufficiently different, their covariances become negligible, and the Mahalanobis distance behaves like a Euclidean norm over independent z-scores.
- If cell states are highly similar (i.e., strongly correlated), their individual contributions are weighted down, preventing redundant information from dominating the measure.

Thus, the Mahalanobis distance can be viewed as a multidimensional extension of a z-score, effectively quantifying the significance of function differences across conditions while accounting for the inherent covariance in our inferences, for example, the smoothening effect introduced by the GP-based posterior over gene expression.

#### 2.3 Implementation

A central challenge in computing the Mahalanobis distance is the inversion of the covariance matrix $\Sigma$. Direct inversion can be numerically unstable, especially when $\Sigma$ is nearly singular. To address this, we employ Cholesky decomposition:

$$\Sigma' = \Sigma + \epsilon \, \mathbb{I}, \tag{2.6}$$

where $\epsilon$ is a small regularization term ensuring numerical stability. The decomposition is then performed:

$$\Sigma' = L\, L^T, \tag{2.7}$$

with $L$ a lower triangular matrix. Instead of computing $\Sigma'^{-1}$ explicitly, we solve for $y$ via forward and backward substitution:

$$L\, y = x, \tag{2.8}$$

$$D(x) = \sqrt{y^T\, y}. \tag{2.9}$$

This method avoids direct matrix inversion, improving computational efficiency and stability.

#### 2.4 Applications

The covariance structure entering the Mahalanobis distance depends on the type of function being compared. For gene expression, which is conditioned directly on noisy observations, the relevant covariance is described in Section 1.3.1. For cell-state density, which is inferred through a latent representation rather than observed directly, the covariance instead incorporates the uncertainty of that representation (denoted $\sigma^2_{\text{post}}$; see Section 1.3.2). This distinction ensures that the uncertainty inherent in each type of function is properly accounted for in the comparison.

### Supplementary Note 3

### Computing $p$-values for Differential Expression

Differential expression analysis in *Kompot* is based on the Mahalanobis distance (Eq. 2.5), which quantifies the deviation between conditions while accounting for the posterior covariance. To interpret these distances statistically, we consider both Bayesian posterior tail probabilities and empirical false discovery rate (FDR) control. This chapter describes the full procedure from posterior tail probabilities through gene-shuffling null construction, local FDR estimation.

#### 3.1 Posterior Tail Probability

From a Bayesian perspective, the Mahalanobis distance corresponds to the radius of an isocline of a multivariate normal posterior distribution. Geometrically, if

$$x \sim \mathcal{N}(0, I_k),$$

then the squared Mahalanobis distance is

$$D^2(x) = x^T x,$$

which follows a $\chi^2$ distribution with $k$ degrees of freedom, where $k$ is the dimensionality of the evaluation space (i.e., the number of cell states, or the number of landmark cell states in the sparse case). Consequently, the Mahalanobis distance itself follows a $\chi$ distribution:

$$D(x) \sim \chi_k.$$

The posterior tail probability (PTP) of an observed distance $d$ is thus

$$\mathrm{PTP}(d) = \mathbb{P}[D \geq d] = 1 - F_{\chi_k}(d), \tag{3.1}$$

where $F_{\chi_k}$ is the cumulative distribution function of the $\chi$ distribution with $k$ degrees of freedom. Intuitively, this probability quantifies the chance of observing a deviation at least as large as $d$ under the posterior.

#### 3.2 Limitations in High Dimensions

In practice, when $k$ is large, posterior tail probabilities tend to be astronomically small. This is because the $\chi_k$ distribution concentrates its mass around $\sqrt{k}$, while deviations scale quickly in high dimensions. Moreover, PTPs do not reflect model-selection uncertainty or numerical approximations inherent in large-scale Gaussian process inference. Therefore, PTPs systematically overstate significance in practical high-dimensional settings.

#### 3.3 Empirical Null Distribution via Gene Shuffling

To address these limitations, *Kompot* employs an empirical strategy: we construct a null distribution of Mahalanobis distances by randomly shuffling gene identities (default: $N_{\text{null}} = 2{,}000$ shufflings). Specifically:

1. $N_{\text{null}}$ gene indices are sampled uniformly at random (without replacement)[1] from the set of all genes.

2. For each sampled null gene $j$, the expression vector $\mathbf{e}_j = (e_{j,1}, \ldots, e_{j,n_1+n_2})$ (combining cells from both conditions) is randomly permuted using a gene-specific random seed. This breaks the association between cell state and gene expression while preserving the marginal expression distribution.

3. The permuted expression vectors are split back into condition-specific matrices: $\tilde{\mathbf{e}}_j^{(1)}$ (condition 1, $n_1$ cells) and $\tilde{\mathbf{e}}_j^{(2)}$ (condition 2, $n_2$ cells).

4. Kompot's GP expression pipeline is applied to the shuffled genes: Gaussian process function estimation, posterior mean and covariance computation, and Mahalanobis distance calculation. This yields a set of null Mahalanobis distances $\{d_j^{\text{null}}\}_{j=1}^{N_{\text{null}}}$.

This procedure preserves the marginal distribution of each gene's expression values and the overall GP fitting procedure, so the null distances reflect the expected distribution of Mahalanobis distances when there is no true condition–expression association.

[1] When the requested $N_{\text{null}}$ exceeds the number of available genes (uncommon at the default $N_{\text{null}} = 2{,}000$, but possible for small antibody panels), Kompot falls back to sampling with replacement and emits a warning.

#### 3.4 Empirical P-values

For each real gene $i$ with observed Mahalanobis distance $d_i$, an empirical right-tailed $p$-value is computed:

$$p_i = \frac{1}{N_{\text{null}}} \sum_{j=1}^{N_{\text{null}}} \mathbf{1}[d_j^{\text{null}} \geq d_i].$$

This is a right-tailed test because larger Mahalanobis distances indicate greater deviation from the null hypothesis of no differential expression. In practice, the computation is vectorized using sorted null distances and binary search for efficiency.

**Handling zero p-values.** When the observed Mahalanobis distance exceeds all null distances, the empirical $p$-value is exactly zero. To avoid numerical issues in downstream calculations, Kompot replaces zero $p$-values with the minimum non-zero $p$-value observed across all genes. If all $p$-values are zero, the minimum is set to $1/N_{\text{null}}$.

#### 3.5 False Discovery Rate Control

From the empirical $p$-values, Kompot controls for multiple testing using two complementary approaches.

##### 3.5.1 Local FDR via Monotone Density Estimation

Local FDR estimation follows the framework of Efron [4, 5], adapted to work directly on Mahalanobis distances rather than $z$-scores. The key insight is that Mahalanobis distances are non-negative and their density under the null hypothesis is monotonically declining: small distances (no deviation from null) are common, while large distances (strong differential expression) are rare. Similarly, the mixture density of all genes (including both null and differentially expressed genes) must also be monotonically declining.

Kompot exploits this structural constraint using the Grenander estimator [6][2], which enforces monotonically declining density estimates via isotonic regression. The procedure consists of the following steps:

**Step 1: Boundary-corrected kernel density estimation.** For both the null Mahalanobis distances $\{d_j^{\text{null}}\}$ and the real distances $\{d_i^{\text{real}}\}$, an initial density estimate is obtained using a Gaussian kernel density estimator (KDE) with boundary correction at zero. The non-negative support of Mahalanobis distances requires special handling to prevent density leakage below zero. We use the *reflection method*: the data are mirrored around the origin to create a symmetric augmented dataset $\{-d_1, \ldots, -d_n, d_1, \ldots, d_n\}$, and the KDE is fitted to this augmented set. The density estimate on the non-negative half is then scaled by a factor of 2 to recover a proper density:

$$\hat{f}_{\text{KDE}}(x) = \frac{2}{nh} \sum_{i=1}^{n} K\left(\frac{x - d_i}{h}\right) + K\left(\frac{x + d_i}{h}\right),$$

where $K$ is the Gaussian kernel and $h$ is the bandwidth selected by Silverman's rule.

**Step 2: Grenander estimator via PAVA.** The initial KDE estimate is projected onto the space of monotonically non-increasing functions using the Pool Adjacent Violators Algorithm (PAVA). PAVA computes the monotone non-increasing sequence $\hat{f}_{\text{mono}}$ closest to $\hat{f}_{\text{KDE}}$ in weighted $L^2$ norm. This is the Grenander estimator of a monotone density [6]. The implementation follows the numerically stable incremental update from `fdrtool` [7] to avoid catastrophic cancellation when merging blocks with similar values. The result is normalized to integrate to 1 over the evaluation grid.

**Step 3: Local FDR computation.** The local FDR for gene $i$ is computed as the ratio of the monotone null density to the monotone mixture density at its observed Mahalanobis distance:

$$\text{lfdr}(d_i) = \frac{\hat{f}_0(d_i)}{\hat{f}(d_i)},$$

where $\hat{f}_0$ is the Grenander-estimated null density and $\hat{f}$ is the Grenander-estimated mixture density. Values are clipped to $[0, 1]$.

**Step 4: Monotonicity of local FDR.** The raw local FDR ratio is subjected to a final PAVA pass to enforce monotonically non-increasing behavior with respect to Mahalanobis distance. This guarantees that genes with larger distances (stronger differential-expression signal) always have lower local FDR than genes with smaller distances, a natural and desirable property.

**Interpolation.** Both density estimation and local FDR computation are performed on a regular grid of 500 points spanning the range of observed distances. Final local FDR values at each gene's observed distance are obtained by linear interpolation. Distances beyond the grid receive fill values of 1.0 (below grid, no signal) or 0.0 (above grid, strong signal).

##### 3.5.2 Significance Calling

By default, a gene is called significantly differentially expressed if its local FDR is below 0.05.

[2]Strictly, the implementation is an isotonic projection of a KDE estimate via the Pool Adjacent Violators Algorithm (PAVA), not the classical Grenander MLE (least-concave-majorant of the empirical CDF). The numerically stable PAVA update and the local- and tail-FDR conventions used here follow the `fdrtool` R package [7].

#### 3.6 Summary

In summary:

1. Posterior tail probabilities (PTPs) provide a direct Bayesian significance measure but are often unstable in high dimensions.

2. Empirical null distributions obtained via gene shuffling provide robust $p$-values and FDR estimates.

3. Local FDR is estimated using the Grenander estimator, which enforces monotonically declining density and local FDR with respect to Mahalanobis distance.

This hybrid approach combines Bayesian modeling with empirical error control, ensuring both sensitivity and robustness in high-dimensional differential expression analyses.

### Supplementary Note 4

### Semi-synthetic Benchmark for Differential Expression

To evaluate the sensitivity and specificity of Kompot's differential expression analysis and compare it against alternative approaches, we designed a semi-synthetic spike-in benchmark with known ground truth. A known log fold change pattern is introduced into a fixed set of genes in the young subset of the aging haematopoietic dataset, and each method is scored on how well it ranks those genes above the rest.

#### 4.1 Semi-synthetic Design

Standard benchmarking of differential expression methods in single-cell data is challenging because ground truth is rarely available in real biological datasets. We devised a semi-synthetic approach that preserves realistic data structure while introducing controlled, known expression changes.

Starting from a real single-cell dataset (Young mice only from the aging hematopoiesis dataset ($n$ = 2,917 cells, 16,285 genes), we constructed pseudo-conditions as follows:

1. **Duplicate:** All Young cells are duplicated to create two identical pseudo-conditions, condA (original) and condB (to be modified).
2. **Define target cells (per scenario):** Each scenario defines which cells change and by how much for each gene. *Celltype* changes one annotated cell type and leaves all other cells untouched. *Celltype-var* changes one merged group of B-cell subtypes in the same way. *Multi-celltype* assigns each gene its own target region: one diffusion component is drawn at random as an axis, a centre is drawn along it, and the cells within 25% of the coordinate range around that centre are split at the centre into two halves that receive opposite-signed changes. *Gradient* assigns every cell a change that varies smoothly with its position along the dominant diffusion component, so there is no boundary between changed and unchanged cells. *Uniform* changes every cell equally.
3. **Select spike-in genes:** For *Celltype* scenario, genes that clear a minimum mean count in the target cell type are drawn. For *Celltype-var*, genes whose mean expression varies most between the finer subtypes merged by its target group are drawn. For *Multi-celltype*, *Gradient* and *Uniform*, moderately expressed genes (25th–75th percentile of nonzero mean expression) are drawn as the ground-truth positive set. Every scenario is then truncated deterministically to exactly 500 spiked genes.
4. **Spike in:** The defined per-cell log fold changes are realized in the counts. Genes are perturbed by binomial thinning, in which each count is replaced by a binomial draw that retains a chosen fraction of its molecules. Thinning is applied to whichever of the two duplicated conditions has to fall, so a gene can be moved down in either direction; multiplying a single condition upwards could not represent a decrease at low counts. Non-integer results are discretised by stochastic rounding, which rounds up or down at random with probability set by the fractional part and is therefore unbiased in expectation. Per-scenario effect sizes are described in Section 4.2.
5. **Add cell-level noise:** To create a realistic background of gene-level variation and ensure a fair comparison across all methods, multiplicative log-normal noise is injected *independently* into both conditions at the raw-count level. Each gene $g$ is assigned a noise magnitude $\sigma_g \sim \text{Uniform}(0, \sigma_{\max})$ with $\sigma_{\max} = 0.3$ (in $\log_2$ units). For each cell $c$ and gene $g$, the noisy count is $x_{gc}^{\text{noisy}} = \text{round}(x_{gc} \cdot 2^{\epsilon_{gc}})$ where $\epsilon_{gc} \sim \mathcal{N}(0, \sigma_g)$, drawn independently for condA and condB. All methods receive the same noisy counts, so the noise characteristics are identical across arms.
6. **Combine:** The condA and condB cells are concatenated into a single object of 2 × 2,917 cells carrying a condition label, so that every method sees one dataset with two conditions rather than two datasets. Both copies keep the diffusion map coordinates of the cell they came from, so the embedding is never recomputed on the spiked data.
7. **Assign simulated mice:** To enable pseudobulk and mixed-model methods that require biological replicates, 8 simulated mice per condition (16 total) are created with unevenly distributed cell composition. Each mouse is assigned a random centre in diffusion map space, and 120 cells are sampled around it with probability proportional to a Gaussian kernel ($\sigma = 0.3$). Those 960 seeded cells are what fixes each mouse's position on the manifold; every remaining cell is then assigned to its nearest centre, so the mice partition the full condition and average roughly 365 cells each. This produces inter-mouse compositional variability analogous to real biological replicates, where different animals have different cell-type proportions.
8. **Add mouse-level noise:** To simulate realistic inter-replicate biological variability, a per-gene per-mouse systematic shift is applied after mouse assignment. Each mouse $m$ and gene $g$ receives a shift $\delta_{gm} \sim \mathcal{N}(0, 0.15)$ in $\log_2$ space, shared across all cells in that mouse. This creates between-replicate variance that does *not* cancel with pseudobulk averaging, preventing pseudobulk methods from obtaining artificially

clean dispersion estimates. Without this component, the benchmark would overstate the performance of pseudobulk methods relative to single-cell approaches, because the duplicated-cell design eliminates the inter-sample variability that is the primary noise source in real multi-sample experiments.

This design ensures that (i) the spike-in operates at the raw count level, respecting the discrete nature of scRNA-seq data, (ii) each method applies its own documented normalization, (iii) the underlying manifold structure is realistic because it derives from real biological data, and (iv) the random per-gene noise creates a challenging background against which methods must distinguish structured (manifold-coherent) from unstructured (random) expression changes.

**Normalization.** Each method normalizes the raw counts by its own documented default: Scanpy's `normalize_total` followed by a log transform, $\log_2(x + c) - \log_2(c)$ for Kompot, $\log_2(\mathrm{CPM} + 1)$ for MAST, a total-count offset for NEBULA, `librarySizeFactors` for LEMUR-unpaired, a natively inferred library size for scVI-DE, TMM for the edgeR arms and for miloDE, and median-of-ratios for the DESeq2 arms. Every method uses the same raw counts.

#### 4.2 Spike-in Scenarios

Every scenario uses **500** spike-in genes (**3.07%** of the 16,285 tested genes), leaving 15,785 negatives. AUPRC is computed directly against that full null. Per-scenario effect sizes are defined such that the five scenarios are comparably hard, judged by how well the injected change can be recovered from the ground truth itself. These choices were made without assessing performance on any of the methods under evaluation to ensure the effect sizes are not tuned to favor one method over another.

***Celltype*:** Each of the 500 spike-in genes draws an independent log fold change from $\mathrm{Uniform}(0, 1.25)$, applied only to the cells of the most abundant estimable cell type (773 of 2,917 cells) and leaving every other cell unchanged. This is the classical setting a cluster-stratified test is designed for: the effect is confined to one cleanly separable population and carries a non-zero mean within it.

***Celltype-var*:** Every spike-in gene receives a single constant log fold change of $\pm 0.55$, applied identically to every cell of one coarse cell group and zero to every cell outside it, with the direction drawn per gene. The group is the aggregate B compartment (537 of 2,917 cells), which merges six finer subtypes forming a developmental continuum from pro-B through pre-B, immature and mature naive to memory B cells. The spiked genes are those whose mean expression varies most between those subtypes. The fold change is therefore uniform across the group, and what varies is the expression it sits on: the constant shift must be recovered on top of real developmental structure that the coarse grouping flattens. A method that treats that within-group spread as dispersion cannot separate biological variability from noise, whereas a per-cell model resolves the sub-states and measures the shift against them.

***Multi-celltype*:** Each of the 500 spike-in genes receives its own randomly chosen target region in the phenotypic landscape. A random diffusion component defines the axis, and a random centre together with the cells spanning 25% of the coordinate range around it defines the cells that show expression change. Opposing effects of $+0.75$ and $-0.75$ are applied to the two halves on either side of that centre. No two genes share a profile, so each gene's aggregate fold change cancels towards zero while its local change stays large.

***Gradient*:** The log fold change of each of the 500 spike-in genes is a sinusoid of the cell's position along the dominant diffusion component. The sinusoid completes four full oscillations from one end of the continuum to the other, so the change alternates between up and down four times along it, with a typical magnitude of 0.45 in $\log_2$ units. Genes are offset against one another in 24 evenly spaced steps, so that the peaks and troughs of different genes fall at different positions rather than all coinciding. The change is realised by binomial thinning (Section 4.1), which allows the down-swing of the sinusoid to be represented at low counts. Because the change reverses several times along the continuum, any method that first groups cells averages opposite-signed parts of the same gene together and recovers less of it.

***Uniform*:** All 500 spike-in genes receive a global shift across every cell. Each gene draws an independent log fold change from $\mathrm{Uniform}(0, 0.75)$, giving a spectrum of effect sizes from near-zero upwards. This is the classical setting in which the change is the same everywhere on the manifold.

A null scenario was additionally run with no spike-in, in which the two conditions are identical copies of the original data and differ only by the random per-gene noise.

#### 4.3 Comparison Methods

Ten arms were benchmarked: Kompot; the neighborhood-based miloDE; the latent-embedding regression method LEMUR; the cell-level models MAST, NEBULA and scVI-DE; DESeq2 and edgeR applied to cluster-stratified pseudobulk profiles; and DESeq2 and edgeR applied to whole-condition pseudobulk in the manner of bulk RNA-seq. Every arm that aggregates cells uses the 8 simulated mice per condition as its replicate unit, so all of them need to account for the same inter-replicate compositional and expression variability.

##### 4.3.1 Kompot

Kompot's differential expression analysis was run with default parameters: GP-based expression function estimation for each condition, 2,000 null gene shuffles for FDR estimation, and the Mahalanobis distance as the primary gene-level test statistic. Mahalanobis distance was used directly as the continuous score for AUROC and AUPRC computation.

##### 4.3.2 Bulk DESeq2 and edgeR

The bulk arms aggregate all cells of a simulated mouse into a single whole-condition pseudobulk profile. Raw counts are summed within each mouse, giving 8 samples per condition and 16 in total, and a two-group test is fitted with condition as the only covariate. DESeq2 [8] is run through the Python implementation PyDESeq2 [9] with a Wald test; edgeR [10] is run on the same 16 profiles with its negative binomial generalized linear model and a likelihood ratio test, so that the two differ only in the statistical framework. The gene-level score is $s_g = -\log_{10}(p_g)$.

##### 4.3.3 Pseudobulk DESeq2 and edgeR

To resolve change within a condition, closer to what Kompot does, the pseudobulk arms stratify by cluster before aggregating. Cells are partitioned by unsupervised Leiden clustering [11], and within each cluster raw counts are summed per simulated mouse, so a cluster contributes up to 8 pseudobulk samples per condition. Samples with fewer than 5 cells are dropped, and a cluster is tested only if at least 2 samples per condition survive. DESeq2 or edgeR is then fitted independently within each cluster, and the gene-level score is the strongest evidence over clusters, $s_g = -\log_{10}\big(\min_j p_{g,j}\big)$.

##### 4.3.4 miloDE

miloDE [12] is a neighborhood-based differential expression method that constructs overlapping cell neighborhoods from a $k$-nearest-neighbor graph and tests for differential expression within each neighborhood using a negative binomial generalized linear model. Unlike the pseudobulk approach above, miloDE's neighborhoods overlap, are locally adaptive in size, and the statistical testing is integrated with the neighborhood construction.

A $k$-nearest-neighbor graph ($k = 30$) was built on the diffusion map embedding, and neighborhoods were assigned with second-order connectivity and filtering of redundant neighborhoods. Differential expression was tested within each neighborhood with condition as the only covariate, using the simulated mouse as the replicate identifier, so that miloDE sees the same 8 replicates per condition as the pseudobulk arms.

**Choice of embedding.** miloDE was run on the same diffusion map as Kompot, so that both methods receive the same representation of cell state and any difference between them reflects the test rather than its input since miloDE leaves this choice to the user [12]. Running miloDE on the 50 Harmony-corrected principal components instead, with the counts, the design, the replicate assignment and every miloDE parameter held fixed, raises its AUPRC in all five scenarios, by 0.074 to 0.219 over 21 paired seed replicates. The reason is likely neighborhood size. At $k = 30$ the diffusion map yields 85 neighborhoods averaging 117 cells, against 43 averaging 302 on the principal components, where the authors recommend several hundred; our diffusion map is stored without eigenvalue weighting, so a Euclidean $k$-nearest-neighbor metric weights its fortieth component as heavily as its first. The ordering of the two methods is unchanged in four of the five scenarios. In *Celltype* it reverses: Kompot's AUPRC margin over miloDE goes from $+0.022$ to $-0.143$. We report the diffusion map run because it is the comparison the benchmark is built to make since holding the representation fixed isolates the test from its input, and giving the two methods different embeddings would confound the two. The principal-component result is reported here since it is the configuration in which miloDE performs best and it changes the performance in one scenario.

**Gene-level aggregation.** miloDE reports per-neighborhood, per-gene results, so a gene-level summary is needed for comparison with Kompot's gene-level Mahalanobis distance. The minimum $p$-value aggregation is used, $s_g = -\log_{10}\big(\min_j p_{g,j}\big)$, where a gene is ranked highly if it is significant in *any* neighborhood.

##### 4.3.5 MAST

MAST [13] models each gene with a two-part hurdle: a logistic component for whether the gene is detected in a cell and a Gaussian component for its expression level given detection. The two are fitted jointly on $\log_2(\text{CPM}+1)$ values at cell-level resolution, with condition as the covariate of interest and the cellular detection rate, the fraction of genes detected in each cell, as a nuisance covariate. Significance is a likelihood ratio test on the condition coefficient over both components, and the gene-level score is $s_g = -\log_{10}(p_g)$. MAST is fitted without a random effect for the simulated mouse, which is the configuration in standard use and the one most benchmarks evaluate; it therefore treats cells as independent observations and does not model the mouse-level variance component that the design injects.

##### 4.3.6 NEBULA

NEBULA [14] is a negative binomial mixed model designed for single-cell differential expression analysis that avoids the information loss of pseudobulk aggregation. Rather than collapsing cells into sample-level pseudobulk counts, NEBULA fits a negative binomial generalized linear mixed model directly to individual cell counts, with the sample identifier (here, simulated mouse) as a random intercept. This properly accounts for within-sample correlation, the key statistical challenge in multi-sample single-cell designs, without discarding cell-level resolution.

NEBULA was run using the fast H-likelihood (HL) approximation, which provides a computationally efficient alternative to full maximum likelihood for large cell counts. The model included condition (condA vs. condB) as the fixed effect and mouse identity as the random effect, with log library size as an offset term. NEBULA internally filters genes with very low expression; genes filtered out by NEBULA receive NA scores. The gene-level score is $s_g = -\log_{10}(p_g)$, where $p_g$ is the $p$-value for the condition coefficient.

##### 4.3.7 scVI-DE

scVI [15] fits a variational autoencoder with a negative binomial likelihood to raw counts and performs differential expression by sampling from the approximate posterior of the two conditions, returning for each gene a probability that its expression differs. We used the Bayes-factor-derived probability from the model's differential-expression routine as the score, which is direction-agnostic by construction.

In the configuration reported here, scVI infers each cell's library size from its total counts. Because the spike-in moves a substantial share of the count mass in one direction, the inferred library absorbs part of that shift, and the resulting compositional artifact is scored as if it were signal. This is a mismatch between the model's library treatment and a spike-in design, not a general property of scVI, and the reported values should not be read as its performance on real data.

##### 4.3.8 LEMUR

LEMUR [16] is a latent embedding multivariate regression framework that models gene expression as a function of both cell state and experimental condition in a shared latent space. Unlike pseudobulk methods, LEMUR operates directly on single cells and can detect condition-dependent expression changes that vary across the manifold. The LEMUR pipeline consists of three steps: (1) fitting a latent embedding model with `lemur()`, (2) computing per-cell predicted fold changes with `test_de()`, and (3) identifying differentially expressed neighborhoods with calibrated p-values via `find_de_neighborhoods()`, which performs pseudobulk aggregation within optimally selected cell neighborhoods.

In our benchmark, LEMUR was run with condition as the only covariate and the simulated mouse as the replicate unit for the neighborhood pseudobulk test.

**Note on LEMUR performance.** LEMUR reaches AUROC 0.62 to 0.71 across the five reported scenarios, above chance but far below the leading arms. We validated that this is not an implementation error by running LEMUR on its own example dataset (glioblastoma, Zhao et al. 2021), where it correctly identifies 82/300 genes at $p < 0.05$ in the paired design. The poor benchmark performance reflects a design mismatch: LEMUR was developed and validated on multi-condition datasets with natural biological variation between conditions [16]. Its vignette demonstrates a *paired* design where each patient contributes cells to both conditions (`design = ~ patient_id + condition`), and the manuscript states that "applications range from comparisons between two conditions with replicates, over paired studies [...] over studies with multiple covariates." In our semi-synthetic benchmark, both conditions are duplicates of the same cells with artificial spike-ins. The LEMUR model absorbs the condition effect into the latent embedding during fitting, leaving insufficient residual signal for the downstream pseudobulk test in `find_de_neighborhoods()`. This is a fundamental limitation of the benchmark's duplicated-cell design for LEMUR's specific modeling approach, not a deficiency of LEMUR itself. LEMUR's results are included for completeness but should not be interpreted as reflecting the method's performance on real biological data.

#### 4.4 Evaluation Metrics

For each scenario, the benchmark evaluates gene-level detection performance. Let $y_g \in \{0, 1\}$ indicate whether gene $g$ was truly spiked ($y_g = 1$) or not, and let $s_g$ be the continuous detection score assigned by each method.

**AUPRC** (Area Under the Precision-Recall Curve): The spiked set is 500 genes against all 15,785 non-spiked genes, so AUPRC is computed directly at the true 3.07% prevalence and its chance baseline is 0.0307. AUPRC is more informative than AUROC when the positive class is small, and is particularly sensitive to false positives among high-ranked genes. It is also, unlike AUROC, invariant to negatives appended below the lowest-ranked positive, a property that matters here because methods differ in how many genes they return a statistic for.

**AUROC** (Area Under the Receiver Operating Characteristic curve): Measures the probability that a randomly chosen spiked gene has a higher score than a randomly chosen non-spiked gene. AUROC = 0.5 corresponds to random ranking.

Methods that return no statistic for a gene have that gene assigned the lowest possible score rather than excluded, so all arms are scored over the same 16,285 genes. All methods receive the same noisy data: cell-level and mouse-level noise are applied at the raw-count level before normalization and log-transformation, ensuring that all methods face the same noise characteristics regardless of whether they operate on raw counts or log-transformed expression. Simulated mouse assignments and all metadata are precomputed in Python and exported for R-based methods, ensuring identical inputs across implementations.

#### 4.5 Results

Kompot and MiloDE are the two leading methods and are tabulated head-to-head below, with AUPRC and AUROC computed against the full null of 15,785 non-spiked genes at the true 3.07% prevalence. $\Delta$ is Kompot minus MiloDE; a negative value favors MiloDE. Intervals and $P$ values are from a paired gene-level bootstrap (2,000 resamples over all genes). That bootstrap resamples gene indices within a single simulated dataset, holding each method's fitted output fixed. It therefore quantifies how much a margin depends on which genes happen to be in the universe. It is *not* an estimate of variability across independent simulations, spike draws or replicate assignments, none of which we measured, and the intervals should not be read as method-level uncertainty.

| Scenario | AUPRC | | | AUROC | | |
|---|---|---|---|---|---|---|
| | Kompot | MiloDE | $\Delta$ | Kompot | MiloDE | $\Delta$ [95% CI], $P$ |
| *Celltype* | **0.348** | 0.326 | +0.022 | 0.854 | **0.890** | −0.036 [−0.051, −0.021], <0.001 |
| *Celltype-var* | **0.454** | 0.270 | +0.184 | 0.953 | 0.932 | +0.021 [+0.011, +0.031] |
| *Multi-celltype* | **0.590** | 0.463 | +0.127 | 0.930 | 0.928 | +0.002 [−0.009, +0.013], 0.71 (tie) |
| *Gradient* | **0.681** | 0.369 | +0.313 | **0.960** | 0.936 | +0.024 [+0.015, +0.033], <0.001 |
| *Uniform* | **0.649** | 0.338 | +0.311 | **0.930** | 0.910 | +0.020 [+0.009, +0.031], 0.003 |

Kompot leads AUPRC in all five, each value 11 to 22 times the 0.0307 chance baseline.Kompot leads AUROC in *Celltype-var*, *Gradient* and *Uniform*, *Multi-celltype* is a tie, and MiloDE leads *Celltype*. The two metrics therefore disagree in *Celltype* alone.

**Partial coverage and its effect on the metrics.** Methods differ in how many genes they return a statistic for, and our benchmarking assigns the lowest possible score to the genes where no statistic was assigned. MiloDE omits 3,117–3,436 genes, 19.1–21.1% of the 16,285, across *Celltype*, *Multi-celltype*, *Gradient* and *Uniform*, the four scenarios for which both methods' full gene-level output was retained. Those omissions are low-expression genes, so none of them could have been spiked. Each is therefore scored as a correct rejection at no cost, while Kompot returns a statistic for the same genes and can rank one above a spiked gene. This favors MiloDE, and applies to any method that filters on expression. The choice of gene exclusion is made by the user when applying Kompot.

The resulting effect however is small and is outweighed by a second one acting in the opposite direction. Kompot reaches AUROC 0.980–0.996 on the omitted block against MiloDE's guaranteed 1.000, so the two are nearly tied there. Overall AUROC averages over all positive–negative gene pairs, so including a near-tied block compresses every margin towards zero. Restricting the comparison to genes both methods score removes that compression and widens each margin: Kompot's lead rises from +0.020 to +0.028 in *Uniform* and from +0.024 to +0.032 in *Gradient*, and MiloDE's lead in *Celltype* rises from 0.036 to 0.041 ($P < 0.001$).

**Effect of replicate composition.** Giving each simulated mouse a different cell composition is what keeps the benchmark realistic. The alternative is to build mice by partitioning cells at random, which gives every mouse the same mixture of cell types. In that case the pseudobulk profiles of the replicates are nearly identical, their estimated dispersion is far smaller than any real experiment would give, and the bulk and mixed-model arms gain power they would not have on real data. Real animals differ in their cell-type proportions, so we assign cells by position on the manifold instead, and the resulting between-replicate variability is what these methods would actually face.

### Supplementary Note 5

### Limitations

Kompot has three main categories of limitations. The first concerns uncertainty arising from model selection and the assumptions inherent to the Gaussian Process formulation. The second concerns imperfections in the latent cell-state representation provided as input. The third concerns distributional assumptions for gene expression and the treatment of overdispersion.

#### 5.1 Model Selection and Statistical Uncertainty

As with any Bayesian inference procedure, Kompot cannot fully account for uncertainty due to model selection. This is a well-known limitation across Bayesian modeling [17, 18]. Because Kompot uses a non-parametric Gaussian Process to model arbitrary smooth functions over the manifold, most sources of model misspecification are absorbed into the GP itself [1]. The remaining unmodeled uncertainty is therefore largely confined to the choice of covariance kernel (Section 1.1.1) and its hyperparameters. We use the Matérn $\frac{5}{2}$ kernel, which is the least smooth option among twice-differentiable kernels. This provides sufficient smoothness for potential applications such as gradient-based optimization algorithms on the phenotypic manifold, while remaining minimally restrictive and avoiding the infinite smoothness of the squared exponential kernel. It avoids over-smoothing and therefore does not systematically underestimate posterior uncertainty [1, 2]. Some uncertainty remains with respect to the kernel length-scale. Our stability analysis shows that Kompot produces highly consistent results across a wide range of length-scales, suggesting that the contribution of this uncertainty is small.

For differential expression, Kompot compares the posterior distributions of gene expression functions between conditions using the Mahalanobis distance (Eq. 2.5). The distance is a practical and interpretable measure, but we view it as an ad hoc summary rather than a direct posterior probability. Although Kompot can supply a posterior tail probability for each distance, we do not recommend using these values for significance testing. In practice, we observe extremely small posterior tail probabilities, which we attribute to numerical issues and the very high dimensionality of the evaluation space. In high dimensions, the integral over radii of the $n$-sphere may become unreliable for practical inference. Instead, we generate an empirical null distribution of Mahalanobis distances and determine local false discovery rates for each dataset (Supplementary Note 3). Using a default local false discovery rate of 0.05 yields conservative and well-calibrated estimates of differential expression.

For differential abundance, posterior uncertainty over the latent parameters comes from a Laplace approximation at the MAP estimate (Section 1.2). Only the diagonal of the Hessian is used, so residual correlations between parameters are not represented. The GP parameterization we adopt (in which the latent parameters are defined as $\mathbf{z} \sim \mathcal{N}(\mathbf{0}, \mathbf{I})$ and the function values recovered via $\mathbf{f} = \mathbf{L}\mathbf{z} + \mu$) removes the dominant source of parameter correlation, namely the prior covariance from cell-to-cell similarity, so the neglected terms are expected to be small.

#### 5.2 Latent Representation of Cell State

The second class of limitations arises from the latent cell-state representation used as input to Kompot. Although Kompot does not prescribe how this representation is constructed, its performance depends on the quality of the cell-state distinction and cell-to-cell distance in that space. We typically recommend diffusion maps, but other embeddings are possible. We do not prescribe a specific dimensionality or level of resolution. These choices are similar to decisions commonly made when selecting the number of principal components or resolution parameters in other single-cell workflows.

Suboptimal embeddings reduce sensitivity but do not lead to incorrect conclusions. The poorest possible representation collapses all cells into a single point. In this setting, Kompot's differential abundance reduces to detecting no changes, and differential expression becomes equivalent to a standard pseudobulk comparison. Too little resolution brings the analysis closer to pseudobulk methods, while excessive resolution introduces noise and makes information sharing between nearby states difficult. Both effects reduce sensitivity without generating spurious results. In principle, one could construct a cell-state embedding that intentionally falsifies results by placing cells that are similar within one condition close to cells from the other condition that have very different expression profiles. However, such adversarial embeddings represent a form of severe misalignment, rather than a typical failure mode of dimensionality reduction.

A more substantial limitation arises from uncertainty in the cell's location in the latent space and from potential subtle misalignment between the two conditions. Kompot does not explicitly model the uncertainty of individual cell positions. It does, however, implicitly model uncertainty in cell-to-cell similarity through the GP prior covariance kernel $k(x, x')$ (Section 1.1.1), which encodes how function values should relate based on distance in the latent space. Still, when embeddings place cells from different conditions in different positions, Kompot will measure differences at corresponding coordinates in latent space. Whether two truly "equivalent" regions are correctly aligned depends on the chosen embedding, the level of dimensional reduction, and any applied batch correction.

In many datasets, perturbations produce subtle changes that are well aligned across conditions, making misalignment negligible. However, when states shift strongly between conditions, embeddings may place equivalent cell types at distinct coordinates. Kompot will then detect broad differential expression that reflects intrinsic differences between the misaligned states rather than condition-specific changes. Likewise, differential abundance may appear as strong gains and losses in neighboring regions that lack any true counterpart in the other condition (Supp. Fig. 14C). These patterns are diagnostic: pronounced but opposing differential abundance in adjacent states or differential expression dominated by genes with similar structure but shifted expression patterns in the two conditions can indicate misalignment rather than biological effect.

This limitation is not unique to Kompot, but inherent to any method that attempts to compare conditions for matched cell types. Differential analysis across conditions always assumes that corresponding states are selected for comparison; if they are not, misalignment can be interpreted as biological change. Cluster-based approaches only avoid this when comparisons remain coarse, for example when entirely new cell types appear that are compared to the rest. For analyses that aim to compare conditions at the level of specific cell states, whether continuously or discretely defined, the main remaining challenge is therefore the same as in **batch correction**: establishing reliable correspondence between the phenotypic landscapes of the two conditions.

#### 5.3 Distributional Assumptions for Gene Expression

Kompot's differential expression analysis relies on the assumption that log-transformed single-cell expression values are approximately normally distributed. This assumption enables the closed-form Gaussian Process posterior (Section 1.3.1) and is reflected in the measurement noise model (Section 1.2.2).

In the default mode, the observation noise variance is treated as constant across the manifold. This does not account for overdispersion, the well-documented phenomenon in RNA-seq data where variance scales with the mean [19].

However, empirical variance is disabled by default. The variance estimated from residuals reflects the probability of making different measurements at a given cell state, not the uncertainty about real biological differences. In our experiments, enabling empirical variance had little impact on gene rankings by Mahalanobis distance, and therefore little impact on FDRs and overall results. Fully feeding back the empirical variance to update the mean inference and posterior uncertainty would require a full Cholesky decomposition for each gene (making the per-gene cost comparable to explicit sample variance computation) and would constitute double-dipping, using the same residuals both to estimate variance and to update the posterior that produced them. We therefore do not recommend this approach and keep empirical variance off by default. The empirical-variance option is nonetheless implemented in Kompot, and demonstrates that a gene-specific, cell-state-dependent variance can be incorporated within the GP framework at minimal additional computational cost. It can be enabled to account for overdispersion-based uncertainty in datasets where heteroscedastic noise is expected to substantially affect gene rankings.

In principle, one could replace the Gaussian framework entirely with exact negative-binomial likelihoods. However, this would eliminate the closed-form posterior and require separate likelihood-based inference for each gene, increasing computational cost by several orders of magnitude.

Accounting for inter-replicate biological variability, or for gene-specific and cell-state-specific observation variance, would require a richer noise model than the constant-variance assumption used here, at substantially greater computational cost.
